# Systems-wide Analysis Revealed Shared and Unique Responses to Moderate and Acute High Temperatures in the Green Alga *Chlamydomonas reinhardtii*

**DOI:** 10.1101/2021.08.17.456552

**Authors:** Ningning Zhang, Erin M. Mattoon, Will McHargue, Benedikt Venn, David Zimmer, Kresti Pecani, Jooyeon Jeong, Cheyenne M. Anderson, Chen Chen, Jeffrey C. Berry, Ming Xia, Shin-Cheng Tzeng, Eric Becker, Leila Pazouki, Bradley Evans, Fred Cross, Jianlin Cheng, Kirk J. Czymmek, Michael Schroda, Timo Mühlhaus, Ru Zhang

**Affiliations:** Donald Danforth Plant Science Center, St. Louis, Missouri 63132, USA; Plant and Microbial Biosciences Program, Division of Biology and Biomedical Sciences, Washington University in Saint Louis, St. Louis, MO 63130, USA; TU Kaiserslautern, Kaiserslautern 67663, Germany; The Rockefeller University, New York, New York 10065, USA; University of Missouri-Columbia, Columbia, Missouri 65211, USA

## Abstract

Different intensities of high temperatures affect the growth of photosynthetic cells in nature. To elucidate the underlying mechanisms, we cultivated the unicellular green alga *Chlamydomonas reinhardtii* under highly controlled photobioreactor conditions and revealed systems-wide shared and unique responses to 24-hour moderate (35°C) and acute (40°C) high temperatures and subsequent recovery at 25°C. We identified previously overlooked unique elements in response to moderate high temperature. Heat at 35°C transiently arrested the cell cycle followed by partial synchronization, up-regulated transcripts/proteins involved in gluconeogenesis/glyoxylate-cycle for carbon uptake, promoted growth, and increased starch accumulation. Heat at 40°C arrested the cell cycle, inhibited growth, resulting in carbon uptake over usage and increased starch accumulation. Both high temperatures induced photoprotection, while 40°C decreased photosynthetic efficiency, distorted thylakoid/pyrenoid ultrastructure, and affected the carbon concentrating mechanism. We demonstrated increased transcript/protein correlation during both heat treatments, suggesting reduced post-transcriptional regulation during heat may help coordinate heat tolerance activities efficiently. During recovery after both heat treatments, transcripts/proteins related to DNA synthesis increased while those involved in photosynthetic light reactions decreased. We propose down-regulating photosynthetic light reactions during DNA replication benefits cell cycle resumption by reducing ROS production. Our results provide potential targets to increase thermotolerance in algae and crops.

## Introduction

High temperatures occur frequently in nature and impair crop yields and algal biofuel production (Zhao et al., 2017; Lobell et al., 2011; Barnabás et al., 2008; Siebert et al., 2014; Iba, 2002). Global warming increases the intensity, duration, and frequency of high temperatures above the optimal range for plant growth. It is projected that for every degree Celsius mean global temperature increase, the yields of major crop species will decrease by 3% ∼ 8% (Zhao et al., 2017; Lobell et al., 2011). Photosynthetic organisms experience different intensities of high temperatures in field conditions. Many crop species have threshold temperatures between 25 and 40°C, above which reduced growth is observed; most heat stress experiments have been conducted at acutely high temperatures near 42°C or above (Janni et al., 2020). Plants frequently experience sustained moderate high temperatures around 35°C in nature, however these conditions have been largely understudied (Sharkey, 2005). Acute high temperature at or above 40°C is more damaging but usually less frequent or shorter-lasting than moderate high temperature in the field. We hypothesize that plants can acclimate to moderate high temperature but may have less capacity to acclimate to acute high temperature. Additionally, we propose that different levels of high temperatures induce shared and unique responses in photosynthetic cells. Understanding how photosynthetic cells respond to and recover from different intensities of high temperatures is imperative for improving crop thermotolerance (Mittler et al., 2012).

High temperatures are known to have a wide variety of impacts on photosynthetic cells. Heat-increased membrane fluidity has been proposed to activate membrane-localized mechanosensitive ion channels leading to increased intracellular calcium concentrations, which may cause signaling cascades to activate heat shock transcription factors (HSFs) (Saidi et al., 2009; Wu et al., 2012; Königshofer et al., 2008). HSFs act in the nucleus to increase transcription of genes involved in heat response, e.g., heat shock proteins (HSPs) (Schroda et al., 2015; Schulz-Raffelt et al., 2007; Schmollinger et al., 2010). A recent work proposed that the accumulating cytosolic unfolded proteins, rather than changes in membrane fluidity, trigger the expression of HSPs in green algae (Rütgers et al., 2017b). Furthermore, high temperatures can decrease the stability of RNAs and alter the transcriptomic landscape of cells under heat stress (Su et al., 2018). Additionally, high temperature at 40°C can cause damage to photosynthetic electron transport chains, reducing photosynthetic efficiency (Zhang and Sharkey, 2009; Sharkey and Zhang, 2010), and leading to increased reactive oxygen species (ROS) accumulation (Janni et al., 2020). Heat-induced ROS production increases DNA damage and the need for DNA repair pathways, although the mechanisms of these processes are poorly understood (Schroda et al., 2015; Velichko et al., 2012). In contrast to the extensive research on the effects during heat, how photosynthetic cells recover from heat is much less studied.

Algae have great potential for biofuel production and bioproduct accumulation, but the knowledge surrounding mechanisms of algal heat responses are largely limited as compared to land plants (Schroda et al., 2015). Outdoor algal ponds frequently experience supra-optimal temperatures at or above 35°C during summer time (Mata et al., 2010), but how algal cells respond to moderate high temperatures remains largely understudied. Many previous algal heat experiments were conducted in flasks incubated in hot water baths (at or above 42°C) with sharp temperature switches, e.g., by resuspending centrifuged cells in pre-warmed medium to initiate the heat treatments (Hemme et al., 2014; Rütgers et al., 2017b). The previous research was valuable for paving the road to understand algal heat responses. Nevertheless, high temperatures in nature, especially in aquatic environments, often increase gradually and the rate of temperature increase may affect heat responses (Mittler et al., 2012). Acute high temperature at 42°C results in algal cell cycle arrest (Mühlhaus et al., 2011; Hemme et al., 2014). Long-term experiments at moderate high temperatures that do not lead to a sustained cell cycle arrest cannot be conducted in flasks because cultures grow into stationary phase, causing nutrient and light limitation and therefore complicating analyses. Consequently, investigating algal heat responses under well-controlled conditions in photobioreactors (PBRs) with turbidostatic modes can mimic the heating speed in nature and reduce compounding factors unrelated with high temperatures during the experiment, improving our understanding of algal heat responses.

The unicellular green alga, *Chlamydomonas reinhardtii* (Chlamydomonas throughout), is an excellent model to study heat responses in photosynthetic cells for many reasons, including its fully sequenced haploid genome, unicellular nature allowing for homogenous treatments, generally smaller gene families than land plants, and extensive genetic resources (Jinkerson and Jonikas, 2015; Crozet et al., 2018; Zhang et al., 2014; Li et al., 2016, 2019; Dhokane et al., 2020). At the cellular level, Chlamydomonas has many similarities with land plants, making it a promising model organism to identify novel elements with putative roles in heat tolerance with implications for crops (Schroda et al., 2015).

A previous transcriptome and lipidome-level analysis in Chlamydomonas under acute high temperature (42°C) over 1 hour (h) revealed changes in lipid metabolism and increased lipid saturation as one of the early heat responses (Légeret et al., 2016). Additionally, a proteome and metabolome-level analysis of acute high temperature (42°C) for 24-h followed by 8-h recovery demonstrated temporally resolved changes in proteins, metabolites, lipids, and cytological parameters in Chlamydomonas (Hemme et al., 2014). Both publications contributed to the foundational knowledge of how Chlamydomonas responds to acute high temperature. However, a temporally resolved transcriptome analysis during and after heat over a relatively long time is lacking and the correlation between transcriptome, proteome, and physiological responses to different intensities of high temperatures remains elusive.

We investigated the response of wild-type Chlamydomonas cells to moderate (35°C) or acute (40°C) high temperatures at transcriptomic, proteomic, cytological, photosynthetic, and ultrastructural levels over a 24-h heat followed by 48-h recovery period in PBRs under well-controlled conditions. Our results showed that some of the responses were shared between the two treatments and the effects of 40°C were typically more extensive than 35°C; however, 35°C induced a unique set of responses that were absent under 40°C. Both 35 and 40°C induced starch accumulation but due to distinct mechanisms. We showed that 35°C transiently inhibited cell cycle followed by synchronization while 40°C halted the cell cycle completely. Heat at 40 but not 35°C reduced photosynthetic efficiency, increased ROS production, and altered chloroplast structures. Furthermore, with the time-resolved paired transcriptome and proteome dataset, we demonstrated increased transcript and protein correlation during high temperature which was reduced during the recovery period. Additionally, we revealed up-regulation of genes/proteins related to DNA replication and down-regulation of those related to photosynthetic light reactions during early recovery after both heat treatments, suggesting potential crosstalk between these two pathways when resuming the cell cycle. These data further our understanding of algal responses to high temperatures and provide novel insights to improve thermotolerance of algae and crop species.

## Results

### Heat at 35°C increased algal growth while 40°C largely inhibited it

The growth rate of Chlamydomonas cells in PBRs grown mixotrophically under well-controlled conditions (light, temperature, air flow, cell density, and nutrient availability) increased with temperature and plateaued at 35°C, but was largely inhibited at 40°C, as compared to the control 25°C (Figure 1A). Hence, we defined 35°C as moderate high temperature and 40°C as acute high temperature under our experimental conditions. To investigate the systems-wide responses of Chlamydomonas to moderate and acute high temperatures, we grew algal cells in PBRs with well-controlled conditions at constant 25°C first, followed by 24-h heat treatment at 35 or 40°C, and recovery at 25°C for 48 h (Figure 1B, Supplemental Figure 1A-C). Temperature increase from 25 to 35 or 40°C took about 30 minutes (min), and neither heat treatment affected cell viability (Supplemental Figure 1D, E). In contrast, we found that a sharp temperature switch without gradual temperature increase reduced cell viability (Supplemental Figure 1F), suggesting heating speed affects thermotolerance. The transcript levels of selected circadian regulated genes, *LHCA1* (Hwang and Herrin, 1994) and *TRXF2* (Barajas-López et al., 2011), did not change under constant 25°C (Supplemental Figure 1G, H), suggesting minimal circadian regulation existed under our experimental conditions with constant light and the observed changes during and after heat were therefore most likely attributable to heat treatments.

**Figure 1.**
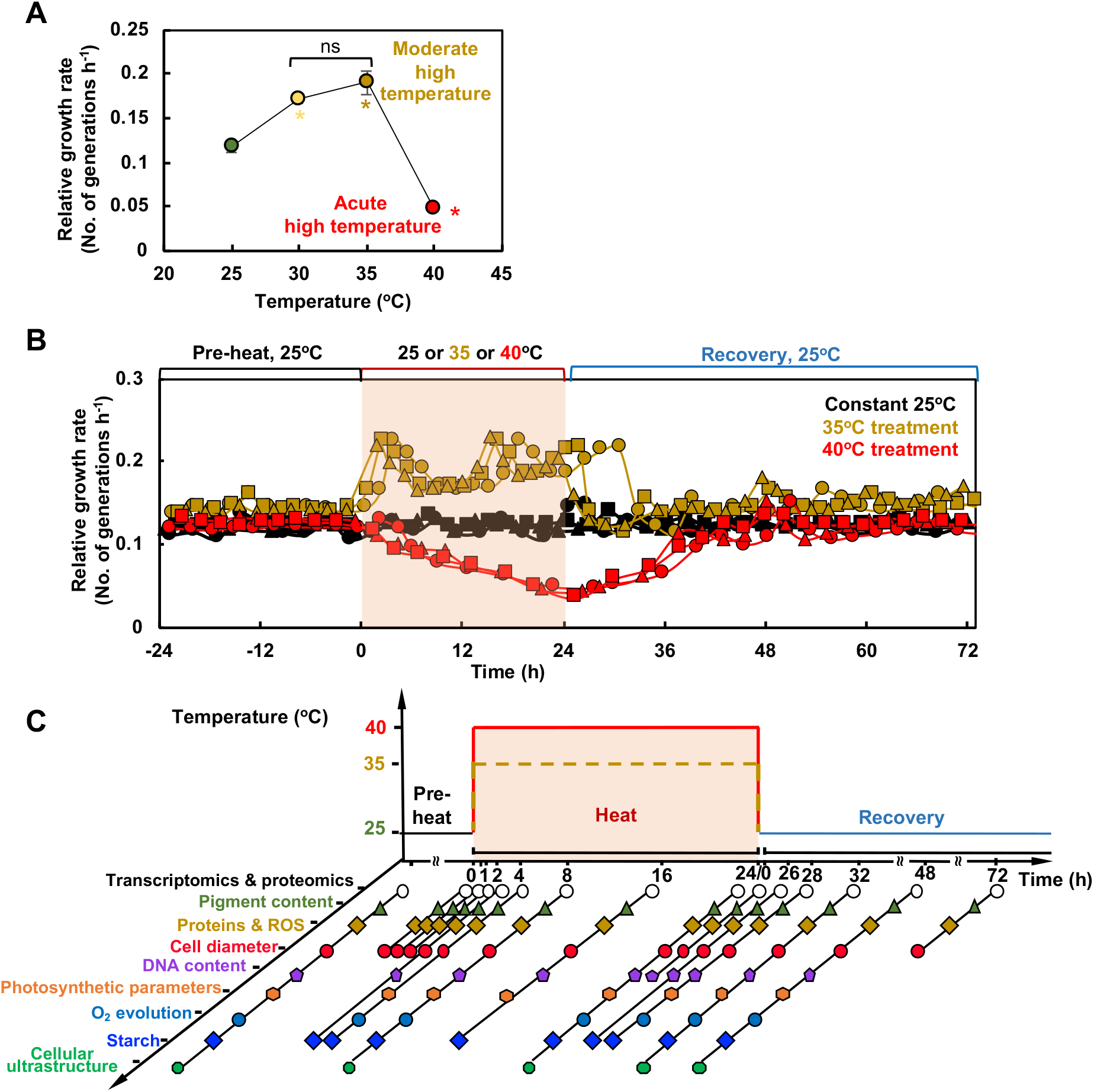
Moderate (35°C) and acute (40°C) high temperatures had contrasting effects on the growth rates of Chlamydomonas cells. **(A)** Chlamydomonas growth rate plateaued around 35°C but was largely inhibited at 40°C. Chlamydomonas cells (CC1690, 21gr, wildtype) were grown in photobioreactors (PBRs) in Tris-acetate-phosphate (TAP) medium under turbidostatic conditions at different temperatures with a light intensity of 100 µmol photons m^−2^ s^−1^ and constantly bubbling of air. Growth rates were calculated based on the cycling of OD_680_ at the end of 2-day treatment of each temperature, see Supplemental Figure 1 and methods for details. Each temperature treatment was conducted in an individual PBR. Mean ± SE, *n*=3 biological replicates. Statistical analyses were performed using a two-tailed t-test assuming unequal variance by comparing treated samples with pre-heat (*, p<0.05, the colors of asterisks match the treatment conditions). Not significant, ns. **(B)** Heat treatment at 35°C (brown) increased growth rates while 40°C (red) inhibited it. Algal cultures in separate PBRs were first acclimated at 25°C for 4 days before the temperature was switched to 35 or 40°C for 24 h, followed by recovery at 25°C for 48 h. Algal cultures grown constantly at 25°C (black) served as controls, which demonstrated steady growth without heat. Three independent biological replicates for each condition were plotted. **(C)** PBR cultures with different treatments were sampled at a series of time points to study heat responses at multiple levels. Symbol colors match the colors of the parameters assayed. (**B**, **C**) The red shaded areas depict the duration of high temperature.

Algal growth increased during 35°C, decreased during 40°C, and remained steady at constant 25°C (Figure 1B). Increased growth under 35°C was confirmed by the medium consumption rates and growth on plates (Supplemental Figure 2). We harvested the PBR cultures throughout the time-course experiment for systems-wide analyses, including transcriptomics, proteomics, cell physiology, photosynthetic parameters, and cellular ultrastructure (Figure 1C). RT-qPCR analysis of select time points showed that both 35 and 40°C induced heat stress marker genes (*HSP22A* and *HSP90A*) and *HSF*s, although the induction amplitude was much larger under 40°C than 35°C (up to 20x, Supplemental Figure 3A-D), suggesting both heat treatments could induce heat responses. RNA-seq data were verified by testing select genes with RT-qPCR, with high levels of consistency between the two methods (Supplemental Figure 3).

### Transcriptomic and proteomic analyses identified shared and unique responses during and after 35 and 40°C treatments

Two-dimensional Uniform Manifold Approximation and Projection (UMAP) of Transcripts per Million (TPM) normalized RNA-seq data resulted in three distinct clusters (Figure 2A). The cluster during heat had a pattern displaying temporally resolved transcriptomes between 35 and 40°C treated samples, showing increasing variance between 35 and 40°C time points throughout the high temperature treatment. The early recovery cluster consisted of 2- and 4-h recovery samples after 35°C heat, as well as 2-, 4-, and 8-h recovery samples after 40°C heat. Late recovery samples clustered with pre-heat samples, suggesting fully recovered transcriptomes by 8-h following 35°C and 24 h following 40°C treatment, respectively. UMAP of proteomics data results in two distinct clusters, separating the 35 and 40°C treated samples and demonstrating temporally resolved proteomes (Figure 2B). However, the samples during heat, early, and late recovery did not fall into their own distinct clusters, indicating resistance to rapid changes on the protein level as compared with the transcript level. The proteome recovered to pre-heat levels by the 48-h recovery after 35 and 40°C.

**Figure 2:**
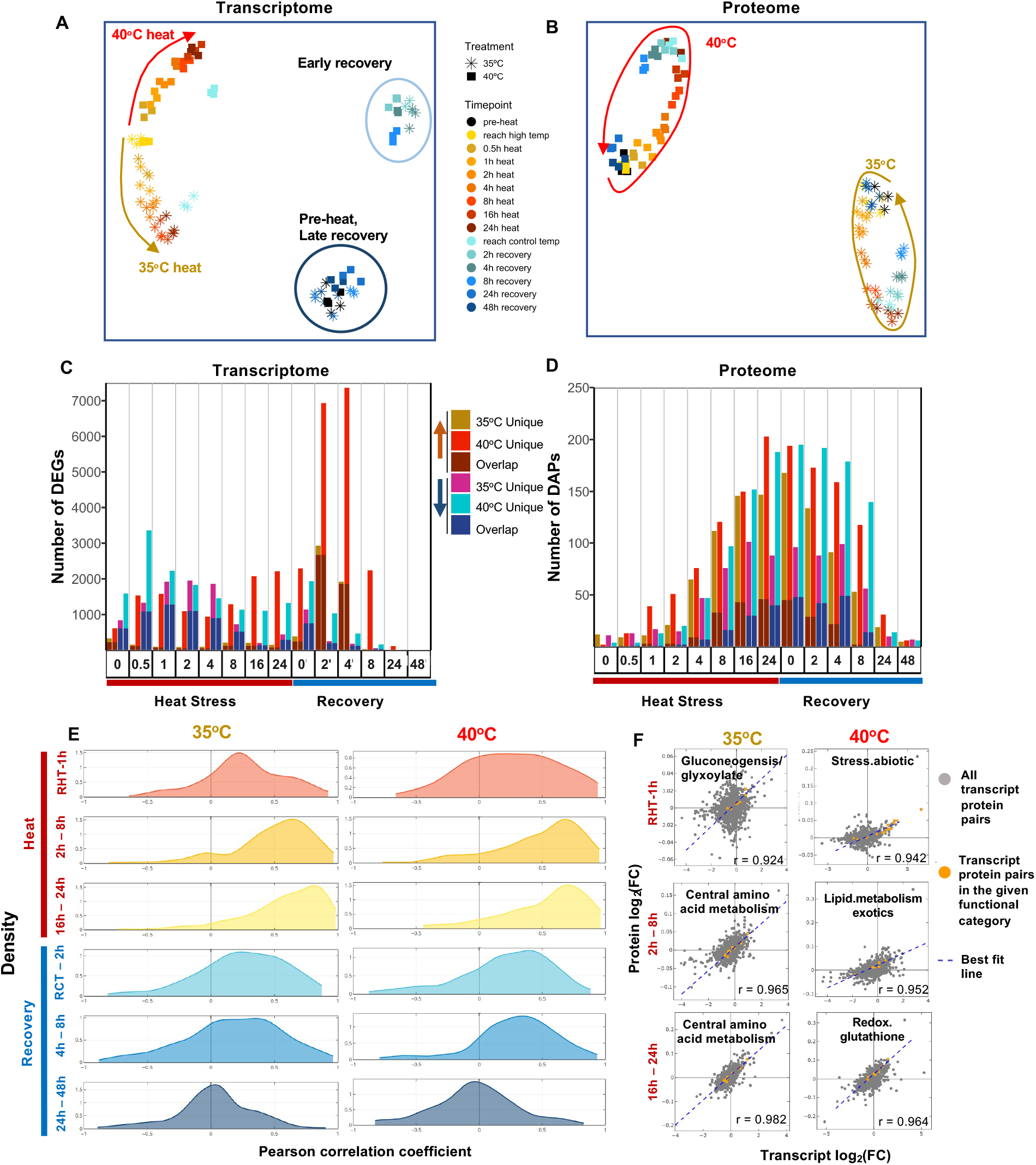
Transcriptomics and proteomics revealed distinct and dynamic responses during and after heat treatments of 35 and 40°C. **(A)** Uniform Manifold Approximation and Projection (UMAP) of Transcripts Per Million (TPM) normalized RNA-seq read counts and **(B)** UMAP of normalized protein intensities. Each data point represents all normalized counts from a single sample. Stars and squares represent algal samples with heat treatments of 35 or 40°C, respectively. Different colors represent different time points. Brown and red arrows show the movement through time of the 35 and 40°C treated samples, respectively. (**C, D**) Number of unique or overlapping Differentially Expressed Genes (DEGs) and Differentially Accumulated Proteins (DAPs) with heat treatment of 35 or 40°C at different time points, respectively. Each time point has four bars: the first bar represents genes up-regulated in 35°C, the second represents genes up-regulated in 40°C, the third represents genes down-regulated in 35°C, and the fourth represents genes down-regulated in 40°C. The bottom portion of the stacked bars represents genes/proteins that are differentially expressed in both treatment groups and the top portion represents genes that are uniquely differentially expressed in the given treatment group at that time point. Significant differential expression in transcriptome data was defined as absolute values of log_2_(fold-change, FC) > 1, FDR < 0.05, and absolute difference of TPM normalized read counts between treatment and pre-heat control ≥ 1. Significant differential accumulation of proteins was defined by Dunnett’s FWER < 0.05. (**E**) Analysis of correlation between transcripts and proteins revealed overall rising positive correlation during the heat treatment and a decreasing correlation during recovery. The time points were subdivided into three heat (red labels) and three recovery (blue labels) windows for 35°C (left) and 40°C (right) treated samples. The density plots of Pearson correlation coefficients between the fold-changes of transcripts and proteins are shown. X, Pearson correlation coefficient. Y, frequency density. Time points during heat: RHT, reach high temperature of 35 or 40°C; 1h, heat at 35 or 40°C for 1 h, similar names for other time points during heat. Time points during recovery: RCT, reach control temperature of 25°C for recovery after heat; 2h, recovery at 25°C for 2 h, similar names for other time points during recovery. See Supplemental Figure 7 for more information. (**F**) Scatter plots of all transcript-protein pairs at 35°C (left) and 40°C (right) for the three-heat time point bins shown in E. X and Y, transcript and protein log_2_FC, compared with pre-heat, respectively. Transcript-protein pairs are shown as gray dots. Best fit lines are shown in blue. The Pearson correlation coefficient is shown at the bottom right corner of each scatterplot. Transcript-protein pairs belonging to MapMan functional categories with the highest Pearson correlation coefficient in the given time point bin are shown in orange.

We employed differential expression modeling to identify differentially expressed genes (DEGs) that were overlappingly or uniquely up- or down-regulated during and after 35 or 40°C heat treatments (Figure 2C, Supplemental Dataset 1). The greatest number of DEGs were identified at 2- and 4-h recovery time points of 40°C treatment, while investigating the distribution of log_2_(fold-change) values for DEGs showed the greatest level of up-regulation at 0.5-h and down-regulation at 16-h of heat at 40°C (Supplemental Figure 4A). Overall, there were more DEGs at most time points in the 40°C treatment as compared to the 35°C treatment (Figure 2C). Analysis of differentially accumulated proteins (DAPs) that were overlappingly or uniquely up- or down-regulated during or after 35 and 40°C showed a smoother distribution of expression pattern than transcriptomic data (Figure 2D, Supplemental Dataset 2). Increasing numbers of DAPs were identified throughout the high temperature period, followed by a gradual reduction throughout the recovery period. The distribution of log_2_(fold-change) for DAPs at each time point also showed a smoother pattern than for transcriptome data (Supplemental Figure 4B). Through transcriptomic and proteomic analyses, we identified shared and unique responses for the 35 and 40°C treatment groups (Supplemental Figure 4C-F, Supplemental Dataset 1 and 2).

The global transcriptome analysis revealed three most dominant transcriptional patterns during the two treatments (Supplemental Figure 5A, B). The first constraint (λ1) divides the transcripts into acclimation and de-acclimation phases; the second constraint (λ2) separates control from disturbed conditions; the third constraint (λ3) shows a more fluctuating fine regulation. The amplitude difference between the treatments at 35 and 40°C suggests an overall higher regulatory activity during the 40 than 35°C treatment. However, there were a set of 108 genes uniquely up-regulated during 35 but not 40°C heat (Supplemental Figure 4C, Supplemental Dataset 1), including *GAPDH* (involved in gluconeogenesis, glycolysis, and Calvin-Benson Cycle), TAL1 (involved in pentose phosphate pathway), *COX15* (involved in mitochondrial assembly), and *CAV4* (encoding a putative calcium channel) (Supplemental Figure 5C-F). Additionally, when investigating the log_2_(fold-change) ratios of overlapping up- or down-regulated genes in both treatment groups at the same time point, we found that although many genes had higher differential expression with 40°C than 35°C treatment, some genes were more highly differentially expressed in the 35°C than the 40°C treatment group (Supplemental Figure 6, Supplemental Dataset 1). Taken together, these results indicate that 35°C induces a unique set of responses in Chlamydomonas that have not been previously described.

MapMan functional enrichment analysis of DEGs at each time point of the 35 or 40°C treatment showed that early induced shared responses to both heat treatments included canonical heat response pathways, protein folding, and lipid metabolism (Table 1, Supplemental Dataset 3). MapMan terms related to DNA synthesis, cell motility, protein processing, and RNA regulation were enriched in overlapping gene sets down-regulated during most time points of both 35 and 40°C heat treatments. DNA synthesis and repair MapMan terms were significantly enriched in genes up-regulated in both 35 and 40°C treated samples during the 2- and 4-h recovery time points. MapMan terms related to amino acid metabolism, mitochondrial electron transport, and purine synthesis were enriched in gene sets uniquely up-regulated during 35°C heat. Carbon fixation (e.g., carbon concentrating mechanism) and starch synthesis related MapMan terms were significantly enriched in gene sets uniquely up-regulated during 40°C heat. MapMan terms related to amino acid metabolism and mitochondrial electron transport were enriched in gene sets uniquely down-regulated during early heat of 40°C, in contrast to 35°C.

**Table 1:**
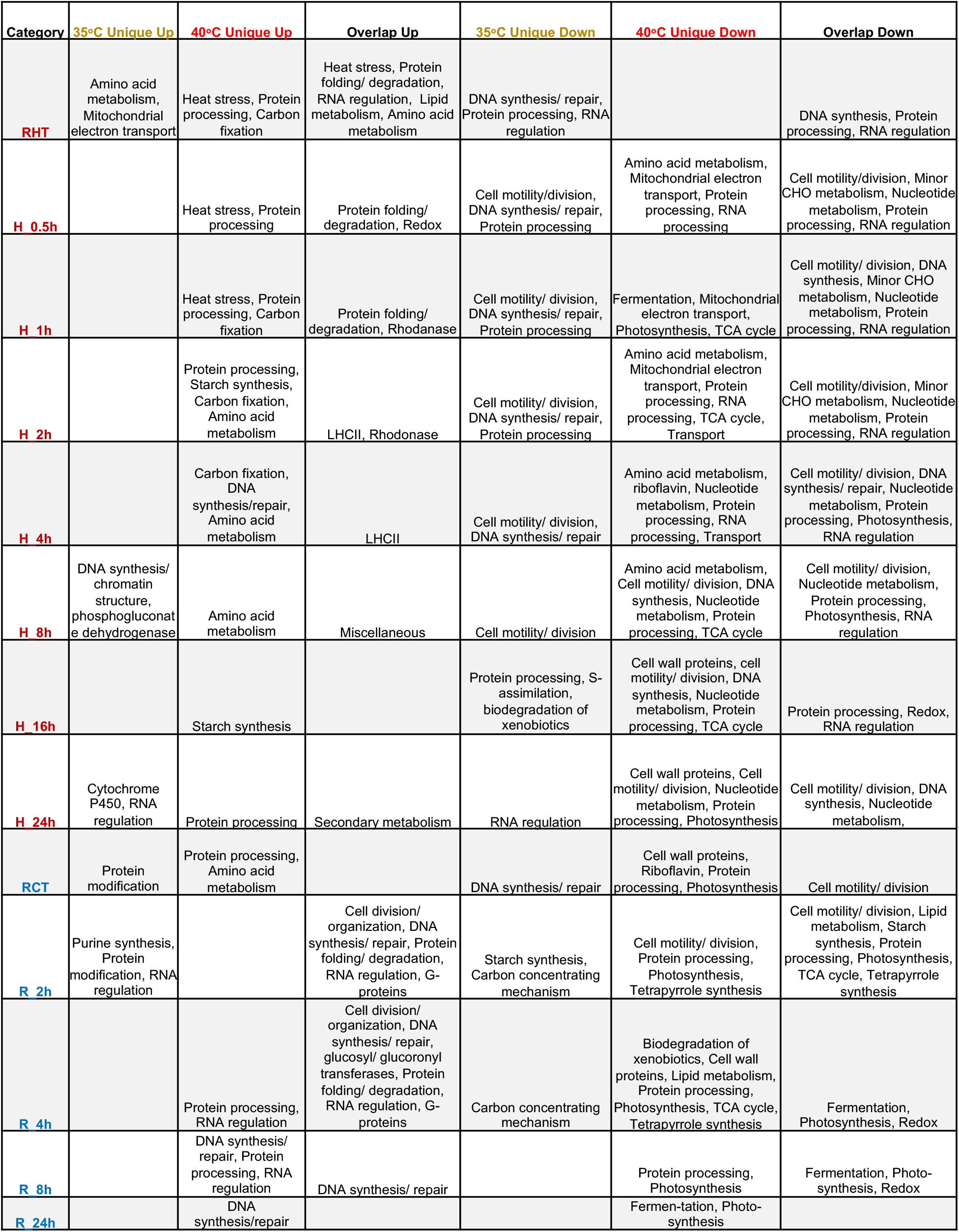
MapMan functional enrichment summary for genes uniquely or overlappingly differentially expressed in 35 and 40°C by time point. For each time point, genes identified as uniquely up/down-regulated or overlappingly up/down-regulated in 35 and 40°C treatments were used for MapMan functional enrichment analysis. Significantly enriched MapMan terms (FDR < 0.05) were sorted into broad functional categories (see Supplemental Dataset 3 for detailed functional information). Broad functional categories with at least one significantly enriched MapMan term are displayed in this table. RHT, reach high temperature of 35 or 40°C. H_1h, heat at 35 or 40°C for 1 h, similar names for other time points during heat. RCT, reach control temperature of 25°C for recovery after heat. R_2h, recovery at 25°C for 2 h, similar names for other time points during recovery.

### Transcript/protein correlation increased during heat but decreased during recovery

Investigation of Pearson correlation coefficients between log_2_(fold-change) values for transcripts and proteins grouped by MapMan functional categories revealed higher positive correlation between transcriptome and proteome during heat than recovery for both 35 and 40°C treatments (Figure 2E, F, Supplemental Figure 7). This indicates that transcriptional regulation dominates the heat period. Further investigation of Pearson correlation coefficients for individual MapMan terms showed that functional categories had varying correlation values throughout the course of high temperatures and recovery (Supplemental Dataset 4, 9). In early time points of the 35°C treatment, functional categories of gluconeogenesis/glyoxylate-cycle had the highest correlation between transcripts and proteins, followed by amino acid and lipid metabolism at later heat time points (Figure 2F). In early time points of the 40°C treatment, functional categories of abiotic stress had the highest correlation between transcripts and proteins, followed by lipid metabolism and redox pathways at later heat time points (Figure 2F). The proteins related to the MapMan bin gluconeogenesis/glyoxylate cycle increased during 35°C but decreased during 40°C, suggesting elevated and reduced gluconeogenesis/glyoxylate-cycle activity during 35 and 40°C heat treatment, respectively (Supplemental Figure 8). Isocitrate lyase (ICL1) is a key enzyme of the glyoxylate cycle in Chlamydomonas (Plancke et al., 2014). The transcript and protein as well as their correlation of ICL1 increased during 35°C heat but decreased during 40°C heat (Supplemental Dataset 9,10, gluconeogenesis _ glyoxylate cycle.html). Our results suggested 35°C treatment increased acetate uptake and assimilation, which may be suppressed by the 40°C treatment.

### Network modeling of transcriptome and proteome revealed expression patterns of key pathways during and after heat treatments

To investigate common transcriptional expression patterns throughout heat and recovery, we performed Weighted Correlation Network Analysis (WGCNA) on TPM-normalized RNA-seq data from 35 and 40°C transcriptomes (Figure 3A, Supplemental Dataset 5). Most genes with known roles in heat response belong to Transcriptomic Module 1 (TM1) with peak expression at 0.5-h heat, including *HSF*s and many *HSP*s. TM1 is also significantly enriched for MapMan terms related to protein folding, lipid metabolism, and starch synthesis. In total, there are 628 genes associated with this module, about 62% of which lack Chlamydomonas descriptions and 25% have no ontology annotations, suggesting novel genes with putative roles in heat response that have been previously undescribed. TM2 has peak expression at 1-h heat and is significantly enriched for MapMan terms relating to the TCA cycle and carbon concentrating mechanism. TM3 has peak expression at the beginning of both heat and recovery periods and is significantly enriched for MapMan terms related to RNA regulation and chromatin remodeling, highlighting the extensive changes in transcriptional regulation caused by changing temperatures. TM4 contains 137 genes that have sustained increased expression throughout 35°C, with slightly reduced or no change in expression in 40°C. This module is significantly enriched for MapMan terms related to amino acid metabolism, ferredoxin, and fatty acid synthesis/elongation. TM4 may represent genes that are unique for 35°C responses. TM5 shows increased expression throughout the heat period at 40°C. This module is enriched for ABA synthesis/degradation, suggesting potential interaction between the ABA pathway and algal heat responses. It has been reported that heat treated Arabidopsis and rice leaves had increased ABA contents and exogenous ABA application improved heat tolerance in WT rice plants (Balfagón et al., 2019, 1; Li et al., 2020a). ABA is reported to be involved in tolerance to oxidative stress (Yoshida et al., 2003), HCO3^−^ uptake (Al-Hijab et al., 2019), and stress acclimation in Chlamydomonas (Colina et al., 2019). TM6 displays similar expression patterns in both 35 and 40°C treatments, with increased expression during early heat and reduced expression during early recovery. This module is enriched for genes in the Calvin cycle, PSII biogenesis, and tetrapyrrole synthesis. TM7 shows increased expression in 40°C uniquely in late heat, which continues through the early and mid-recovery periods. This module is notably enriched for protein degradation, ubiquitin E3 RING, and autophagy, which may indicate the need to degrade certain proteins and cellular components during and after prolonged acute high temperature at 40°C. TM8 shows increased expression only at the reach 25°C time point immediately after the cooling from the 35 and 40°C treatments. This module is notably enriched for bZIP transcription factors, oxidases, calcium signaling, and protein posttranslational modification, which may be partially controlling the broad changes observed when recovering from high temperatures. TM9 and TM10 both have peaks in expression in recovery (2 h, 4 h for TM9; and 8 h for TM10) and have significantly enriched MapMan terms relating to DNA synthesis. TM11 peaks in expression at 8-h recovery and is significantly enriched for MapMan terms relating to protein posttranslational modification, consistent with decreased correlation between transcript and protein levels at late recovery stages (Figure 2E). TM12 shows reduced expression during the early recovery period and is significantly enriched for MapMan terms relating to photosynthetic light reactions and protein degradation.

**Figure 3:**
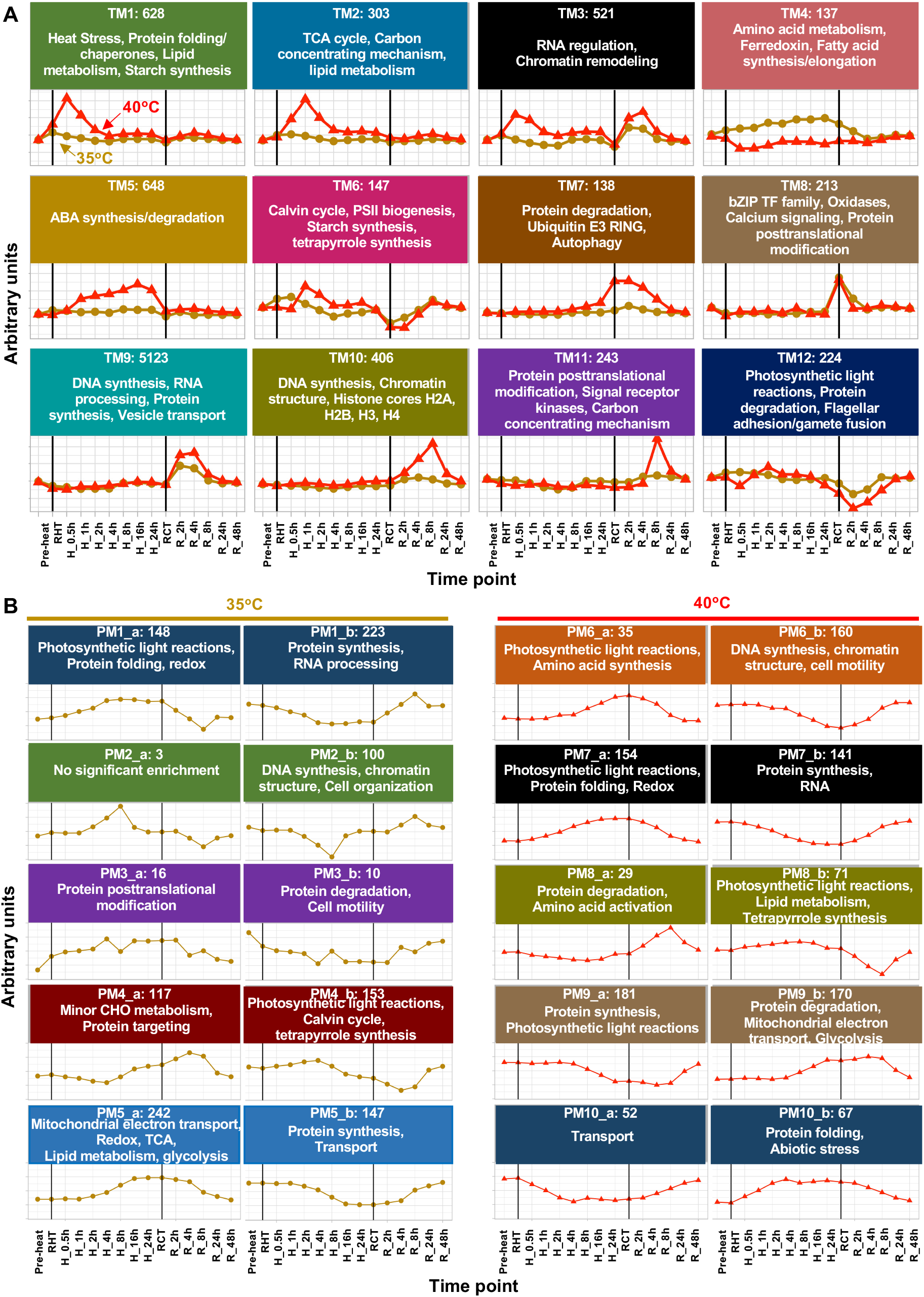
Transcriptomic and proteomic network modeling revealed differential regulation of key biological pathways with treatments of 35 or 40°C. **(A)** Weighted correlation network analysis of transcriptome data identified gene modules with similar expression patterns. TM: transcriptomic module. The consensus gene expression patterns for 40°C (red triangle) and 35°C (brown circle) for each module are displayed. The number at the top of each facet represents the total number of genes significantly associated with the given module (ANOVA, FDR < 0.05, genes can only belong to a single module). Select statistically significantly enriched MapMan functional terms are displayed in each facet (FDR < 0.05). **(B)** Correlation networks of protein abundance over time courses for 35 and 40°C heat treatments. PM, proteomic module. For each module, the eigenvectors as aggregated signal shape is depicted. Prominent functional terms that were enriched in the respective module are given for proteins correlating positively (PM_a) or negatively (PM_b) to their corresponding eigenvector (FDR < 0.05). The number at the top of each facet represents the total number of proteins significantly associated with the given module. Black vertical lines indicate the start and end of the heat treatment at either 35 or 40°C. Pre-heat, before heat treatments. RHT, reach high temperature of 35 or 40°C. H_1h, heat at 35 or 40°C for 1 h, similar names for other time points during heat. RCT, reach control temperature of 25°C for recovery after heat. R_2h, recovery at 25°C for 2 h, similar names for other time points during recovery. The y axes are in arbitrary units.

We verified the expression patterns of several key pathways of interest and transcription factors by visualizing log_2_(fold-change) values from differential expression modeling (Supplemental Figures 9,10, Supplemental Dataset 6). The RNA-seq results of select pathways from differential expression modeling were highly consistent with WGCNA modeling and provided gene-level resolution of interesting trends within these pathways. The down-regulation of select transcripts involved in photosynthetic light reactions during early recovery was also verified by RT-qPCR (Supplemental Figures 9 J, K).

Network modeling was performed separately for the proteomes of the 35 and 40°C treated samples due to differences in peptides identified through LC-MS/MS between the two treatment groups and the relatively smaller number of proteins identified compared to transcripts. This analysis identified common expression patterns (Proteomics Module, PM) in the proteome data (Figure 3B, Supplemental Dataset 7). Prominent functional terms that were enriched in the respective module are given for proteins correlating positively (PM_a) or negatively (PM_b) to their corresponding eigenvector (FDR < 0.05). Proteins that increased during both 35 and 40°C heat treatments are enriched for MapMan terms relating to part of photosynthetic light reactions, protein folding, and redox (PM1_a, PM6_a, PM7_a). Proteins that increased during the recovery phase after both heat treatments are enriched for MapMan terms relating to protein synthesis (PM1_b, PM7_b), DNA synthesis and chromatin structure (PM2_b, PM6_b). Proteins related to photosynthetic light reactions first decreased and then increased during the recovery of both treatments (PM1_a, PM4_b, PM8_b). Unique responses for the 35°C treatment include proteins related to mitochondrial electron transport and lipid metabolism increased during heat (PM5_a) and proteins related to RNA processing and cell organization increased during the recovery phase after 35°C heat (PM1_b, PM2_b). Unique responses for the 40°C treatment include proteins related to abiotic stress increased during heat (PM10_b) and proteins related to cell motility and RNA increased during the recovery phase after 40°C heat (PM6_b, PM7_b). Network modeling of transcriptome and proteome data yielded consistent patterns for several key pathways during and after heat treatments, e.g., heat responses, photosynthetic light reactions, and DNA synthesis.

### Heat at 35°C synchronized the cell cycle while 40°C arrested it

The increased transcript and protein levels related to DNA synthesis during recovery (Figure 3) prompted us to investigate the expression pattern of cell cycle related genes because DNA synthesis usually takes place immediately before and during cell division in Chlamydomonas (Cross and Umen, 2015). Both 35 and 40°C disrupted the expression pattern of cell cycle genes as compared to the pre-heat level, which recovered after 8-h heat treatment at 35°C but not during the entire 40°C heat treatment (Figure 4A). Most cell cycle related genes had increased expression during 2- and 4-h of recovery in both treatment groups.

**Figure 4.**
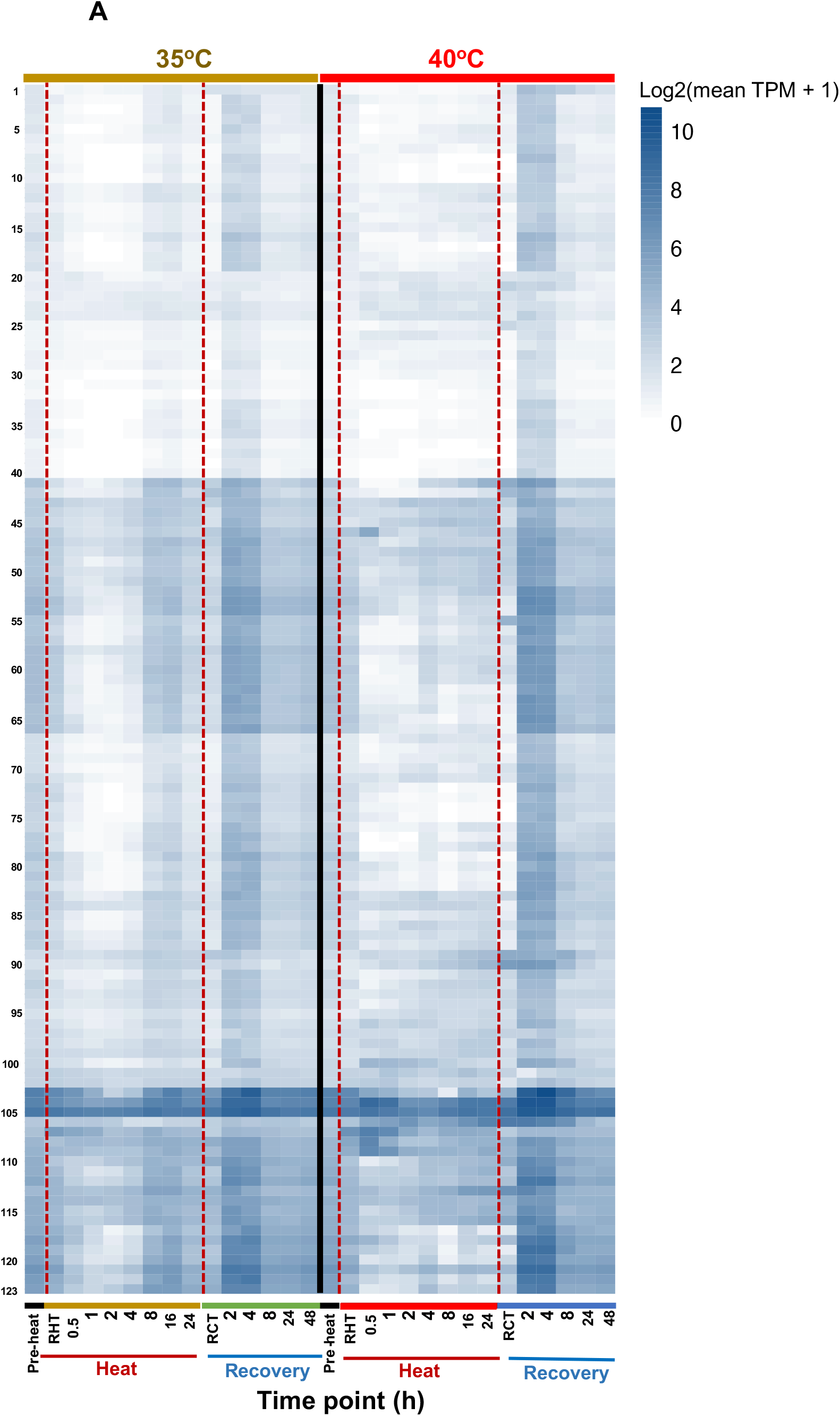
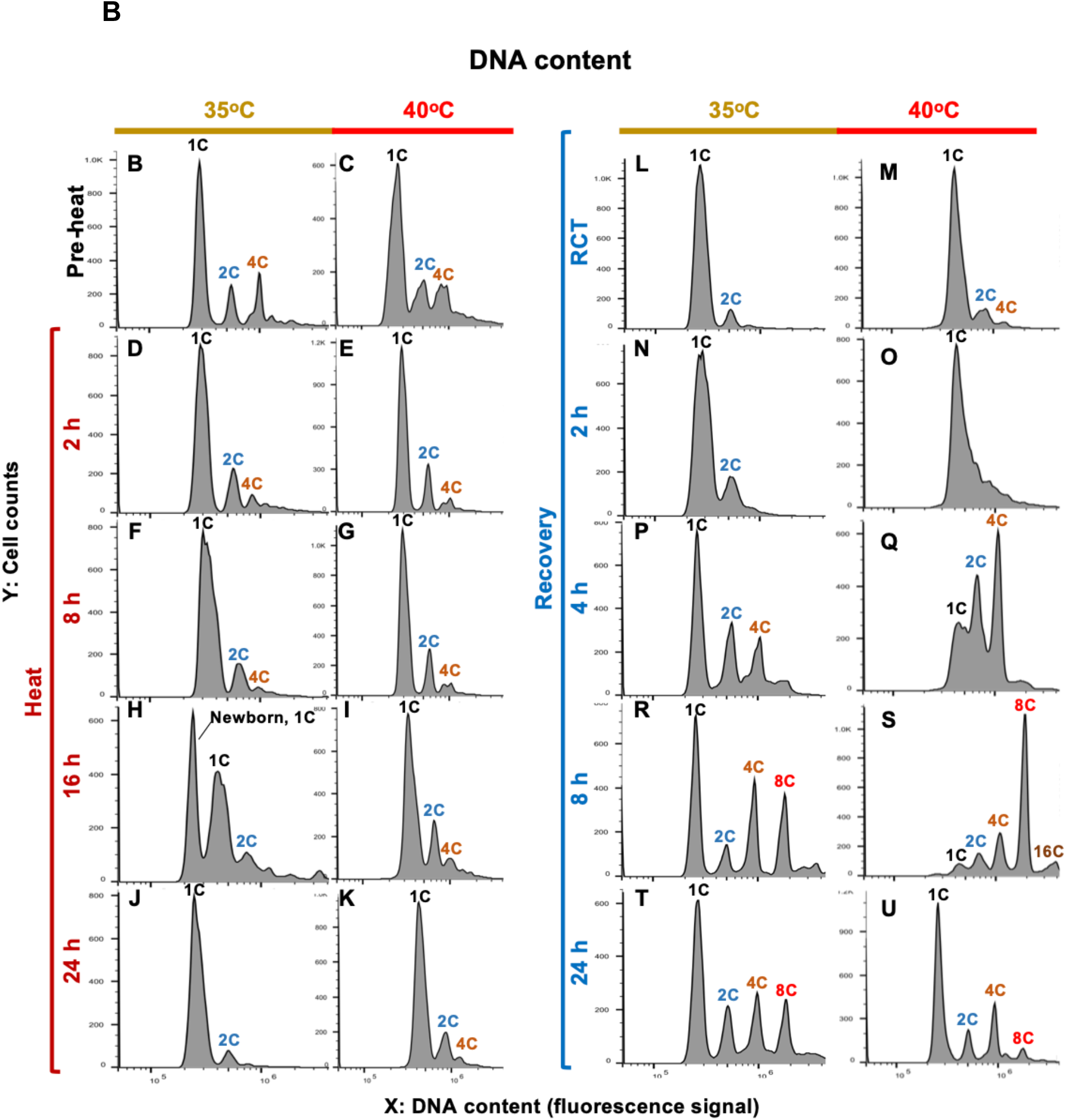
Heat at 35°C synchronized the cell cycle while heat at 40°C arrested it. **(A)** Expression of cell cycle genes during and after heat treatment of 35 or 40°C. Genes used for the expression pattern analysis are listed in Supplemental Dataset 6. Gene expression patterns for treatments of 35°C (left) and 40°C (right) are displayed as log_2_(mean TPM value + 1). TPM, transcripts per million. Darker blue colors indicate higher expression. Red dashed lines indicate the start and end of the heat treatments. Pre-heat, before heat treatment. RHT, reach high temperature of 35 or 40°C. RCT, reach control temperature of 25°C for recovery after heat. (**B-U**) FACS (fluorescence-activated cell sorting) analysis of the DNA content in algal samples harvested at different time points before, during, and after heat treatment at 35 or 40°C. For each figure panel, X is DNA content determined by plotting a histogram of fluorescence level (area of the fluorescence channel signal) in log-scale; Y is the cell counts in linear scale. DNA copy No. are labeled on top of each corresponding DNA content peak, 1C (single DNA copy number), 2C, 4C, 8C, 16C.

During the 40°C heat treatment, we observed irregular expression patterns of cell cycle related genes as compared to pre-heat, which led us hypothesize that the cell cycle had been arrested during 40°C. To investigate this hypothesis, we quantified cellular DNA content using flow cytometry (Figure 4B-U). Pre-heat cultures showed typical asynchronous populations: most cells had 1C (single DNA copy number) while a small fraction of cells had 2C and 4C (Figure 4B, C). After 16-h at 35°C heat treatment, the broad 1C size distribution from 8-h split; some of the bigger cells went to 2C and the rest of the small cells stayed at 1C (Figure 4F, H). The 2C population then divided by 24-h at 35°C, resulting in almost exclusively small 1C cells with the same cell size as 1C cells at pre-heat, suggesting culture synchrony (Figure 4J). The cell division during 8-16 h of heat at 35°C was consistent with the recovery of cell cycle genes at the same time points (Figure 4A). After 2-h of recovery at 25°C following 35°C heat, there were much fewer 2C and 4C cells than pre-heat, suggesting a partially synchronized population, until an almost complete recovery by 4-h at 25°C (Figure 4L, N, P, R, T). These results indicated that the 35°C treatment synchronized cell division in Chlamydomonas.

Heat treatment at 40°C inhibited DNA replication and cell division, with 3 peaks of 1C, 2C, and 4C persisting during all 40°C heat time points (Figure 4C, E, G, I, K). By the end of the 24-h heat treatment at 40°C, the three cell populations had a much larger cell size, as evidenced by the right shift of the DNA peaks due to an increased cell size background effect (Figure 4K, more information of the background effect can be found in the Methods). After 2-4 h recovery from 40°C, cells started to replicate DNA between 2- and 4-h of recovery at 25°C, resulting in reduced 1C but increased 2C and 4C cell populations (Figure 4Q). DNA replication continued until 8-h of recovery, resulting in the accumulation of high-ploidy level cells, ranging from 1C to 16C (Figure 4S). By 24-h recovery, the cellular DNA content had almost recovered to the pre-heat level (Figure 4U).

### Cytological parameters confirmed cell cycle arrest during 40°C heat

Brightfield images and cell size quantification of algal cells showed that the 40°C treated cells had continuously increased cell size throughout the high temperature period, followed by gradual recovery to the pre-stress level after returning to 25°C for 24 h (Figure 5A, B, C). The quantity of chlorophyll, carotenoids, protein, and ROS per cell all increased during the 40°C heat treatment and gradually recovered after heat (Figure 5D-G). When normalized to cell volume, only chlorophyll and carotenoid contents increased during 40°C heat treatment while no change of protein and ROS contents was observed (Supplemental Figure 11A-D). The ratios of chlorophyll a/b and chlorophyll/carotenoid decreased during 40°C heat (Supplemental Figure 11E, F). The changes of chlorophyll a and b were consistent with that of total chlorophyll (Supplemental Figure 10G-J). Cell diameter, cell volume, chlorophyll and carotenoid contents only changed transiently after shifting to 35°C and after shifting back to 25°C (Figure 5B-E, Supplemental Figure 11).

**Figure 5.**
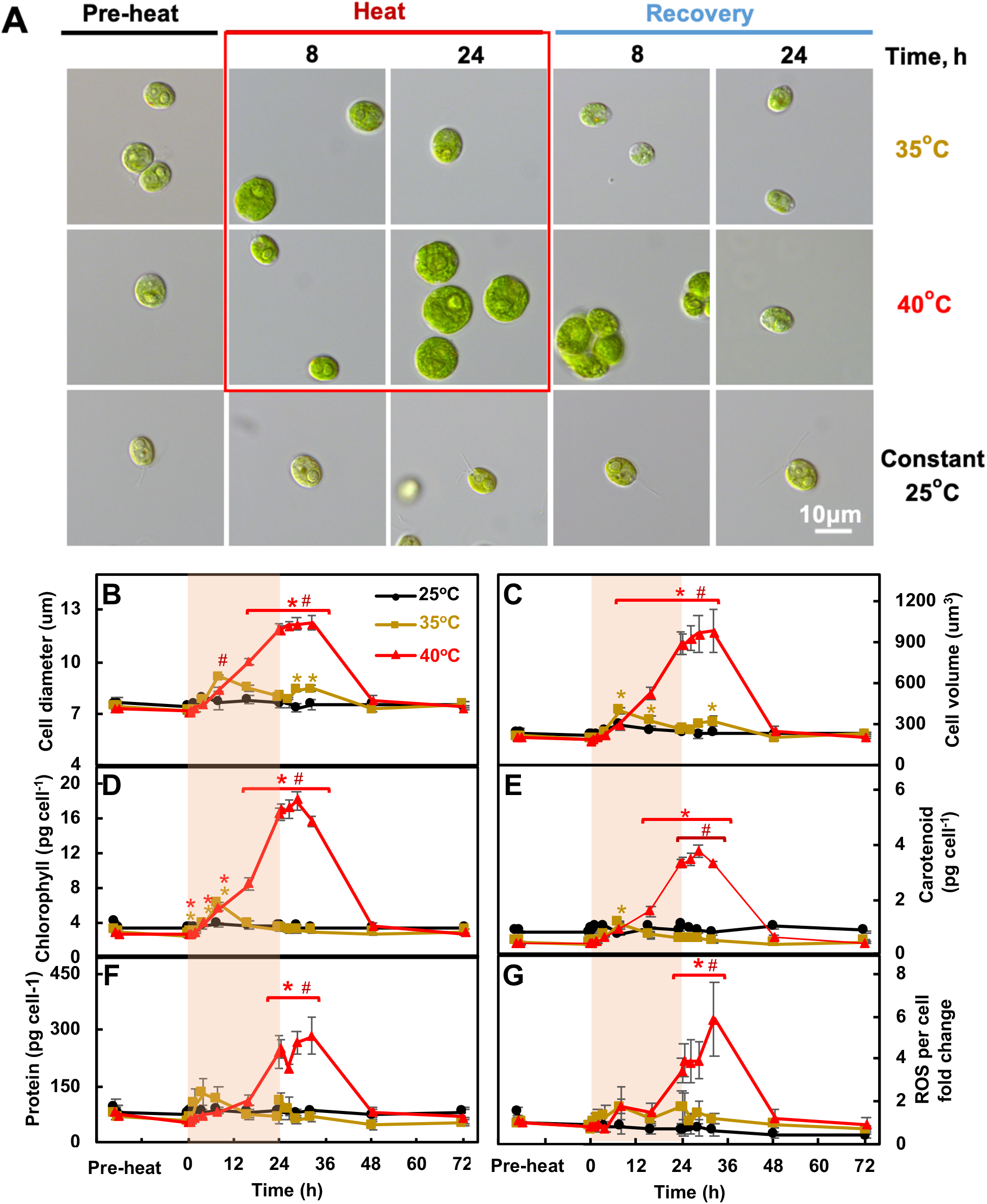
Heat of 40°C persistently increased cell size and cellular levels of pigments, protein, and ROS while these effects were transient with 35°C heat. **(A)** Light microscopic images of Chlamydomonas cells. **(B, C)** Cell diameters and volume determined using a Coulter Counter. **(D-F)** Total chlorophyll, carotenoid, protein content per cell. **(G)** Fold-change of reactive oxygen species (ROS) levels per cell quantified using CM-H2DCFDA ROS indicator. Mean ± SE, *n*=3 biological replicates. Black, brown and red curves represent experiments with constant 25°C, or with treatments at 35 or 40°C respectively. Red shaded areas depict the duration of high temperature. Statistical analyses were performed using a two-tailed t-test assuming unequal variance by comparing treated samples with pre-heat (*, p<0.05, the colors of asterisks match the treatment conditions) or between 35 and 40°C at the same time point (#, p<0.05).

### Heat at 40°C impaired photosynthesis while the effects at 35°C were minor

We hypothesized that heat treatments might affect photosynthetic activities based on the changes of pigment contents (Figure 5D, E), differentially regulated genes related to photosynthesis (Supplemental Figure 9A, B), and the kinetics of transcripts and proteins related to the MapMan bin photosynthetic light reactions (Figure 6, Supplemental Dataset 10). Most transcripts related to photosynthetic light reactions decreased during the early recovery from both heat treatments, followed by a gradual returning to the pre-stress levels (Figure 6A, B, E, F, I, J), consistent with the network modeling data (Figure 3). Proteins related to PSI, LHCII (light harvesting complexes II), and LHCI increased during both heat treatments and decreased back to the pre-heat levels during the recovery (Figure 6C, D, G, H). Proteins related to the ATP synthase decreased during the 40°C heat treatment and the early/middle recovery of both 35 and 40°C heat treatments (Figure 6K, L). Overall, the kinetics of transcripts and proteins related to the MapMan bin photosynthetic light reactions showed similar trends in both 35 and 40°C treatments.

**Figure 6.**
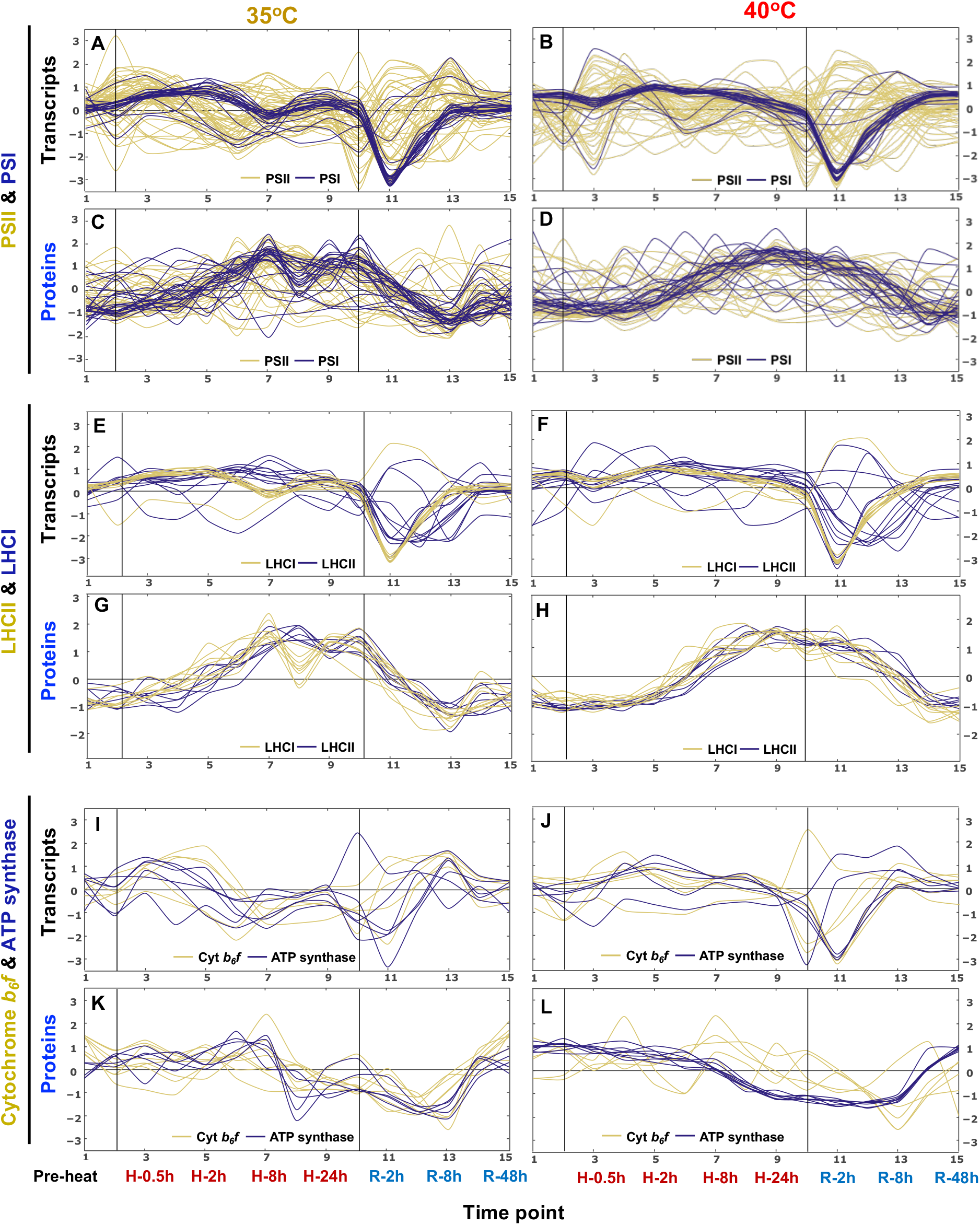
Transcripts and proteins related to photosynthetic light reactions changed dynamically during and after 35 and 40°C heat treatments. Transcript (**A, B, E, F, I, J**) and protein (**C**, **D, G, H, K, L**) signals related to the MapMan bin photosynthetic light reactions, including PSII and PSI (**A-D**), LHCII and LHCI (**E-H**), Cytochrome *b_6_f* and ATP synthase (**I-L**), were standardized to z scores (standardized to zero mean and unit variance) and are plotted against equally spaced time point increments. The black vertical lines indicate the start and end of heat treatments at 35°C (**A**, **C, E, G, I, K**) and 40°C (**B, D, F, H, J, L**), respectively. Time points are labeled at the bottom. Timepoint 1: pre-heat. Time points 2-9, heat treatment at 35 or 40°C, including reaching high temperature, 0.5, 1, 2, 4, 8, 16, 24 h during heat; time points 10-15, recovery phase after heat treatment, including reaching control temperature, 2, 4, 8, 24, 48 h during recovery. See the interactive figures with gene IDs and annotations in Supplemental Dataset 10, the groups of PS.lightreaction (PS for photosynthesis).

To investigate whether these pronounced changes of proteins related to LHCI, LHCII, PSI and ATP synthase under 35 and 40°C affected photosynthesis, we measured various photosynthetic parameters during and after the heat treatments (Figures 7-9). The PSII operating efficiency and linear electron flow rates largely decreased during 40°C heat (especially under light intensities exceeding the growth light of 100 µmol photons m^−2^ s^−1^) while the 35°C heat treatment did not extensively affect these photosynthetic parameters (Figure 7A-D). The Q_A_ redox state reflects the balance between excitation energy at PSII and the rate of the Calvin-Benson Cycle (Fu et al., 2017; Głowacka et al., 2018). The reduced Q_A_ is proportional to the fraction of PSII centers that are closed (Baker et al., 2007). Under 35°C, Q_A_ was more oxidized than pre-heat at 4- and 8-h heat and then more reduced by the end of 24-h heat; Q_A_ was more reduced at time points later than 4-h during the 40°C heat (Figure 7E, F), suggesting more reduced and less active electron transport chain under 40°C than 35°C. Both 35 and 40°C increased the formation of NPQ; however, the increased NPQ was steady during 4-24 h of heat at 35°C while during 40°C heat, NPQ first increased to a maximum at 8-h heat, then decreased by 24-h heat (Figure 7G, H), suggesting that accumulative heat damages under prolonged exposure to 40°C eventually exceeded the photoprotective capacity of NPQ. Relative PSII antenna size increased during the 40°C heat treatment, while it increased transiently during the 35°C heat treatment (Figure 7I, Supplemental Figure 12).

**Figure 7.**
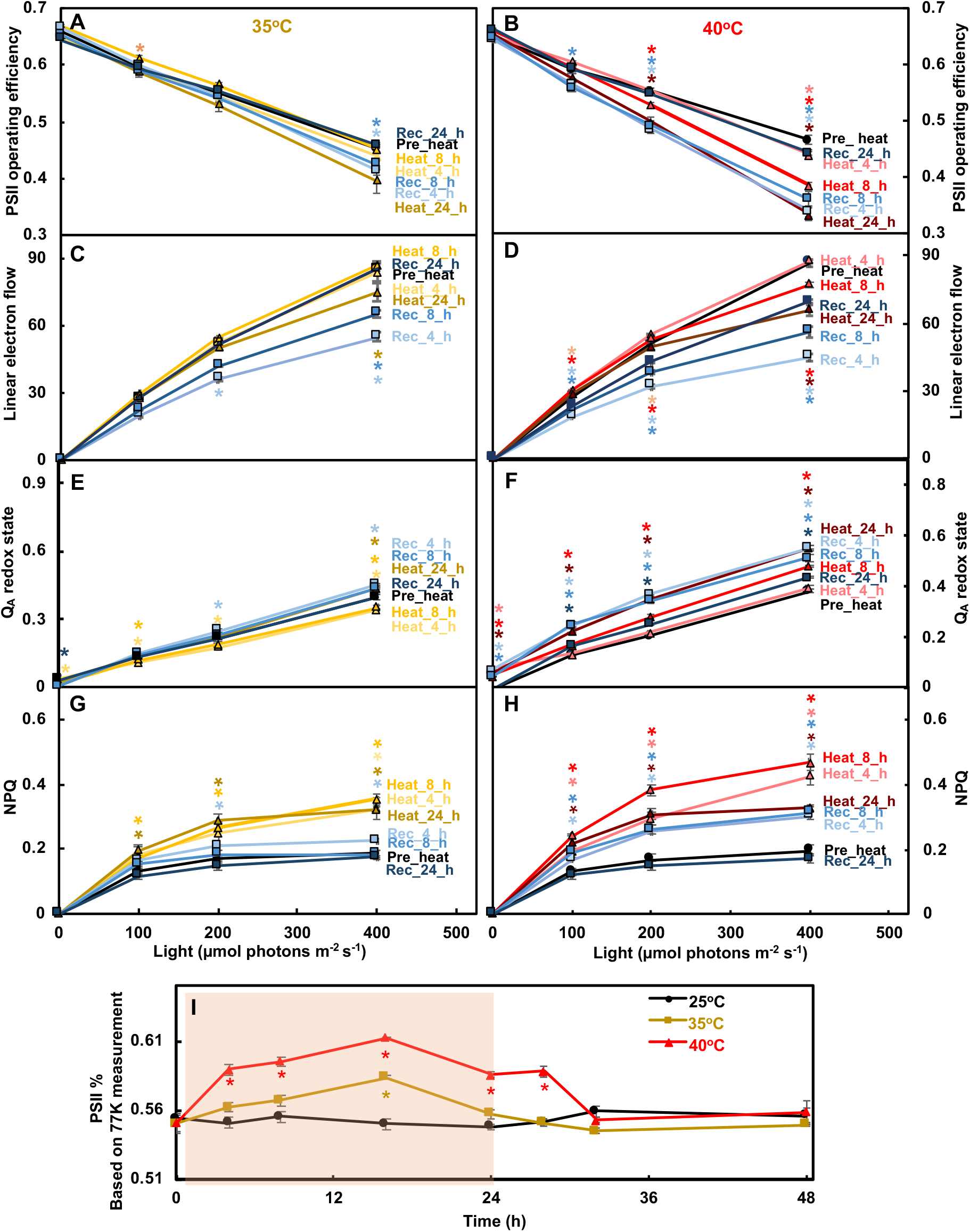
Heat of 40°C impaired PSII efficiency more than heat of 35°C while both induced NPQ. Algal cultures harvested from PBRs before, during and after heat treatment at 35°C (**A, C, E, G, I**) or 40°C (**B, D, F, H, I**) were used for measuring photosynthetic parameters. (**A-H**) Photosynthetic parameters measured using room temperature chlorophyll fluorescence. (**A, B**) PSII operating efficiency. (**C, D**) Linear electron flow, accounting for the changes of PSII antenna size during the treatments as in (**I**). (**E, F**) Q_A_ redox state, the redox state of chloroplastic quinone A (Q_A_), the primary electron acceptor downstream of PSII; the bigger number of Q_A_ redox state means more reduced Q_A_. (**G**, **H**) Non-photochemical quenching, NPQ. (**I**) Relative PSII antenna fraction, percentage of light distributed to PSII measured by 77 K chlorophyll fluorescence. Mean ± SE, *n*=3 biological replicates. Statistical analyses were performed using two-tailed t-test assuming unequal variance by comparing treated samples with the pre-heat samples under the same light (**A-H**) or constant 25°C samples at the same time point (**I**). A-H, p values were corrected by FDR. *, p<0.05, the colors and positions of asterisks match the treatment conditions and time points, respectively.

**Figure 8.**
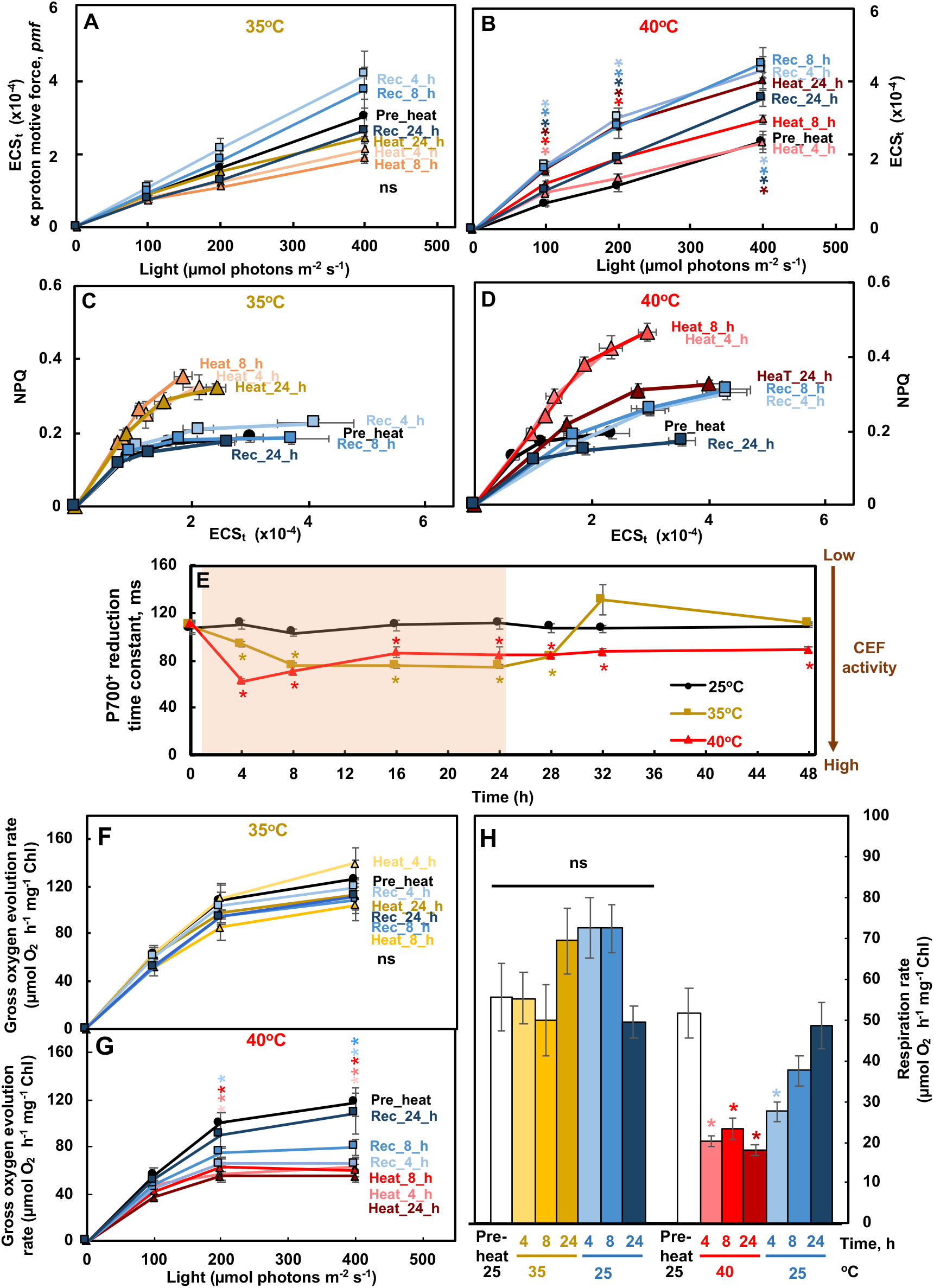
Heat at 40°C induced transthylakoid proton motive force, NPQ, CEF, but reduced O_2_ evolution and respiration rates in Chlamydomonas cells. (**A, B**) Heat treatment of 40°C increased the transthylakoid proton motive force (*pmf*). ECS_t_, measured by electrochromic shift (ECS), represents the transthylakoid *pmf*. (**C, D**) NPQ was more sensitive to ECS_t_ (or *pmf*) during both heat treatments, with higher NPQ produced at a given *pmf*. NPQ, non-photochemical quenching, measured using room temperature chlorophyll fluorescence. (**E**) Both 35°C and 40°C induced the activity of cyclic electron flow around PSI (CEF) although with different dynamics and reversibility. P700^+^ reduction to measure CEF in the presence of 10 µmol DCMU to block PSII activity; the smaller P700^+^ reduction time constant indicates faster P700^+^ reduction and higher CEF activity. The red shaded area depicts the duration of the high temperature. (**F, G, H**) Gross O_2_ evolution rates and respiration rates were reduced during the 40°C heat treatment, measured using a Hansatech Chlorolab 2 Clark-type oxygen electrode. Mean± SE, *n*=5 biological replicates. Statistical analyses were performed using two-tailed t-test assuming unequal variance by comparing treated samples with the pre-heat samples under the same light (**A, B, F, G, H**) or constant 25°C samples at the same time point (**E**). A, B, F, G, p values were corrected by FDR. *, p<0.05, the colors and positions of asterisks match the treatment conditions and time points, respectively. Not significant, ns.

**Figure 9.**
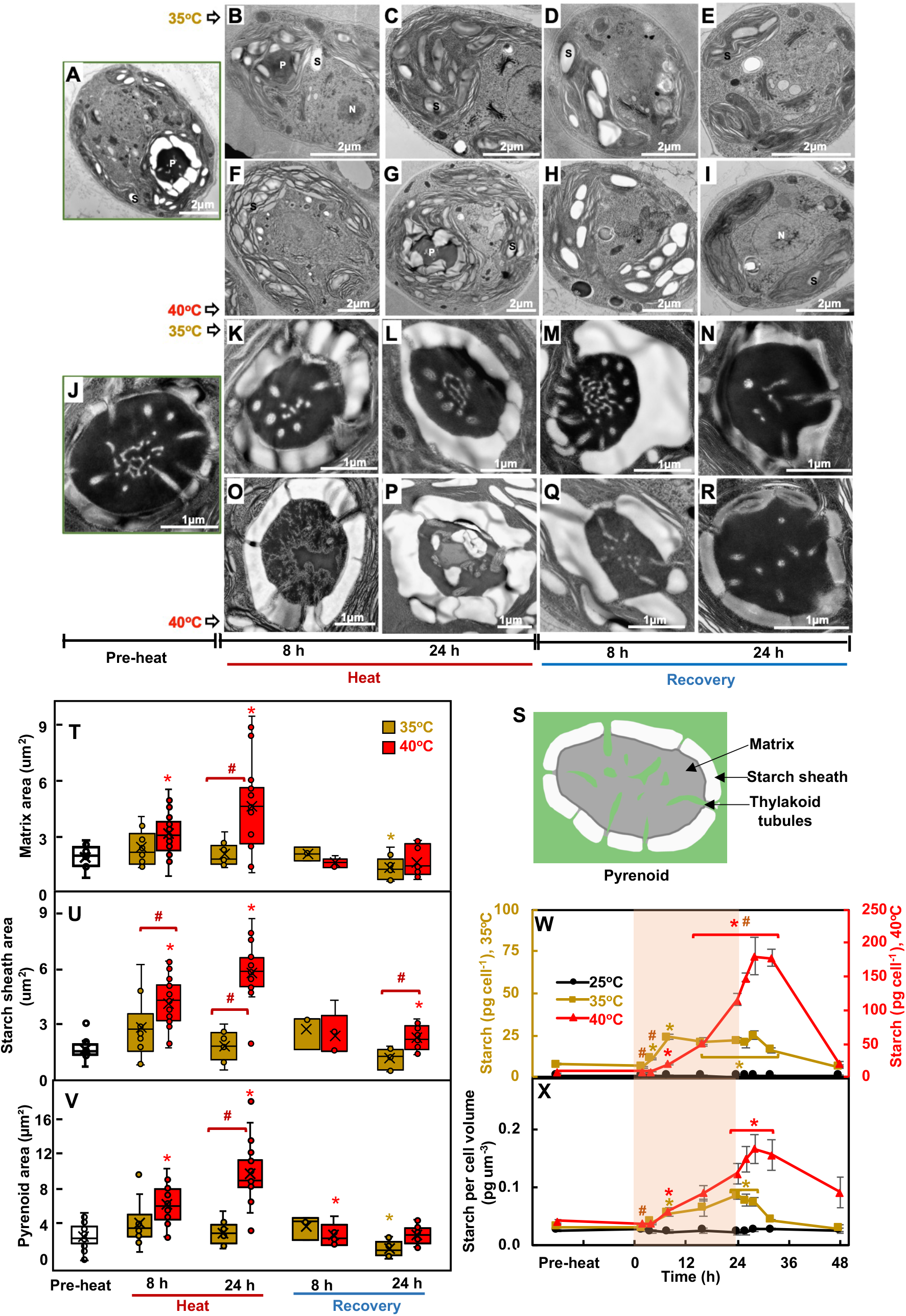
Both 35 and 40°C treatments stimulated starch accumulation while only 40°C altered thylakoid and pyrenoid ultrastructure. **(A-R)** Representative transmission electron microscopy (TEM) images of algal cells or pyrenoids at different time points before (**A, J**), during, and after heat treatment at 35°C (**B-E, K-N**) or 40°C (**F-I, O-R**). (**S**) Cartoon representation of Chlamydomonas pyrenoid structure. (**T, U, V**) Areas of pyrenoid matrix, starch sheath and the whole pyrenoids, respectively, quantified using ImageJ and TEM images from algal samples harvested before, during, and after heat treatments of either 35°C (brown) or 40°C (red) at the indicated time points. The data is presented as boxplot based on Tukey-style whiskers. Median values are represented by the horizontal black lines and mean values by the X sign inside each rectangular box. (**W, X**) Starch quantification using starch assay kits. Values are mean ± SE, *n* = 3 biological replicates. The red shaded areas depict the duration of the high temperature. (**T-X**) Statistical analyses were performed using two-tailed t-tests assuming unequal variance by comparing treated samples with pre-heat (*, p<0.05, the colors of asterisks match the treatment conditions) or between 35 and 40°C at the same time point (#, p<0.05).

Additionally, we performed electrochromic shift (ECS) measurements to monitor the effects of heat on the transthylakoid proton motive force (*pmf*, estimated by ECS_t_) (Figure 8A, B). The *pmf* increased particularly at late time points during the 40°C treatment, followed by a very slow and partial recovery after shifting cells back to 25°C. No significant changes in *pmf* were observed during and after the 35°C treatment. Proton conductivity decreased during both 35 and 40°C heat treatments, suggesting reduced ATP synthase activity (Supplemental Figure 13A, B), consistent with reduced abundance of proteins related to ATP synthase (Figure 6K, L). During both heat treatments, NPQ formation became more sensitive to *pmf*, with higher NPQ formed at a given *pmf* compared to the pre-heat condition (Figure 8C, D), consistent with a previous report in tobacco plants (Zhang et al., 2009). The increased sensitivity of NPQ was collapsed by the end of the 24-h heat treatment at 40°C (Figure 8D). P700 measurement revealed that the activity of cyclic electron flow around PSI (CEF) increased, and PSI became more reduced during both 35 and 40°C heat which recovered quickly after 35°C treatment but much more slowly after 40°C treatment (Figure 8E, Supplemental Figure 13C). Furthermore, gross photosynthetic O_2_ evolution rates and dark respiration rates hardly changed during the 35°C treatment but dropped significantly during the 40°C heat treatment (Figure 8F, G, H). Photosynthetic parameters had no significant changes in cultures maintained under constant 25°C (Supplemental Figure 14).

### Heat at 40°C altered thylakoid and pyrenoid ultrastructure

The effects of high temperatures on photosynthesis prompted us to investigate cellular ultrastructure using transmission electron microscopy (TEM) (Figure 9A-R). Thylakoids became disorganized and loosely packed in cells treated with 40°C (Figure 9F, G). Investigation of pyrenoid ultrastructure (Figure 9S) showed that cells treated with 40°C had altered pyrenoid matrices and absence of thylakoid tubules inside the pyrenoid (Figure 9O, P), suggesting an inefficient carbon concentrating mechanisms (CCM). No changes in pyrenoid ultrastructure were observed in cells treated with 35°C (Figure 9K, L). ImageJ quantification of pyrenoid structures showed that cells treated with 40°C had increased pyrenoid areas, which was attributed to increase areas of both pyrenoid matrix and starch sheath (Figure 9T, U, V). The increased pyrenoid size was abolished after 8 h of recovery. Biochemical quantification of starch contents showed that both 35 and 40°C treatments increased starch levels per cell and per cell volume, which decreased during recovery (Figure 9W, X). At the end of the heat treatments, cells exposed to 40°C had a higher starch content per cell than those exposed to 35°C, but the differences between the two treatments were not significant per cell volume.

## Discussion

We investigated how Chlamydomonas cells respond to moderate (35°C) and acute (40°C) high temperature at systems-wide levels (Figure 1C). Our results show that 35 and 40°C triggered shared and unique heat responses in Chlamydomonas (Figure 10).

**Figure 10.**
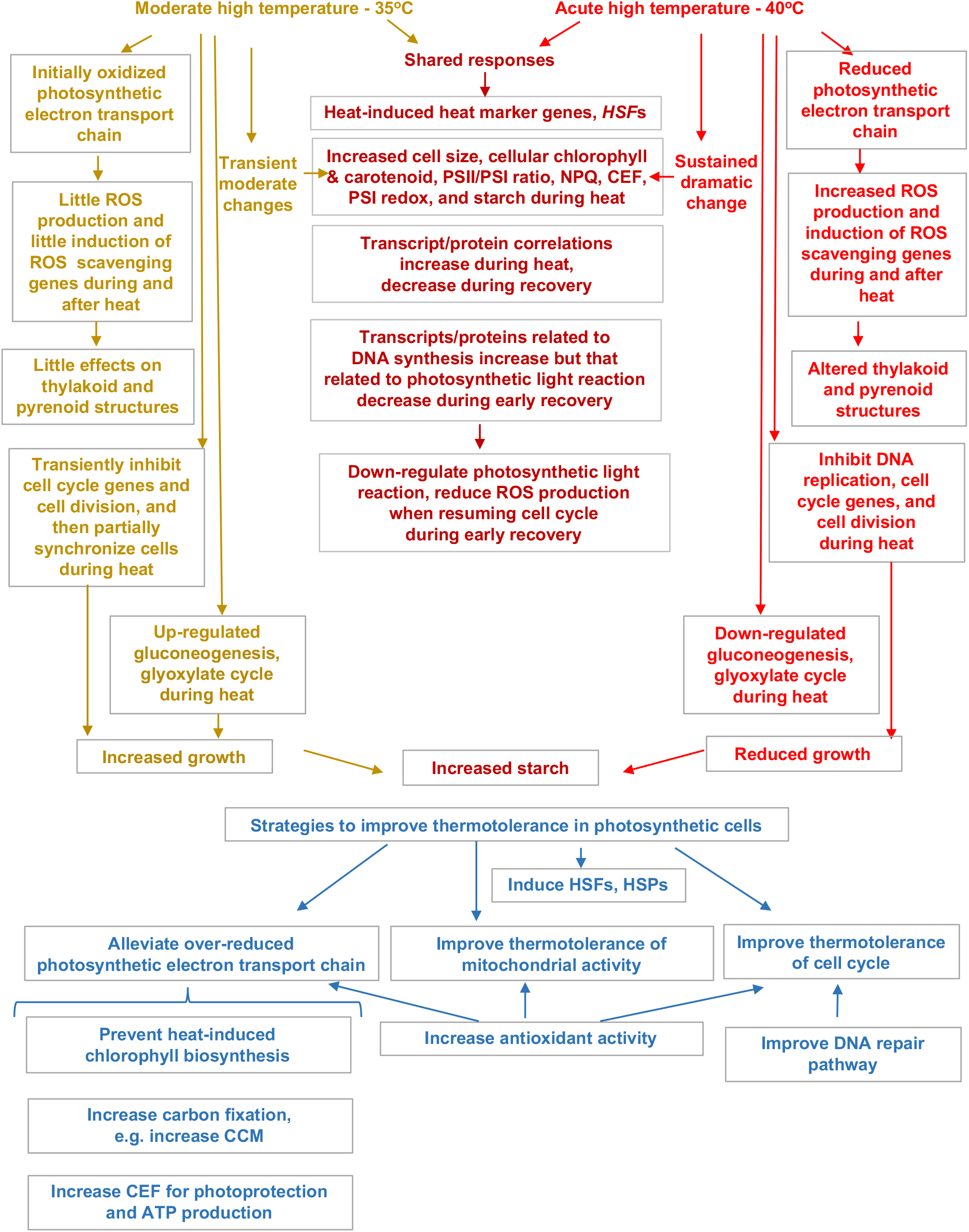
Overview of shared and unique responses to moderate and acute high temperature of 35 and 40°C in Chlamydomonas and strategies to improve thermotolerance in photosynthetic cells.

### Chlamydomonas had shared responses to heat treatments at 35 and 40°C

Both high temperatures induced the expression of *HSF1* and *HSF2*, as well as canonical heat marker genes *HSP22A* and *HSP90A*, increased cell size, chlorophyll and carotenoid contents, PSII/PSI ratio, NPQ, CEF, PSI redox state, and starch formation (Figure 10). The changes under 35°C were often transient and moderate while those under 40°C were sustained and dramatic. The correlation between transcripts and proteins increased during both heat treatments, suggesting that responses during heat were largely transcriptionally regulated. Functional categories of gluconeogenesis/glyoxylate-cycle and abiotic stress had the highest correlation between transcripts and proteins in early time points of 35 and 40°C treatment (Figure 2F), respectively. High correlation values of these functional categories indicate that these responses may occur rapidly without much post-transcriptional regulation, which may help coordinate activities to adapt to high temperature quickly and efficiently. The decreased correlation between transcripts and proteins during both recoveries was consistent with increased protein posttranslational modification after heat treatments based on the RNA-seq network modeling results (Figure 3A, TM8/11).

The increased NPQ during 35 and 40°C heat suggested that both heat treatments compromised photosynthesis under the growth light of 100 µmol photons m^−2^ s^−1^ (Figure 7G, H). Heat at 40°C reduced photosynthetic efficiency much more than 35°C, especially when photosynthetic parameters were evaluated with light intensities higher than the growth light, i.e., 200 and 400 µmol photons m^−2^ s^−1^ (Figure 7,8). Under the growth light we employed, the differences in photosynthetic parameters between 35 and 40°C were smaller with comparable values, consistent with the similar kinetics of transcripts and proteins related to the MapMan bin photosynthetic light reactions (Figure 6). We conclude that in algal cultures grown in PBRs under a growth light of 100 µmol photons m^−2^ s^−1^, both 35 and 40°C heat treatments affected photosynthetic efficiency but photosynthetic activity was maintained at a comparable level during both heat treatments; however, increasing light intensities exaggerated the heat-induced damages to photosynthesis, especially with the 40°C treatment.

During early recovery from both heat treatments, transcripts and proteins related to DNA synthesis increased while those related to photosynthetic light reactions decreased (Figure 3). In synchronized algal cultures under day/night cycles, genes related to DNA synthesis and cell cycle peak during the early dark phase when the genes related to photosynthetic light reactions had minimal expression (Zones et al., 2015; Strenkert et al., 2019). However, under constant light, genes related to DNA synthesis and photosynthetic light reactions may express simultaneously but their quantitative expression manner under constant light is understudied. The induction of cell cycle genes after recovery were comparable after both 35 and 40°C heat (Figure 4A), but the significant down-regulation of genes related to photosynthetic light reactions only occurred in the recovery of 40 but not 35°C (Supplemental Figure 9A), suggesting the uncoupling of these two processes during the recovery of heat treatments may be different from that under day/night cycles. ROS accumulation impairs DNA replication and induces DNA damage (Anderson et al., 2020; Robert and Wagner, 2020). Mammalian cells are more sensitive to high temperatures during DNA replication than other cell cycle stages (Velichko et al., 2013). The photosynthetic light reactions are a major source of ROS production (Hasanuzzaman et al., 2020). Our data showed increased ROS accumulation during the early recovery phase of 40°C but not 35°C. We propose that down-regulation of photosynthetic light reactions during DNA synthesis is beneficial to resuming the cell cycle by reducing ROS production during the early recovery. The mechanisms of the opposite transcriptional regulation of DNA replication and photosynthetic light reactions during the recovery from high temperatures is unknown but interesting for future research. One form of ROS is H_2_O_2_, which is highly diffusible and stable (Hasanuzzaman et al., 2020). Recently, Niemeyer et al. employed hypersensitive H_2_O_2_ sensors in different compartments of Chlamydomonas cells and showed that H_2_O_2_ levels increased in the nucleus after heat treatment, suggesting diffusion of H_2_O_2_ from other cellular compartments to the nucleus (Niemeyer et al., 2021). H_2_O_2_ has been proposed as a secondary messenger in signal transduction (Waszczak et al., 2018). Increased H_2_O_2_ in the nucleus may affect gene expression, e.g., those involved in photosynthetic light reactions.

Both high temperatures affected cell cycle genes, but during 35°C cells could recover the expression of cell cycle genes to the pre-heat level after 8-h of heat treatment with a partially synchronized cell cycle at the end of the 24-h heat treatment (Figure 4). In contrast, under the 40°C heat treatment, the expression of cell cycle genes was abolished. Additionally, we observed down-regulation of genes encoding cell wall proteins during both 35 and 40°C heat treatments and many of these genes were up-regulated during the recovery of 35 and 40°C; the differential regulation of cell wall genes was more dramatic with 40°C than 35°C treatment (Supplemental Figure 9I). The Chlamydomonas cell wall protects cells from environmental challenges; it is proposed that osmotic/mechanical stresses and cell wall integrity regulate the expression of cell wall genes in Chlamydomonas (Cronmiller et al., 2019). However, the underlying mechanisms of how the cell wall responds to high temperatures are largely under-explored. Cell walls form around daughter cells during cell division. Under diurnal regulation, many cell wall genes are up-regulated following the up-regulation of many cell cycle genes during cell division (Zones et al., 2015). The expression pattern of cell wall genes in our data may be related to the inhibited and resumed cell cycles during and after heat treatments, respectively.

### Chlamydomonas responds uniquely to heat treatments of 35 and 40°C

Heat at 35 and 40°C induced unique transcriptional responses (Figure 2, Supplemental Figures 4, 5, 6, 9, 10). Heat at 35°C induced a unique gene set that was not induced under 40°C, including genes involved in gluconeogenesis and glycolysis, mitochondrial assembly, and a putative calcium channel (Supplemental Figure 5C-F, Supplemental Dataset 1, 3). While most of the overlapping DEGs between 35 and 40°C treatments were more strongly differentially expressed at 40°C than at 35°C, a fraction of the overlapping DEGs displayed a larger fold-change at 35°C than at 40°C (Supplemental Figure 6). One group of genes with higher up-regulation at 35°C than at 40°C are those low-CO_2_ inducible (LCI) genes, e.g., *LCI26* (at reaching high temperature and 8-h heat), *LCI19* (16-h heat) (Supplemental Dataset 1), suggesting an effort to compensate for increased CO_2_ demands with increased growth at 35°C. Many of these uniquely regulated genes in the 35°C treatment have unknown functions and may include novel candidates important for acclimation to moderate high temperature.

Both 35 and 40°C induced starch accumulation but possibly for different reasons (Figure 9W, X, 10). The increased starch in 35°C treated cells may be due to increased acetate uptake/assimilation, as evidenced by an induction of gluconeogenesis and the glyoxylate cycle (Figure 2F, Supplemental Figure 8). In Chlamydomonas, acetate uptake feeds into the glyoxylate cycle and gluconeogenesis for starch biosynthesis (Johnson and Alric, 2013, 2012). The increased starch in 40°C treated cells may be due to inhibited cell division (Figures 4, 5), resulting in starch storage exceeding its usage. Starch accumulation could also be an electron sink to alleviate the over-reduced electron transport chain during 40°C heat treatment (Hemme et al., 2014). Heat treated Arabidopsis plants (42°C for 7 h) also had increased starch (Balfagón et al., 2019). Several genes involved in starch biosynthesis were induced during 40°C heat and early recovery (Supplemental Figure 9F). The over-accumulated starch during 40°C may also contribute to the down-regulation of genes involved in acetate uptake and assimilation (Supplemental Figure 9G). Most of the reducing power from acetate assimilation is used in mitochondrial respiration (Johnson and Alric, 2012). However, the down-regulated transcripts related to mitochondrial electron transport (Supplemental Figure 15C), the heat sensitivity of mitochondrial respiration rates (Figure 8H), and the over-accumulated starch may restrict acetate uptake and assimilation during 40°C heat treatment.

Heat at 35°C stimulated growth but 40°C inhibited growth (Figure 1A, B). We quantified growth based on the number of cell divisions and medium consumption in PBRs under the turbidostatic mode, monitored by OD_680_ which is proportional to chlorophyll content and cell density (Supplemental Figure 1A-C). There are different ways to quantify growth. Under our experimental conditions with little nutrient and light limitation, our results showed that cells exposed to 35°C reached the target OD_680_ faster than cell exposed to 40°C or kept at constant 25°C. The stimulated growth in liquid cultures under 35°C was confirmed by growth on plates (Supplemental Figure 2) and it may also be consistent with increased transcripts and proteins related to mitochondrial electron transport as well as increased mitochondrial relative volume in 35°C-treated cells (Supplemental Figure 15). Maize plants grown under moderate high temperature of 33°C had increased biomass but decreased biomass under higher temperature of 37°C as compared to controls at 31°C (Li et al., 2020b), consistent with our data in Chlamydomonas. With increasing high temperatures, the growth of photosynthetic organisms may accelerate to some degree first and then decrease when the high temperature exceeds a certain heat threshold.

### Chlamydomonas employs several strategies to tolerate acute heat of 40°C

The adaptive transcriptional changes in response to 40°C include rapid induction of transcripts encoding HSFs, HSPs, photoprotection, and ROS scavenging enzymes (Supplemental Figure 9A, C, E, 10A). Chlamydomonas has two HSFs, HSF1 and HSF2 (Schroda et al., 2015). HSF1 is a canonical HSF similar to plant class A HSFs and a key regulator of the stress response in Chlamydomonas (Schulz-Raffelt et al., 2007), while the function of HSF2 is unclear. Our transcriptome data showed that both *HSF1* and *HSF2* were induced during early heat of 35 and 40°C (Supplemental Figure 10A), suggesting the potential role of HSF2 in heat regulation. Interestingly, *HSF1* was also induced during the early recovery phase, possibly due to its potential roles in maintaining some heat responsive genes after heat treatment. HSF1 was shown to also be involved in altering chromatin structure for sustained gene expression (Schroda et al., 2015; Strenkert et al., 2011). HSP22E/F are small heat shock proteins targeted to the chloroplast, function in preventing aggregation of unfolded proteins, and are induced at temperatures at or above 39°C in Chlamydomonas (Rütgers et al., 2017b, 2017a). The transcripts of *HSP22E/F* were induced transiently but strongly during 0.5- to 1-h heat of 40°C, but also during the first 4-h of recovery after 40°C heat (Supplemental Figure 9C), suggesting their roles not only during heat but also the recovery from heat.

Additionally, cells treated with 40°C heat had increased photoprotection (Figure 7, 8) and related transcripts were up-regulated (Supplemental Figure 9A, e.g., *LHCSR*, *ELI*, and *PSBS*). With compromised photosynthesis and over-reduced photosynthetic electron transport chain under 40°C, the growth light used in our experiments became excessive and increased ROS production, thus the increased transthylakoid proton motive force (*pmf*), NPQ formation, sensitivity of NPQ to the *pmf*, and CEF activity were all helpful to dissipate excess light energy and reduce heat-induced oxidative stress. CEF generates only ATP but no NADPH, balances the ATP/NADPH ratio, contributes to the generation of *pmf*, and protects both PSI and PSII from photo-oxidative damage (Yamori and Shikanai, 2016; Johnson, 2011). Increased CEF activity has been frequently reported under various stressful conditions in land plants (Zhang and Sharkey, 2009; He et al., 2015; Huang et al., 2012) and algae (Johnson et al., 2014; Saroussi et al., 2016; Aihara et al., 2016). It is proposed that the reduced plastoquinone (PQ) pool activates CEF in algae (Alric et al., 2010; Alric, 2010; Aihara et al., 2016). CEF is also proposed to provide the extra ATP needed for the carbon concentrating mechanisms (CCM) in Chlamydomonas (Lucker and Kramer, 2013). Our results showed that induced CEF at 40°C (Figure 8E) concurred with the increased PQ redox status (Figure 7F), the induced transcripts involved in CCM (Supplemental Figure 9B), and the increased proteins of PSI subunits (Figure 6D).

Cells treated with 40°C heat had increased PSII/PSI ratio measured by 77 K chlorophyll fluorescence (Figure 7I), as previously reported (Hemme et al., 2014). The 77 K chlorophyll fluorescence is often used to monitor the stoichiometries of PSII and PSI (Lamb et al., 2018). The ratios of PSII and PSI from the 77 K fluorescence emission is an indicator of the relative antenna size of each photosystem (Wood et al., 2019). Hemme et al., (2014) reported that 42°C treated Chlamydomonas cells showed a blue shift of PSI emission peak from 713 to 710 nm, suggesting detachment of LHCI from PSI. In 40°C treated Chlamydomonas cells, we also observed the minor blue shift of the PSI emission peak but also an increased emission peak around 695 nm (Supplemental Figure 12), which is associated with the PSII core antenna CP47 (Lamb et al., 2018). Our spectral changes indicated reduced or detached PSI antenna but increased PSII antenna, thus an increased PSII/PSI ratio, suggesting relatively smaller antenna associated with PSI than PSII in cells treated with 40°C heat. Under high salt conditions, the Antarctic alga Chlamydomonas sp. UWO 241 forms a PSI-Cytochrome *b_6_f* supercomplex with constitutively high rates of CEF but absence of a discernible PSI peak in 77 K chlorophyll fluorescence emission (Szyszka-Mroz et al., 2015, 1; Cook et al., 2019; Kalra et al., 2020). Chlamydomonas forms the PSI-Cytochrome *b_6_f* supercomplex to facilitate CEF under anaerobic conditions (Iwai et al., 2010; Steinbeck et al., 2018). Steinbeck et al. (2018) proposed that the dissociation of LHCA2/9 from PSI supported the formation of the PSI-Cytochrome *b_6_f* supercomplex. In Chlamydomonas, the PSI core associates with LHCI which is comprised of ten LHCA subunits and LHCA2/9 are suggested to be weakly bound to the PSI core (Su et al., 2019). The PSI chlorophyll fluorescence under 77 K is mainly due to chlorophyll a in the LHCAs (Lamb et al., 2018). Combining our results with previous reports, we propose that heat-induced dissociation of LHCAs (possibly LHCA2/9) from the PSI core may facilitate the formation of the PSI-Cytochrome *b_6_f* supercomplex and increase CEF activity.

Cells treated with 40°C heat had altered pyrenoid structures (Figure 9 O, P). Algae utilize pyrenoids to concentrate CO_2_ around Rubisco through the CCM (Wunder et al., 2019; Meyer et al., 2017). In Chlamydomonas, pyrenoids consist of three major components: starch sheath (a diffusion barrier to slow CO_2_ escape), pyrenoid matrix (Rubisco enrichment for CO_2_ fixation), and thylakoid tubules (delivery of concentrated CO_2_ and diffusion path of Calvin-Benson Cycle metabolites) (Figure 9S) (Hennacy and Jonikas, 2020). Several pyrenoid-localized proteins sharing a conserved Rubisco-binding motif are proposed to mediate the assembly of the pyrenoid in Chlamydomonas (Meyer et al., 2020): the linker protein Essential Pyrenoid Component 1 (EPYC1) links Rubisco to form the pyrenoid matrix (Mackinder et al., 2016; He et al., 2020); the starch-binding protein Starch Granules Abnormal 1 (SAGA1) mediates interactions between the matrix and the surrounding starch sheath (Itakura et al., 2019); the thylakoid-tubule-localized transmembrane proteins RBMP1/2 mediate Rubisco binding to the thylakoid tubules in the pyrenoid (Meyer et al., 2020). From our TEM images, thylakoid tubules appeared to be absent from the pyrenoid matrix in cells treated with 40°C heat, which may suggest that 40°C heat disrupts the interaction of thylakoid tubules with the pyrenoid matrix and compromises CCM efficiency. The transcripts of *EPYC1*, *SAGA1*, and *RBMP2* were induced during 40°C heat (Supplemental Figure 9B). We propose that 40°C heat may increase the disorder of the pyrenoid structure and Chlamydomonas cells compensate for this by inducing transcripts encoding the pyrenoid-structure-maintaining proteins mentioned above. Several other transcripts related to the CCM, e.g., low CO_2_ inducible proteins, LCIA/D/E/C (helping maintaining CO_2_ concentration in pyrenoids), were all up-regulated during 40°C heat (Supplemental Figure 9B), which may suggest the attempt to maintain the CCM and compensate for the heat induced photorespiration (Hemme et al., 2014) as well as CO_2_ leakage from pyrenoids. *SAGA1* and *LCIA/D/E* were also induced during early recovery, suggesting the efforts to recover the CCM and/or coordinate pyrenoid division with cell division after 40°C heat. The increased CCM transcripts during 40°C heat and early recovery may be an adaptive response to alleviate the over-reduced electron transport chain.

The increased chlorophyll during 40°C heat may be a maladaptive response. Cells treated with 40°C heat had more than 4x increased chlorophyll per cell (Figure 5D), which could not be fully explained by increased cell volume (Supplemental Figure 11A). Increased chlorophyll in heat treated Chlamydomonas cells has been reported previously (Hemme et al., 2014), but the underlying mechanisms are unclear. Heat at 40°C appeared to promote chlorophyll biosynthesis. The gene encoding the key chlorophyll synthesis enzyme, porphobilinogen deaminase, *PBGD2* (Lohr et al., 2005), was up-regulated during 40°C heat (Supplemental Figure 9D). Considering the compromised photosynthesis and inhibited growth during 40°C heat, increasing chlorophyll levels to this extent is toxic. The elevated chlorophyll may lead to increased light harvesting with decreased photosynthesis in 40°C treated cells, resulting in ROS production (Figure 5G). Chlorophyll contents positively correlate with nitrogen availability (Bassi et al., 2018) and we found many genes related to the nitrogen assimilation pathways were up-regulated during 40°C heat (Supplemental Figure 9H), providing a possible explanation for increased chlorophyll during 40°C heat. Maize plants showed greater sensitivity to high temperatures with increased nitrogen fertilization (Ordóñez et al., 2015), which may support the possible links among nitrogen assimilation, chlorophyll biosynthesis, and heat responses. In land plants, long-term (e.g. several days) heat stress reduces chlorophyll content (Wang et al., 2018; Rossi et al., 2017), however, the underlying mechanisms by which chlorophyll is degraded during long-term heat remain elusive (Wang et al., 2018). Chlamydomonas cells treated at 39°C for more than one day had initially increased chlorophyll (8-h heat) followed by chlorophyll loss, cell bleaching, and death (33-h heat) (Zachleder et al., 2019). It is possible that preventing chlorophyll increase during acute high temperature (especially early stage) could lead to improved thermotolerance in algae.

Combing our systems-wide analyses, we could distinguish adaptive versus maladaptive heat responses as mentioned above. The potential engineering targets for improved thermotolerance may include these adaptive heat responses, e.g., heat induced HSFs, HSPs, photoprotection, CEF, antioxidant pathways, and CCM transcripts (Figure 10). The maladaptive responses could also be the targets for improved thermotolerance if we could find mediating solutions to reduce these changes, e.g., heat-induced chlorophyll. The cell cycle arrest induced by acute high temperature may be maladaptive but also adaptive: the halted cell division may be one of the main reasons for over-accumulated starch, reduced photosynthetic electron transport chain, and increased ROS production; on the other side, cell cycle arrest may prevent damages/errors during DNA replication under acute high temperature when DNA repair pathways are compromised by heat (Velichko et al., 2013). Thus, increasing DNA repair pathway and the thermotolerance of cell cycle pathways may also be strategies to improve heat tolerance in photosynthetic cells. The increased gluconeogenesis and glyoxylate cycles may contribute to the increased growth under 35°C heat; however, the faster-growing cell population with increased carbon metabolism probably experience starvation under nutrient limiting conditions thus reducing biomass accumulation and cell growth in relatively short time. Furthermore, mitochondrial activity was stimulated slightly by 35°C heat but sensitive to 40°C heat (Figure 8 H, Supplemental Figure 15), suggesting mitochondrial activity could also be a target to improve thermotolerance. Additionally, the genes associated with TM1 with early heat induction may include pathways that are essential for heat tolerance (Figure 3A, Supplemental Dataset 5). Finally, we compared our algal heat transcriptome with that in Arabidopsis (heat at 42°C for 7 h) (Balfagón et al., 2019) and identified a set of highly conserved heat-induced genes sets (Supplemental Dataset 1), which may provide potential targets to improve heat tolerance in land plants.

In summary, Chlamydomonas is an excellent model to study the heat response and its regulation at the cellular level. Our research helped fill the knowledge gaps regarding how algae respond to and recover from different intensities of high temperatures at multiple levels, discovered the increased transcript/protein correlation during heat treatments, showed the dynamics of photosynthesis in response to high temperatures, and revealed the antagonistic interaction between DNA replication and photosynthetic light reactions during the recovery from both moderate and acute high temperatures. Through systems-wide analyses, we advanced our understanding of algal heat responses and identified engineering targets to improve thermotolerance in green algae and land plants.

## Methods

### Strains and culture conditions

*Chlamydomonas reinhardtii* wildtype strain CC1690 (also called *21gr*, *mt^+^*, from the Chlamydomonas resource center) was used in all experiments. CC1690 were grown in standard Tris-acetate-phosphate (TAP) medium in 400 mL photobioreactors (PBRs) (Photon System Instruments, FMT 150/400-RB). Cultures were illuminated with constant 100 µmol photons m^−2^ s^−1^ light (50% red: 50% blue), mixed by bubbling with filtered air at a flow rate of 1 L/min. After PBR inoculation at initial cell density of 0.5×10^6^ cells/mL, cultures were allowed to grow to a target cell density of 2.00×10^6^ cells/mL corresponding to around 4.0 µg/mL chlorophyll content in log-phase growth at 25°C. Then, the target cell density was maintained turbidostatically using OD_680_ by allowing the culture to grow to 8% above the target cell density before being diluted to 8% below the target cell density with fresh TAP medium provided through peristaltic pumps. Through the turbidostatic mode, the cultures were maintained under constant nutrients and light and had exponential growth between dilution events. The OD_680_ measurement during growth phases in between dilution events was log_2_ transformed, and the growth rate was calculated using the slope of log_2_(OD_680_), while the inverse of the slope yielded the doubling time of the culture (T_d_) (Supplementary Figure 1A, B, C). All algal liquid cultivation used in this paper was conducted in PBRs with the conditions mentioned above.

### High temperature treatments in PBRs

Algal cultures in PBRs were maintained turbidostatically using OD_680_ for 4 days at 25°C to allow cultures to adapt to steady growth conditions before heat treatments. PBR temperatures were then shifted to moderate or acute high temperature conditions (35 or 40°C in different PBRs) for 24 h, then shifted back to 25°C for 48 h for recovery. PBR cultures grown under constant 25°C served as controls. Cultures were maintained turbidostatically during the entire experiment and harvested at different time points for various measurements. Cell density and mean cell diameter were measured using a Coulter Counter (Multisizer 3, Beckman Counter, Brea, CA). For the data in Figure 1A, algal cultures were maintained in PBRs at 25°C for 4 days before switching to 30, 35, or 40°C for 2 days (different temperature switches in separate PBRs). The relative growth rates were calculated at the end of 2-day treatment of each temperature.

### Spotting test for cell viability and growth

Cultures harvested from PBRs were diluted to 2×10^4^ cells mL^−-1^ or 1×10^5^ cells mL^−1^ with TAP medium and 10 µL aliquots of the diluted cultures were spotted on 1.5% TAP agar plates and grown in temperature-controlled incubators under 25 or 35°C with constant white LED light of 150 µmol photons m^−2^ s^−1^ for 44 h or 3 days. After 44-h growth, algal spots with 200 cells were imaged by a dissecting Leica microscopy and were used for growth quantification. Colony number and area were quantified using ImageJ. Viability was calculated as the number of colonies on plates divided by the number of cells spotted. Algal spots with 200 and 1000 cells were imaged after 3-day-growth for visual representations.

### High temperature treatments in water bath

To check the effects of heating speed on cell viability (Supplemental Figure 1F), PBR cultures before heat treatment were incubated in a water bath which was heated from 25 to 41°C in 25 min then kept at 41°C for 2 h (gradual heating), or directly heated in a water bath which was pre-heated to 41°C then kept at 41°C for 2 h (sharp temperature switch). Cell viabilities after 2-h 41°C heat treatment (either gradual or sharp heating) were assayed using the spotting test as above. PBRs cannot switch from the control to high temperatures in less 25 min so a water bath was used for this test, as previously reported (Hemme et al., 2014; Rütgers et al., 2017b).

### RNA extraction and RT-qPCR

At each time point, 2 mL PBR cultures were pelleted with Tween-20 (0.005%, v/v) by centrifugation at 1,100 x g and 4°C for 2 min. The cell pellet was flash frozen in liquid nitrogen and stored at −80°C before processing. Total RNA was extracted with the TRIzol reagent (Thermo Fisher Scientific, Cat No. 15596026) as described before with some modifications (Zones et al., 2015). RNA was purified by RNeasy mini-column (Qiagen, Cat No. 74106) after on column digestion with RNase-free DNase (Qiagen, Cat No. 79256) according to the manufacturer’s instructions. RNA was quantified with Qubit™ RNA BR Assay Kit, (Life technology, Cat No. Q10210). Total 0.4 μg RNA was reverse transcribed with oligo dT primers using SuperScript® III First-Strand Synthesis System (Life technology, Cat No. 18080-051) according to the manufacturer’s instructions. Quantitative real-time PCR (RT-qPCR) analysis was carried out using a CFX384 Real-Time System (C 1000 Touch Thermal Cycler, Bio-Rad, Hercules, California) using SensiFAST SYBR No-ROS kit (Bioline, BIO-98020). PCR was set up as follows: (1) 2 min at 95°C; (2) 40 cycles of 5 s at 95°C, 10 s at 60°C and 15 s at 72°C; (3) final melt curve at 60°C for 60 s, followed by continuous ramping of temperature to 99°C at a rate of 0.5°C s^−1^. Melting curves and qPCR products was checked after PCR cycles to make sure there are no primer dimers or unspecific PCR products. All qPCR products were sequenced to verify their identifies. Expression of G-protein β-subunit-like polypeptide CBLP (Cre06.g278222) (Schloss, 1990) remain stable among all time points, and were used as internal controls (Xie et al., 2013). The relative expressions were calculated relative to its expression in pre-heat by the _ΔΔ_CT method (Livak and Schmittgen, 2001). The qPCR primers for tested genes (*CBLP*, *HSP22A*, *HSP90A*, and others) are listed in Supplementary Table 1. PCR efficiencies were checked and were employed to the 2^−ΔΔCT^ method to calculate the relative expression as described previously (Livak and Schmittgen, 2001; Hellemans et al., 2007; Remans et al., 2014). Three biological replicates for each time point and treatment were conducted.

### Transcriptomics

RNA libraries were prepared and sequenced by the Joint Genome Institute (JGI, Community Science Program) using the NovaSeq platform and generated 150-nt paired-end reads, with the goal of 20 million genome-mappable reads per sample. Samples were quality control filtered using the JGI BBDuk and BBMap pipelines (Bushnell). BBDuk trims adapter sequences, and quality trims reads with low complexity, low quality scores, and reads with 1 or more “N” bases. BBMap removes reads that map to common contaminant genomes. Samples were quality assessed using FastQC (Andrews, 2010) and mapped to the *Chlamydomonas reinhardtii* v5.6 genome (Merchant et al., 2007) using STAR (Dobin and Gingeras, 2015; Dobin et al., 2013) with the maximum number of mapped loci set to 1 and the maximum mismatches per read set to 1, resulting in >92% of all reads being uniquely mapped to the *Chlamydomonas reinhardtii* v5.6 genome. Reads per feature were counted via HT-seq count (Anders et al., 2014). The dataset was filtered for genes that met minimum read count cutoffs of at least 10 mapped reads in at least 10% of the samples, resulting in 15,541 genes for downstream analyses. Two-dimensional Uniform Manifold Approximation and Projection (UMAP) was used to reveal clusters of RNA-seq data (Ma et al., 2021).

Differential expression modeling was performed on transcripts <u>p</u>er <u>m</u>illion (TPM) normalized read counts using a generalized linear mixed-effect model and a negative binomial distribution. Treatment time points were compared to pre-heat controls. Significant differential expression was defined as |log_2_ fold-change > 1, FDR < 0.05, and |(control mean TPM) – (treatment mean TPM)| ≥ 1. FDR correction was performed using the Benjamini-Hochberg method (Benjamini and Hochberg, 1995). Heatmaps were generated using the R package pheatmap (version 1.0.12. https://CRAN.R-project.org/package=pheatmap). Weighted correlation network analysis (WGCNA) was performed on TPM normalized read counts that met minimum read count cutoffs (Langfelder and Horvath, 2008). Expression patterns from 35°C and 40°C were modeled together as a signed network, requiring at least 50 genes per module and combining modules with similarity >0.25. All genes were tested against all eigengene modules using ANOVA (FDR < 0.05) and were assigned to the module with highest significance. Functional enrichment analysis was performed using hypergeometric tests with subsequent FDR control based on MapMan annotations for *Chlamydomonas reinhardtii* (FDR < 0.05) (Benjamini and Hochberg, 1995). Surprisal analysis was applied to dissect the transcriptional regulatory response into its major components (Remacle et al., 2010). Constraint potentials were calculated for each transcriptional data matrix, whose signals determine the deviations of the transcriptome from a theoretical balance state of minimal free energy (Schneider et al., 2020). The overall importance of the transcriptional pattern was estimated from its amplitude and decreases with increasing constraint index. Accordingly, the three major constrains were kept.

### Proteomics

Algal samples (2 mL) harvested from PBRs were centrifuged to remove supernatant, flash frozen in liquid nitrogen, and stored at −80°C until use. Proteins were extracted using the IST sample preparation kit (PreOmics GmbH, Germany). Cell pellets (about 200 µg protein) were lysed in 50 µL lysis buffer, and 1/5^th^ of the lysate was added into 40 µL lysis buffer and sheared in a sonicator (VWR Aquasonic 250D, 35Khz) for 3 min, then digested in a heating block at 37°C (500 rpm, 3 h) followed by stopping the digestion with the stop buffer. Peptides were purified using the PreOmics cartridges according to the manufacturer’s instruction. Eluted peptides were transferred to 96 well plates, dried under vacuum for 2 h, and then dissolved in 30 µL of LC-loading buffer included in the kit. Finally, 5 µL of the suspension was used for LC-MS/MS analysis. LC-MS/MS was carried out on an Orbitrap Fusion Lumos (Thermo Fisher Scientific, San Jose, CA) mass spectrometer coupled with a U3000 RSLCnano HPLC (Thermo Fisher Scientific, San Jose, CA). The peptide separation was carried out on a C18 column (Fritted Glass Column, 25 cm × 75 μm, Reprosil-Pur 120 C18-AQ, 1.9 μm, made by ESI Source Solution, LLC., Woburn, MA) at a flow rate of 0.3 μL/min and the following gradient: Time = 0–4 min, 2% B isocratic; 4–8 min, 2–10% B; 8–83 min, 10–25% B; 83–97 min, 25–50% B; 97–105 min, 50–98%. Mobile phase consisted of A, 0.1% formic acid; mobile phase B, 0.1% formic acid in acetonitrile. The instrument was operated in the data-dependent acquisition mode in which each MS1 scan was followed by Higher-energy collisional dissociation (HCD) of as many precursor ions in 2 second cycle (Top Speed method). The mass range for the MS1 done using the FTMS was 365 to 1800 m/z with resolving power set to 60,000 @ 200 m/z and the automatic gain control (AGC) target set to 1,000,000 ions with a maximum fill time of 100 ms. The selected precursors were fragmented in the ion trap using an isolation window of 1.5 m/z, an AGC target value of 10,000 ions, a maximum fill time of 100 ms, a normalized collision energy of 35 and activation time of 30 ms. Dynamic exclusion was performed with a repeat count of 1, exclusion duration of 30 s, and a minimum MS ion count for triggering MS/MS set to 5000 counts.

#### Protein identification and quantification

Quantitative analysis of MS measurements was performed using MaxQuant 1.6.12.0 (Cox and Mann, 2008). The library used to perform peptide spectrum matching was created based on version JGI5.5 of the C. reinhardtii genome. The search space was extended including methionine oxidation and acetylation of protein N-termini as variable modifications. The false discovery rate (FDR) thresholds for peptide spectrum matching and protein identification were set to 1%. Protein abundances were estimated using the Label free Quantification (LFQ) algorithm (Cox et al., 2014).

#### Data normalization and protein-level missing value imputation

Normalization and missing value imputation were performed independently for 35°C and 40°C time courses. Proteins were excluded from the data set if there was no biological replicate group with more than one value among different time points, as these proteins were considered not suitable for quantitative downstream analysis. Following normalization using the median-of-ratios method (Anders and Huber, 2010), we used global variance estimates and local gene wise mean estimates to impute missing data points as independent draws from normal distributions. If a protein showed no quantification in a group of biological replicates at one time point, it was checked if the protein was present in the adjacent time points (the time point before and after the one in query). If this was the case, the protein mean was imputed using k-Nearest-Neighbour imputation, followed by sampling from a normal distribution. If adjacent time points had no values, no imputation was performed for the time point in query. All further protein analyses were based on imputed values.

#### Network generation

All protein groups resulting from non-proteotypic peptides were duplicated to singletons and the intensities of each protein were log-transformed. To ignore proteins with constant abundance signals, they were filtered for significance by one-way ANOVA (p<0.05). To generate a correlation matrix, the Pearson correlation coefficient was used. An absolute correlation threshold was determined by random matrix theory (Luo et al., 2007). These thresholds were determined to be ρ=0.8194 and ρ=0.8675 for 35 and 40°C respectively. After filtering the correlation matrix accordingly, the nodes and edges were isolated and visualized in Gephi (www.gephi.org) using ForceAtlas2 (Jacomy et al., 2014) and built-in modularity determination (Blondel et al., 2008).

#### Statistical testing of proteins

Proteins that were identified by ambiguous (non-proteotypic) peptides or had missing values in at least one replicate were not considered for statistical testing. All time points were tested for significant accumulation or depletion in respect to a control measured prior to the start of the heat treatment. Dunnett’s multiple-comparison test was applied with alpha levels of 0.05 and 0.01.

#### Proteomics enrichment analyses

To investigate the module compositions a term enrichment based on MapMan ontology was performed. The ontology tree of each term was expanded, so every protein could exist multiple times, corresponding to the number of ontology term levels. A hypergeometric test for each functional term was applied with subsequent FDR control by the Benjamini-Hochberg method (Benjamini and Hochberg, 1995). To derive a representative signal shape of all proteins included in a module, its eigenvector was calculated based on complete time series, which were log-transformed and centered to an intensity of zero mean and unit variance (Langfelder and Horvath, 2007). For negatively correlating proteins, its corresponding eigenvector was inverted. Additional term enrichments were applied following the previous schema to gain insights into functions of positively or negatively correlating proteins of each module.

#### Correlation of transcripts and proteins

The log_2_(fold-change) of transcript reads and protein abundance were calculated in respect to the pre-heat samples. All available transcript/protein pairs were taken into consideration for correlation analysis. The heat period (HS) as well as the recovery period (RE) were split up into three windows each (HS: 0-1 h, 2-8 h, 16-24 h during the heat period; RE: 0-2 h, 4-8 h, 24-48 h during the recovery period after heat treatment). The average log_2_(fold-change) is determined for each window. Every identifier, that had both a transcript and protein associated with it, resulted in a transcript-protein fold-change pair for each window. By collecting all identifiers that are associated with a respective functional term a scatter plot of transcript-protein fold-change pairs were generated, and the Pearson correlation coefficient was calculated. By collecting the correlation coefficient for every present functional term, a density plot for each of the six windows was created. The resulting correlation coefficient histograms were smoothed using Silvermans rule of thumb for kernel density estimation (Silverman, 1986). See Supplemental Figure 7 for the illustration.

All computational analyses on protein intensities were conducted using the open-source F# libraries FSharp.Stats, BioFSharp, and Plotly.NET. Linear regression, Benjamini– Hochberg correction, smoothing, correlation measures, and eigenvector calculations were performed using the FSharp.Stats version 0.4.1-beta. For ontology annotation and GSEA based on hypergeometric tests, we used BioFSharp version 2.0.0-beta4. Visualization of transcript-protein correlation was performed using Plotly.NET version 2.0.0-alpha5.

### DNA content and ploidy

DNA content was analyzed by FACS (fluorescence-activated cell sorting) with modified protocol (Tulin and Cross, 2014). Cell cultures (10 mL) were collected and fixed in 30 mL ethanol:acetic acid (3:1) for 15 min at room temperature. Cells were spun down at room temperature at 4000 x g for 1 min and washed once with 1 mL phosphate-buffered saline (PBS). Then cells were collected by centrifugation, resuspended in 2 mL PBS with RNAse A (100 µg/mL) for 2 h at 37°C, centrifuged again and finally resuspended in 2 mL PBS + 500 nM Sytox Green (Thermo Fisher Scientific, S7020). FACS was performed on an Accuri C6 instrument (BD Biosciences, Franklin Lakes, NJ), reading 20,000 cells per sample in the FL1 channel (488 nm exciting laser; emission filter: 530 ± 15 nm). A 90% attenuator was used to reduce signal below saturation levels. Data were analyzed using FlowJo software (BD Biosciences). Assignment of cell populations representing 1C, or >1C DNA content was determined. The raw fluorescence signal of 2C was not two-times the signal at 1C because there was background staining, which was cell-size-dependent (Tulin and Cross, 2014). Therefore, high-ploidy cells (which did not get much bigger during a multiple fission cycle) had DNA signal that was about proportional to actual DNA contents, while in low-ploidy cells the background contributed more. Thus, in the population of pre-heat samples, background is about 0.5 x10^5^ of the DNA content fluorescence signal (x axis). Before background subtraction, 1C, 2C, 4C cells peaked at 3 x10^5^, 5.5 x10^5^, 10 x10^5^ of DNA content fluorescence signal, respectively. After background subtraction, 1C, 2C, 4C cells were at 2.5 x10^5^, 5 x10^5^, 9.5 x10^5^, respectively. There was also a small peak at 20 x10^5^ which was 8C. At 24 h heat of 40°C, cell size was about 2x bigger (Figure 5A-C), with background of around 2 x10^5^, causing the right shift of the 1C peak (Figure 4K).

### Cell imaging using light microscopy

Cultures harvested at select time points were fixed with 0.2% glutaraldehyde (VWR, Cat No. 76177-346). Cells were imaged with a Leica DMI6000 B microscope and a 63x (NA1.4) oil-immersion objective. Images shown are representative of results from at least three independent experiments (Figure 5A).

### Pigment analysis

Three biological replicates (each with two technical replicates) of 1mL of PBR cultures were harvested at different time points, mixed with 2.5 μL 2% Tween20 (Sigma, P9416-100ML) to help cell pelleting, centrifuged at 14,000 rpm at 4°C, and stored in −80°C after removal of supernatant. Cell pellets were later thawed, resuspended in 1mL of HPLC grade methanol (100%, Sigma, 34860-4L-R), vortexed for 1min, incubated in the dark for 5min at 4°C, and centrifuged at 21,000 rpm at 4°C for 5min. Supernatant containing pigments was analyzed at 470, 652, 665nm in a spectrophotometer (IMPLEN Nonophotometer P300) for carotenoids and chlorophyll a/b concentrations in μg mL^−1^ using the following equations: Chl a + Chl b = 22.12*A_652_ + 2.71*A_665_, Chl a = 16.29*A665 – 8.54*A652, and Chl b = 30.66*A652 – 13.58*A665 (Porra et al., 1989), and carotenoids = (1000*A470 – 2.86*Chl a – 129.2*Chl b)/221 (Wellburn, 1994).

### Protein concentration and ROS determination

Frozen sample pellets were thawed on ice and lysed by sonication in 10 mM Tris-HCl buffer (Joo et al., 2005). Protein concentrations were determined by the method of Lowry (Lowry et al. 1951). The level of ROS was determined by the method described before with some modifications (Pérez-Pérez et al., 2012). A 40 μg total protein extract was used for ROS measurement. Each sample was aliquoted and one of them was added with ascorbate (Thomas Scientific LLC, C988F55) to a final concentration of 100 mM. The samples containing ascorbate were used as background signal and subtracted from each experimental value later. The samples were incubated for 15 min at 25°C. The ROS indicator CM-H2DCFDA (Life Technologies, C6827) dissolved in 20% (v/v) DMSO was then added to a final concentration of 10 μM and incubated for 30 min at 30°C. The samples were transferred to 96 well microplates and ROS-related fluorescence was measured using Tecan Microplate reader M200 PRO with excitation at 485 nm and emission at 525 nm. The results were obtained from three biological replicates. Relative fold-change of ROS signal (compared to pre-heat) was either normalized to cell number or cell volume. Each of the three biological replicates included two independent measurements with two technical replicates. The ROS signal was normalized to cell number or cell volume.

### Spectroscopic measurement of photosynthetic parameters

Photosynthetic measurements (chlorophyll fluorescence, electrochromic shift, and P700) were conducted using a multi-wavelength kinetic spectrophotometer/fluorometer with a stirring enabled cuvette holder (standard 1 cm pathlength) designed and assembled by the laboratory of Dr. David Kramer at Michigan State University using the method described before with some modifications (Lucker and Kramer, 2013). A 2.5 mL volume of algal cells were sampled from the photobioreactors, supplemented with 25 µL of fresh 0.5 M NaHCO_3_, loaded into a fluorometry cuvette (C0918, Sigma-Aldrich), and dark-adapted with a 10-min exposure to far-red light with peak emission of 730 nm at ∼35 µmol photons m^−2^ s^−1^. Maximum efficiency of PSII (F_v_/F_m_) was measured with the application of a saturating pulse of actinic light with peak emission of 625 nm at the end of the dark adaptation period and after turning off the far-red light. Our tests indicated that far-red illumination could increase the values of F_v_/F_m_ in dark-adapted algal cells. Far-red light was turned off during all chlorophyll fluorescence measurements to prevent its effect on chlorophyll fluorescence signals. Fluorescence measurements were taken with measuring pulses of 100 µs duration. The pulsed measuring beam was provided by a 505 nm peak emission LED filtered through a BG18 (Edmund Optics) color glass filter. After dark-adaptation, the algal sample was illuminated by a pair of light emitting diodes (LEDs) (Luxeon III LXHL-PD09, Philips) with maximal emission at 620 nm, directed toward both sides of the cuvette, perpendicular to the measuring beam. Subsequent chlorophyll fluorescence and dark interval relaxation kinetic (DIRK) measurements were taken after 7.5-min adaptation of sequentially increasing light intensities of 100, 200, and 400 µmol photons m^−2^ s^−1^. DIRK traces measure the electrochromic shift (ECS). ECS results from light-dark-transition induced electric field effects on carotenoid absorbance bands (Witt, 1979; Baker et al., 2007) and is a useful tool to monitor proton fluxes and the transthylakoid proton motive force (*pmf*) *in vivo* (Kramer et al., 2004; Cruz, 2004). ECS measurements were taken at each light intensity following the chlorophyll fluorescence measurements. DIRK traces were run with measuring beam of peak 520 nm and pulse duration of 14 µs capturing absorption changes from a light-dark-light cycle with each

#### P700 measurement

Algal cultures were sampled directly from PBRs at different time points with different treatments, supplemented with 0.5 M NaHCO_3_, incubated with 10 µM DCMU (3-(3,4-Dichlorophenyl)-1,1-dimethylurea, Sigma, D2425) to block PSII activity, and dark adapted for 5 min before the measurement. PSI redox kinetics were monitored using a 703 nm measuring beam pulsed at 10-ms intervals using the spectrophotometer mentioned above as described previously with some modifications (Lucker and Kramer, 2013). After a 5 s baseline measurement in the dark, samples were exposed to 720 nm far-red light at an intensity of 30 µmol photons m^−2^ s^−1^ for 5 s to preferentially oxidize PSI before monitoring the reduction of oxidized P700 (P700^+^) in dark for 5 s. Addition of saturating flash at the end of far-red illumination did not change the amplitude of P700^+^ and the reduction time constant of P700^+^ significantly so the saturating flash was skipped in the finalized P700 measurement. Five measurements were averaged into one trace, and the reduction time constant of PSI was calculated by fitting to a first order exponential function to the reduction phase of the averaged trace. The reduction time constant of P700^+^ in the absence of PSII activity (with DCMU) can be used to estimate the activity of cyclic electron flow around PSI (CEF) (Alric, 2010).

#### 77 K chlorophyll fluorescence measurement

Chlorophyll fluorescence emission spectra were monitored at 77 K to estimate antenna sizes (Lamb et al., 2018; Murakami, 1997). Sampled algal cultures from PBRs were immediately loaded into NMR tubes (VWR, cat. No. 16004-860) and frozen in liquid nitrogen. While still submerged in liquid nitrogen, frozen samples were exposed to excitation LED light with peak emission at 430 nm through a bifurcated fiber optic cable coupled to an LED light source (Lucker and Kramer, 2013). Components were held in alignment using a 3D printed device. Chlorophyll emission spectra were recorded on an Ocean Optics spectrometer (cat. No. OCEAN-HDX-XR), and three consecutive traces were averaged into one measurement. Further dilution of the PBR samples gave identical fluorescence emission peak distributions, indicating little distortion of the signals by reabsorption. Spectral data were normalized to the PSII spectral maximum value at 686 nm, and the relative percentage of PSII was calculated using the normalized PSII peak and PSI peak (714 nm) using this formula PSII%= PSII peak / (PSII peak + PSI peak).

See Supplemental Table 2 for formulas of presented photosynthetic parameters. All measurements were taken with at least three biological replicates. Statistical significance was assessed using a two-tailed t-test of unequal variance. For more than 20 comparisons, FDR using Benjamin-Hochberg correction method was performed with adjusted p value < 0.05 as significance.

### Oxygen evolution measurements

Oxygen evolution was measured with Hansatech Chlorolab 2 based on a Clark-type oxygen electrode, following the method described before with some modifications (Jeong et al., 2017). Two-mL of cells supplemented with 20 μL of 0.5 M NaHCO_3_ were incubated in the dark for 10 min. The dark respiration rate was measured at the end of the dark incubation. The rate of oxygen evolution was measured at increasing light intensities (100, 200 and 400 μmol photons m^−2^ s^−1^). Each light intensity lasted 5 min and the rates of oxygen evolution at each light intensity step were recorded for 1 min before the end of each light phase. The results were obtained from independent measurements of five biological replicates. Statistical analyses were performed using two-tailed t-test assuming unequal variance by comparing treated samples with the pre-heat samples under the same light.

### Transmission Electron Microscopy (TEM)

Algal samples (10 mL, around 2×10^6^ cells/ml) harvested from PBRs were concentrated (1000 g, 2 min, room temperature), drawn into dialysis tubing (Spectrapor®, Spectrum Laboratories, Inc., Cat No. 132294), immersed in 20% (w/v) Bovine Serum Albumin (BSA. Sigma, Cat No. A7030-100G) and immediately frozen in a high-pressure freezer (Leica EM ICE; Leica Microsystems, IL, USA). Freeze substitution was carried out in 2% (v/v) osmium tetroxide in 97%/5% acetone/water (v/v) as follows: −80°C for 3 days, −20°C for 1 day, 4°C for 2 h, and then room temp for 2 h. Samples were washed 4 times in acetone, incubated in 1% thiocarbohydrazide (TCH) (EM Sciences, 21900) in acetone for 1 h, washed 4 times in acetone, incubated for 1 h in 2% (v/v) osmium tetroxide and washed 4 times in acetone. Samples were then laced in saturated uranyl acetate in 100% acetone overnight, washed in acetone, transferred to 1:1 ethanol:acetone and stained with saturated lead acetate in 1:1 ethanol: acetone for 2 h. Subsequently, cells were then dehydrated in 100% graded acetone and embedded in hard formulation Epon-Araldite (Embed 812, EM Sciences, Cat No. 14121). Embedments were cut into small blocks and mounted in the vise-chuck of a Reichert Ultracut UCT ultramicrotome (Leica, Buffalo Grove, IL, USA). Ultrathin sections (∼60 to 70 nm) were cut using a diamond knife (type ultra 35°C, Diatome), mounted on copper grids (FCFT300-CU-50, VWR, Radnor, PA, USA), and counterstained with lead citrate for 8 min (Reynolds, 1963).

Sample were imaged with a LEO 912 AB Energy Filter Transmission Electron Microscope (Zeiss, Oberkochen, Germany). Micrographs were acquired with iTEM software (ver. 5.2) (Olympus Soft Imaging Solutions GmbH, Germany) with a TRS 2048 x 2048k slow-scan charge-coupled device (CCD) camera (TRÖNDLE Restlichtverstärkersysteme, Germany). Thirty electron micrographs were quantified for each time point and treatment. Each TEM image was acquired at 8,000X magnification and 1.37 nm pixel resolution. TEM images were analyzed with Stereology Analyzer software version 4.3.3 to quantify the relative volume mitochondria. Grid type was set as “point” with a sampling step of 500×500 pixels and pattern size of 15×15 pixels. The percent of relative volume for mitochondria was collected after identifying all grid points within one cell and further analyzed in excel. TEM images with a magnification of 8K were used in the Fiji (ImageJ, FIJI software, National Institutes of Health) analysis. The images were scaled to 0.7299 pixel/nm in ImageJ before the analysis of the pyrenoid area. Two different statistical tests were used to find the significance of p-values. The Kolmogorov-Smirnov test was used for relative volume data since it is commonly used to find significance between data in a form of ratios. A two-tailed t-test with unequal variance was used for area data from ImageJ. All statistical tests compared the treatment conditions to the pre-heat.

### Starch Quantification

Starch was extracted and quantified according to modified method from kit (Megazyme, K-TSTA-100A). Frozen sample pellets were homogenized using Qiagen TissueLyser and washed twice in 80% (v/v) ethanol at 85°C. The insoluble fraction was resuspended in DMSO and heated at 110°C for 10 min to improve starch solubilization. Starch hydrolysis and quantification were performed following kit protocol. Starch content was either normalized to cell number or cell volume. Each condition has three biological replicates.

### Data availability

The datasets analyzed in this paper are included in this published article and supplementary information files. The RNA-seq data discussed in this publication have been deposited in the NCBI Gene Expression Omnibus (Edgar et al., 2002) and are accessible through GEO accession number GSE182207. The mass spectrometric proteomic data is available via the ProteomeXchange Consortium partner repository, PRIDE (Perez-Riverol et al., 2019; Deutsch et al., 2020), with the dataset identifier PXD027778. Other information is available from the corresponding author on request.

## Supporting information

supplementary_dataset_1_RNAseq_overview

supplementary_dataset_2_proteomics_overview

supplementary_dataset_3_MapManenrichment

upplementary_dataset_4_PearsonCorrelation

supplementary_dataset_5_WGCNA

supplementary_dataset_6_genes_used_for_heatmaps

supplementary_dataset_7_proteomics_network

supplementary_dataset_8_zScores_transcripts_proteins

8supplementary_dataset9_transcript_protein_correlation

upplementary_dataset10_transcript_protein_kinetics

Fig9_TEM_high_resolution

## Supplemental Datasets

**Supplemental Dataset 1:** Transcriptomics overview. Transcripts per million (TPM) normalized read counts for all genes in all samples, differential expression output data for all genes in all time points, summary of overlapping differentially expressed genes between treatments and time points, genes uniquely up-regulated in 35°C heat, genes more highly differentially expressed in 35°C than 40°C, comparison between this high temperature dataset and the recent work in Arabidopsis (Balfagón et al., 2019).

**Supplemental Dataset 2:** Proteomics overview. Normalized protein spectral counts for all proteins identified in each sample, differential accumulation output data for all proteins in all time points, and summary of overlapping differentially accumulated proteins between 35 and 40°C treatment groups.

**Supplemental Dataset 3:** MapMan functional enrichment for uniquely and overlappingly differentially expressed genes between 35 and 40°C at each time point.

**Supplemental Dataset 4:** Pearson Correlation Coefficients for transcript-protein pairs in individual MapMan functional categories.

**Supplemental Dataset 5:** WGCNA output, genes belonging to each module and functional enrichment of each module, and summary of genes with unknown functions in each module.

**Supplemental Dataset 6:** Genes used to generate heatmaps for pathways of interest.

**Supplemental Dataset 7:** Proteomics network modeling output, proteins belonging to each module and functional enrichment of each module.

**Supplemental Dataset 8**: Z-scores for transcripts and proteins with 35 and 40°C treatment and their MapMan functional categories.

**Supplemental Dataset 9: Transcript and protein correlation plots grouped by MapMan function bins.** The heat treatment period (HS) and the recovery period (RE) were split up into three windows each (HS1-3: 0-1 h, 2-8 h, 16-24 h during the heat period; RE1-3: 0-2 h, 4-8 h, 24-48 h during the recovery period after heat treatment, labelled at the bottom of the figures). The average log_2_(fold changes), lfc, of transcripts and protein pairs in respect to the pre-heat were determined for the three heat (HS, top panels) and three recovery windows (RE, bottom panels), respectively. X and Y, transcript and protein lfc compared with the pre-heat, respectively. All transcript-protein pairs are shown as gray dots. Best fit lines for all transcript-protein pairs are shown in blue and Pearson correlation coefficient are shown on the right in the order for each panel. Transcript-protein pairs belonging to the indicated MapMan functional category (labeled on top of scatterplots) are shown in orange. Each file is .html format and can be opened as an interactive figure with gene IDs and annotations in a web-browser. The figures can be viewed in detail by clicking on “Show closest data on hover” in the upper right corner of the .html file (Click the upper right corner to see the hidden button).

**Supplemental Dataset 10: Transcript and protein kinetics grouped by MapMan function bins.** Transcript and protein signals related to each MapMan function bin were standardized to z scores (standardized to zero mean and unit variance) and are plotted against equally spaced time point increments as labeled on the x axis. Time points (TP) are labeled at the bottom. Time point 1: pre-heat. Time points 2-9, heat treatment at 35 or 40°C, including reaching high temperature, 0.5, 1, 2, 4, 8, 16, 24 h during heat; time points 10-15, recovery phase after heat treatment, including reaching control temperature, 2, 4, 8, 24, 48 h during recovery. Each file is .html format and can be opened as an interactive figure with gene IDs and annotations in a web-browser. The curves can be viewed in detail by clicking on “Show closest data on hover” in the upper right corner of the .html file (Click the upper right corner to see the hidden button).

---

**Supplemental Table 1:**
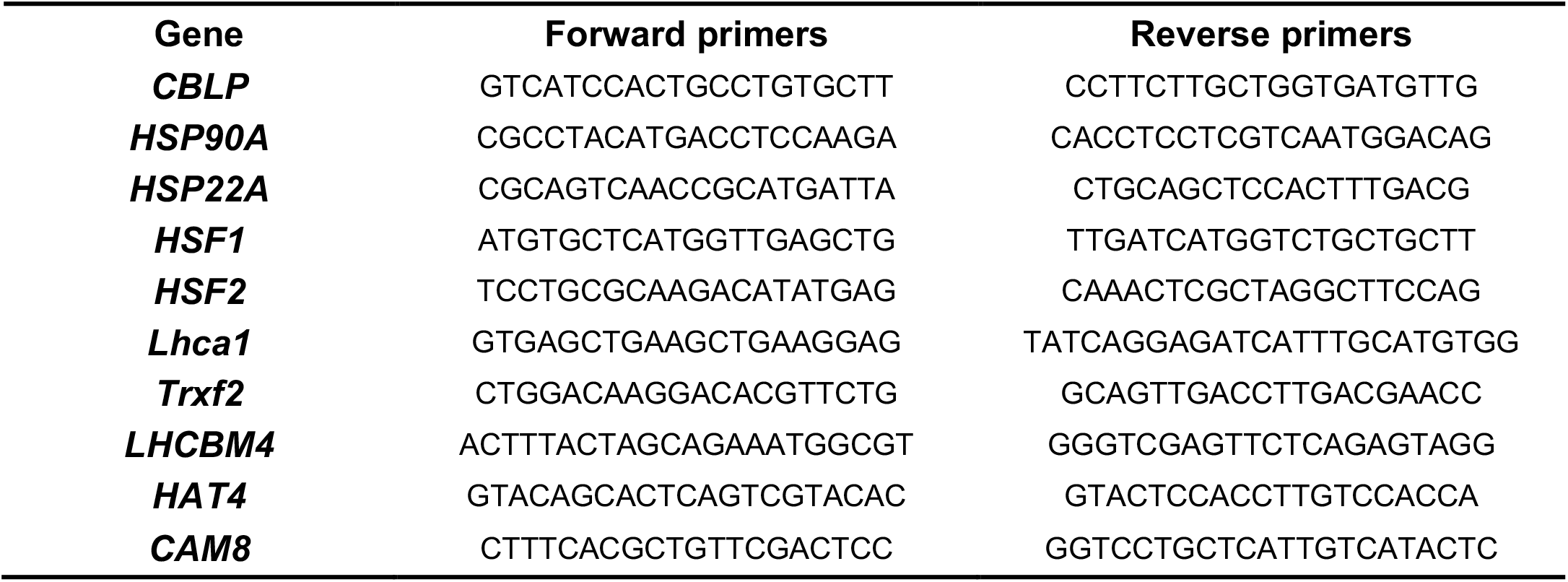
Primers used for qPCR.

**Supplemental Table 2:**
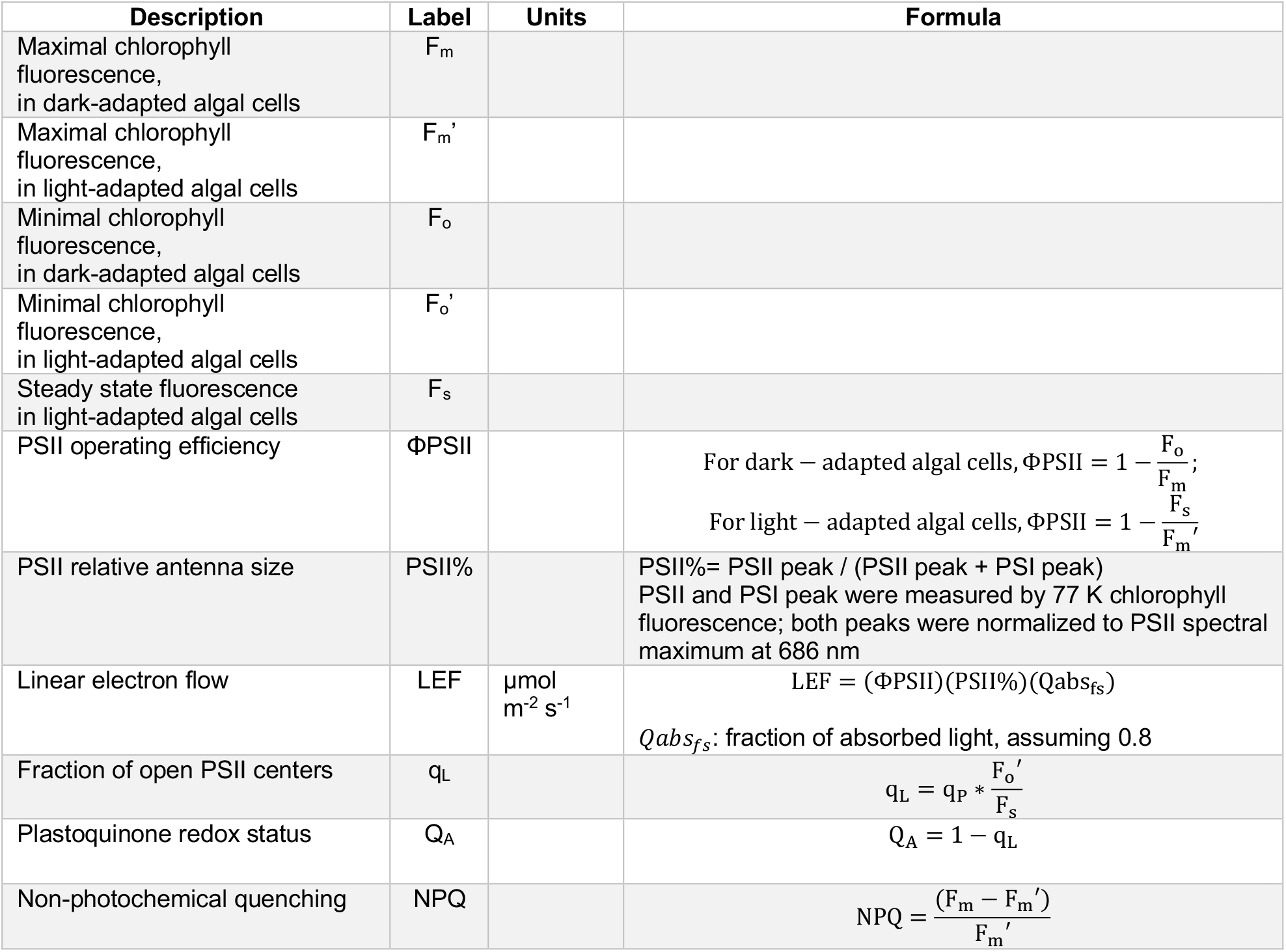
Formulas used to calculate photosynthetic parameters.

## Acknowledgments

This research was supported by the start-up funding from the Donald Danforth Plant Science Center (DDPSC), the Department of Energy (DOE) Basic Energy Sciences (BES) Photosynthetic Systems (PS) grant (Award No. 0019464), the DOE Biological & Environmental Research (BER) grant (Award No. 0020400), and the RNA-seq service support from the Department of Energy (DOE) Joint Genome Institute (JGI) Community Science Program (CSP, 503414) to R.Z.. The work conducted by the U.S. Department of Energy Joint Genome Institute is supported by the Office of Science of the U.S. Department of Energy under Contract No DE-AC02-05CH11231. E.M. was supported by the William H. Danforth Fellowship in Plant Sciences and Washington University in St. Louis. M.S. was funded by Deutsche Forschungsgemeinschaft (TRR175, projects C02 and D02). We would like to thank researchers from the Kramer Laboratory (Drs. David Kramer, Jeffrey Cruz, Robert Zegarac, Ben Lucker, Geoffry Davis) for their helpful suggestions and discussion to set up and optimize the IDEAspec for spectroscopic measurements in algal cells. Dr. Geoffry Davis is also thanked for his suggestions for 77 K chlorophyll fluorescence measurement. We would like to thank Drs. Jeremy Schmutz, Susannah Tringe, Christa Pennacchio, Chris Daum, and Ronan O’Malley for RNA-seq service at DOE JGI. We acknowledge imaging support from the Advanced Bioimaging Laboratory (RRID:SCR_018951) at DDPSC and the usage of the LEO 912AB Energy Filter TEM acquired through a National Science Foundation (NSF) Major Research Instrumentation grant (DBI-0116650). We also acknowledge proteomics support from the Proteomics and Mass Spectrometry at DDPSC and usage of the Orbitrap Fusion Lumos LC-MS/MS, which was funded by the National Science Foundation under Grant No. DBI-1827534. We would like to thank Dr. James Umen for helpful discussion about our research and Sarah Rommelfanger for suggestions on RNA-seq analysis. We also want to thank Drs. Xin Wang and Blake Meyers for valuable feedback about the manuscript.

## Author contributions

R.Z. supervised the whole project. R.Z. and N.Z. designed and planned all the experiments. R.Z. and M.S. discussed and designed the time points for high temperature treatments. N.Z. led the project, characterized cell physiologies (including cell density, size, viability, protein, ROS, starch, light microscopic images), conducted RT-qPCRs and extracted RNA for RNA-seq analysis. W.M. and M.X. grew and maintained algal cultures in photobioreactors. N.Z, W.M., C.A. J.J, M.X. and E.M. harvested algal samples from photobioreactors with different treatments at different time points. E.M. led RNA-seq data analysis and generated all related figures. J.C.B. provided suggestions for RNA-seq analysis and statistical analysis. N.Z. extracted protein for proteomics. S.T. and B.E. quantified protein abundance using LC-MS/MS. B.V., D. Z., T.M., E.M. and N.Z. analyzed the proteomics data. C.C. and J.C. provided suggestions for transcriptomic and proteomic analysis. K.P., F.C and N.Z. performed DNA content analysis. C.A. and M.X. quantified chlorophyll and carotenoid contents. W.M. performed all spectroscopic measurements of photosynthetic parameters. J.J. and L.P. performed O_2_ evolution measurements. N.Z. and J.J. optimized the ROS protocol. N.Z., E.B. and K.J.C. performed TEM analysis. R.Z., E.M., and N.Z. led the writing of the manuscript with the contribution of all other co-authors. R.Z., E.M., N.Z., M.S., B.V., T.M., J.J, K.J.C., J.C.B., K.P, B.E. revised the manuscript. All the authors discussed the results and contributed to data interpretation.

## Competing interests

The authors declare no competing interests.

## Supplemental Figures

**Supplemental Figure 1:**
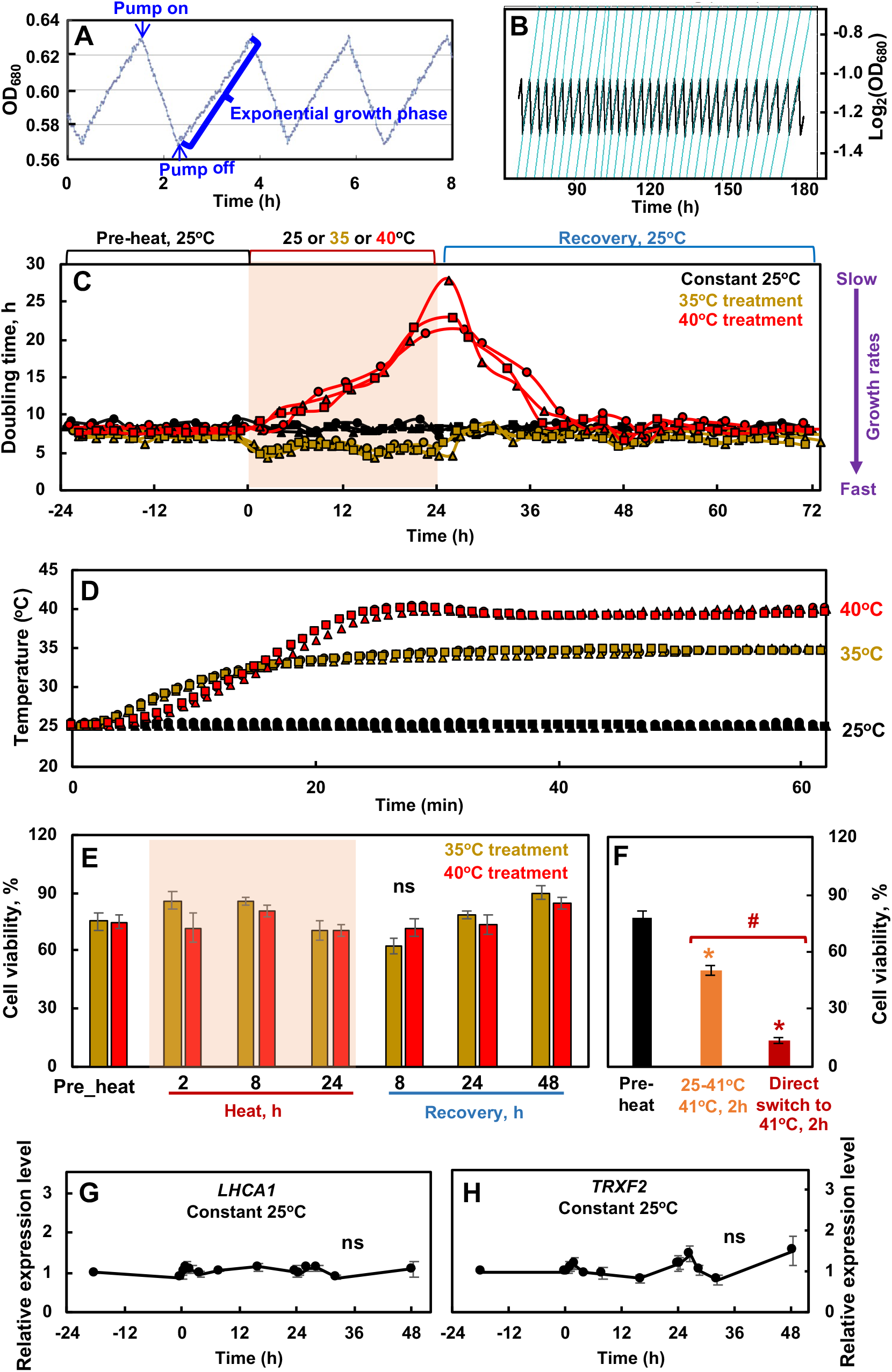
Algal cultures were grown in photobioreactors (PBR) under well-controlled conditions with turbidostatic mode for different temperature treatments. (**A**) PBR cultures were grown with air level CO_2_ in Tris-acetate-phosphate (TAP) medium under constant light (100 µmol photons m^−2^ s^−1^ with equal amount of blue and red light) turbidostatically within a small range of OD_680_ which monitors chlorophyll content. When PBR cultures grew to the set maximum OD_680_, pumps turned on to add fresh medium and dilute the cultures to the set minimal OD_680_, then pumps turned off to allow for exponential growth to the maximum set OD_680_. (B) The doubling time of the PBR cultures were calculated by fitting the OD_680_ curves during exponential growth. (C) Doubling time of the PBR cultures before, during and after treatment at 35, 40, or 25°C. Doubling time is inverse of relative growth rates and smaller doubling time represents faster growth. Three independent biological replicates are plotted for each treatment. Constant 25°C served as controls which showed steady growth without heat treatment. (D) PBR heating profiles. PBR temperatures changed from 25 to 35 or 40°C gradually within 30 min. Three independent biological replicates are plotted for each temperature treatment. (E) Heat treatment in PBRs at 35 or 40°C up to 24 h did not affect cell viability. Algal cells with different heat treatments were diluted and spotted on TAP plates, grown under 150 µmol photons m^−2^ s^−1^ white LED light, 25°C for 44 h before microscopic imaging. Colony numbers were quantified using ImageJ. Cell viability was calculated as the number of colonies on plates divided by the number of cells spotted. Values are mean ± SE, *n* = 3 biological replicates. Statistical analyses were performed with two-tailed t-test assuming unequal variance by comparing different time points with the pre-heat samples. No significance (ns) among different time points (p>0.05). (F) Heating speed affected algal cell viability and direct heating at 41°C in water bath significantly reduced algal cell viability. Algal cultures were harvested from PBRs before heat treatment (pre-heat, black bars), incubated in a water bath which was heated from 25 to 41°C gradually in 25 min then kept at 41°C for 2 h (orange bars), or directly heated in a water bath which was pre-heated to 41°C (sharp temperature switch) then kept at 41°C for 2 h (red bars). Cell viability was quantified as in E. Values are mean ± SE, *n* = 3 biological replicates. Statistical analyses were performed using two-tailed t-tests assuming unequal variance by comparing treated samples with pre-heat (*, p<0.05, the colors of asterisks match the treatment conditions) or between the two heating methods (#, p<0.05). (G, H) The circadian regulated genes *LHCA1* and *TRXF2* had constant expression levels without heat treatments. The relative expressions were calculated from RT-qPCR by normalizing to the reference gene *CBLP* and pre-heat-stress level. Mean ± SE, *n* = 3 biological replicates. Statistical analyses were performed with two-tailed t-test assuming unequal variance by comparing different time points with the first time point. No significance (ns) among different time points (p>0.05). (C, E) Red shaded area depicts the duration of high temperatures.

**Supplemental Figure 2.**
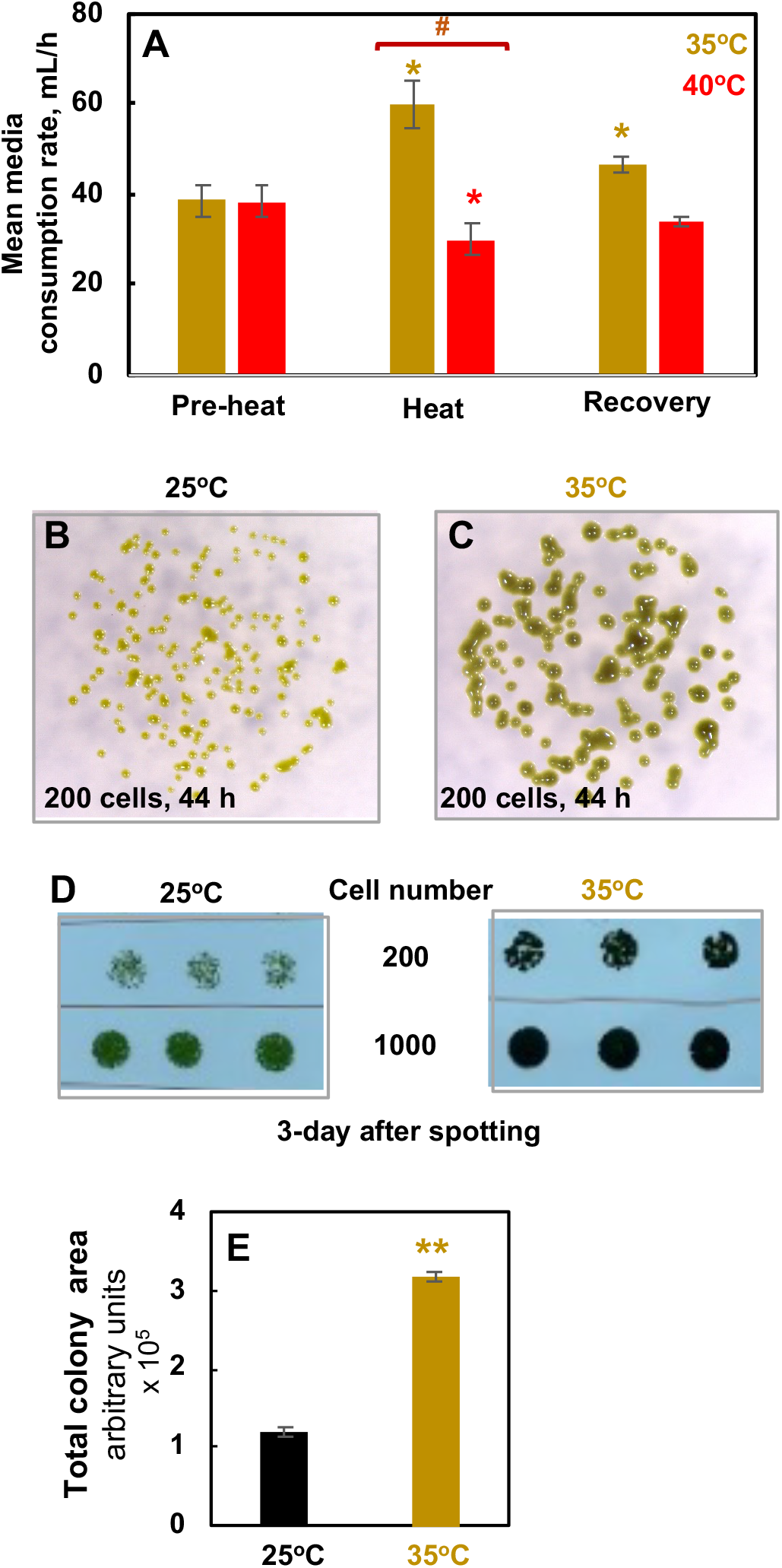
Moderate high temperature at 35°C increased algal growth rates. (**A**) PBR mean media consumption before heat, at the end of 24-h heat of 35 or 40°C, and at the end of the recovery at 25°C, total media consumption volume divided by time. Cell growth induced turbidostatic dilution and consumed medium. Mean ± SE, *n* = 4. (**B-E**) The increased growth rates under 35°C were confirmed by spotting tests on plates. Algal cells harvested from PBRs at 25°C without heat treatments were diluted and spotted on TAP plates, grown under 150 µmol photons m^−2^ s^−1^ white LED light, in incubators of 25 or 35°C for 44 h (**B, C**) or 3 days (**D**) before imaging. B, C, D show one representative of three biological replicates. (**E**) Algal spots with 200 cells were imaged after 44-h growth and colony areas were quantified using ImageJ. Values are mean ± SE, *n* = 3 biological replicates. (**A**, **E**) Statistical analyses were performed with two-tailed t-test assuming unequal variance by comparing treated conditions with the pre-heat or the 25°C conditions (*, p<0.05; **, p<0.01; the colors of asterisks match treatment conditions) or between 35 and 40°C heat treatment (#, p<0.05).

**Supplemental Figure 3.**
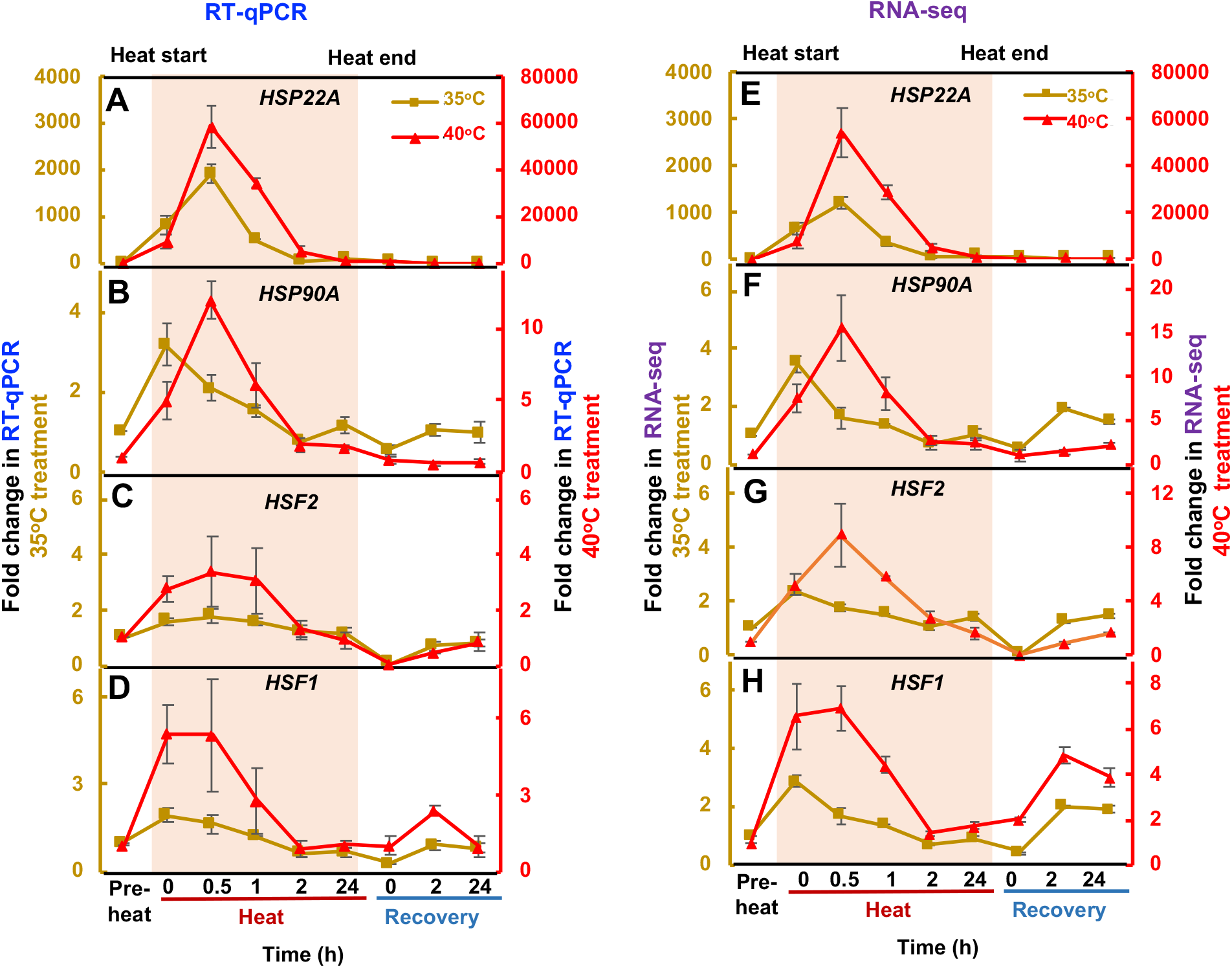
RT-qPCR analysis was consistent with RNA-seq results. Transcript fold-changes of two heat stress marker genes (*HSP22A*, *HSP90A*) and two heat shock transcription factors (*HSF1* and *HSF2*) were calculated based on RT-qPCR (**A**-**D**) or RNA-seq (**E-H**) results. Different y scales were used for samples with 35°C (left) or 40°C (right) treatments. Red shaded area depicts the duration of high temperature. For RT-qPCR results, the fold-changes were calculated by normalizing the relative expression values at different time points with different treatments to the reference gene *CBLP* and the pre-heat time point. For RNA-seq results, the fold-changes were calculated based on Transcripts Per Million (TPM) normalized RNA-seq read counts. Values are mean ± SE, *n* = 3 biological replicates.

**Supplemental Figure 4:**
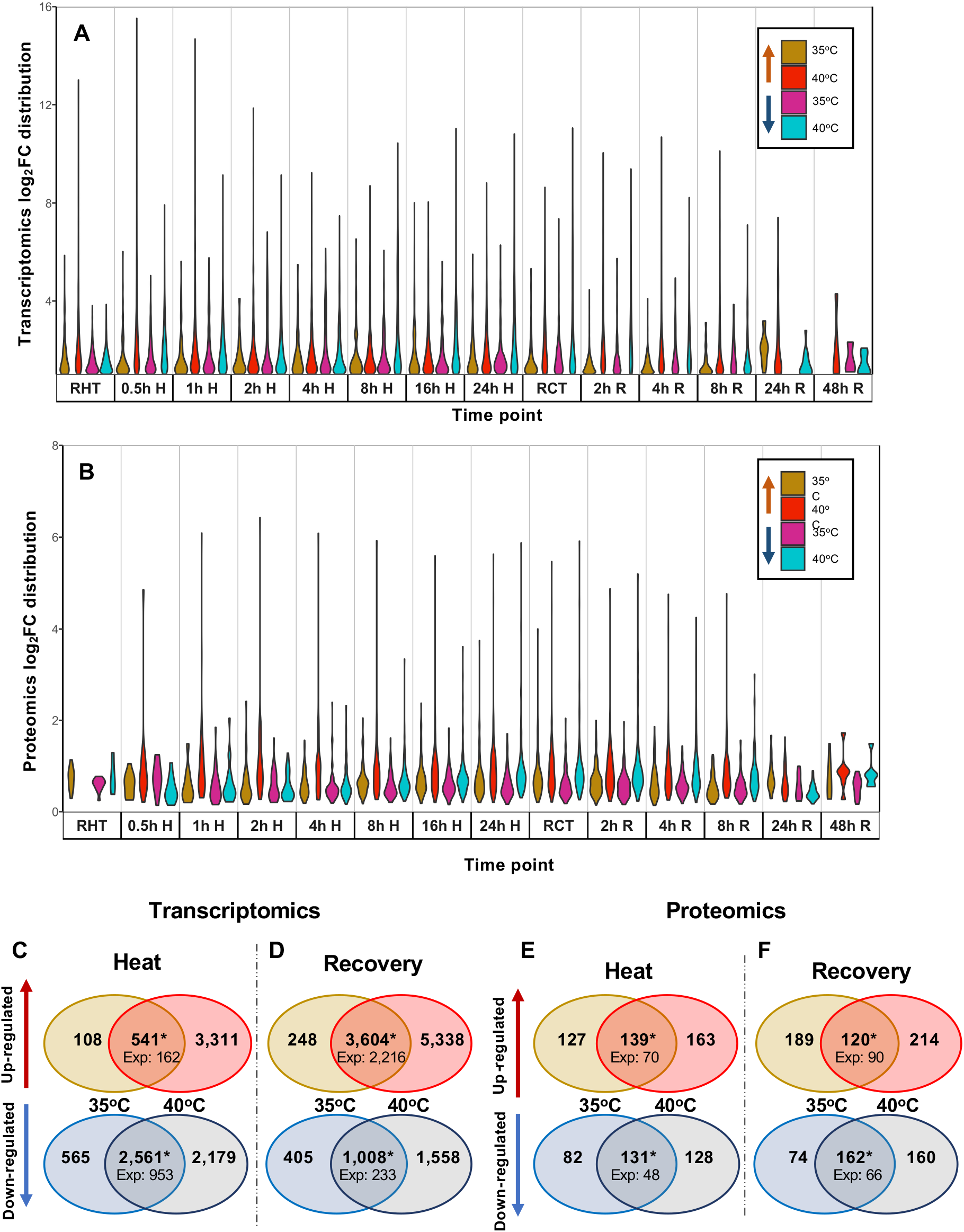
Transcriptomic and proteomic analyses revealed shared and unique regulation of transcripts and proteins during and after heat treatments of 35 or 40°C. (**A, B**) Log_2_(fold-change, FC) distribution of Differentially Expressed Genes (DEGs) and Differentially Accumulated Proteins (DAPs) at different time points, respectively. For each time point, the first two violins represent up-regulated transcripts/proteins, and the last two violins represent down-regulated transcripts/proteins. For down-regulated transcripts/proteins, the inverse of the log_2_FC was displayed. The width of the violins is proportional to the fraction of transcripts/proteins at a certain fold-change value out of the total DEGs/DAPs at a time point. (**C-F**) Venn diagrams of transcripts (**C**, **D**) and proteins (**E, F**) differentially expressed in at least one time point during heat treatment (**C, E**) or recovery (**D, F**). For each panel: top, up-regulated transcripts/proteins; bottom: down-regulated transcripts/proteins. Only transcripts and proteins identified in both treatment groups were used for this analysis. Expected values are the number of transcripts/proteins expected to have overlapping differential expression between the 35 and 40°C treatment groups based on random chance (Fisher’s exact test, *: p< 1.29 x10^−226^).

**Supplemental Figure 5:**
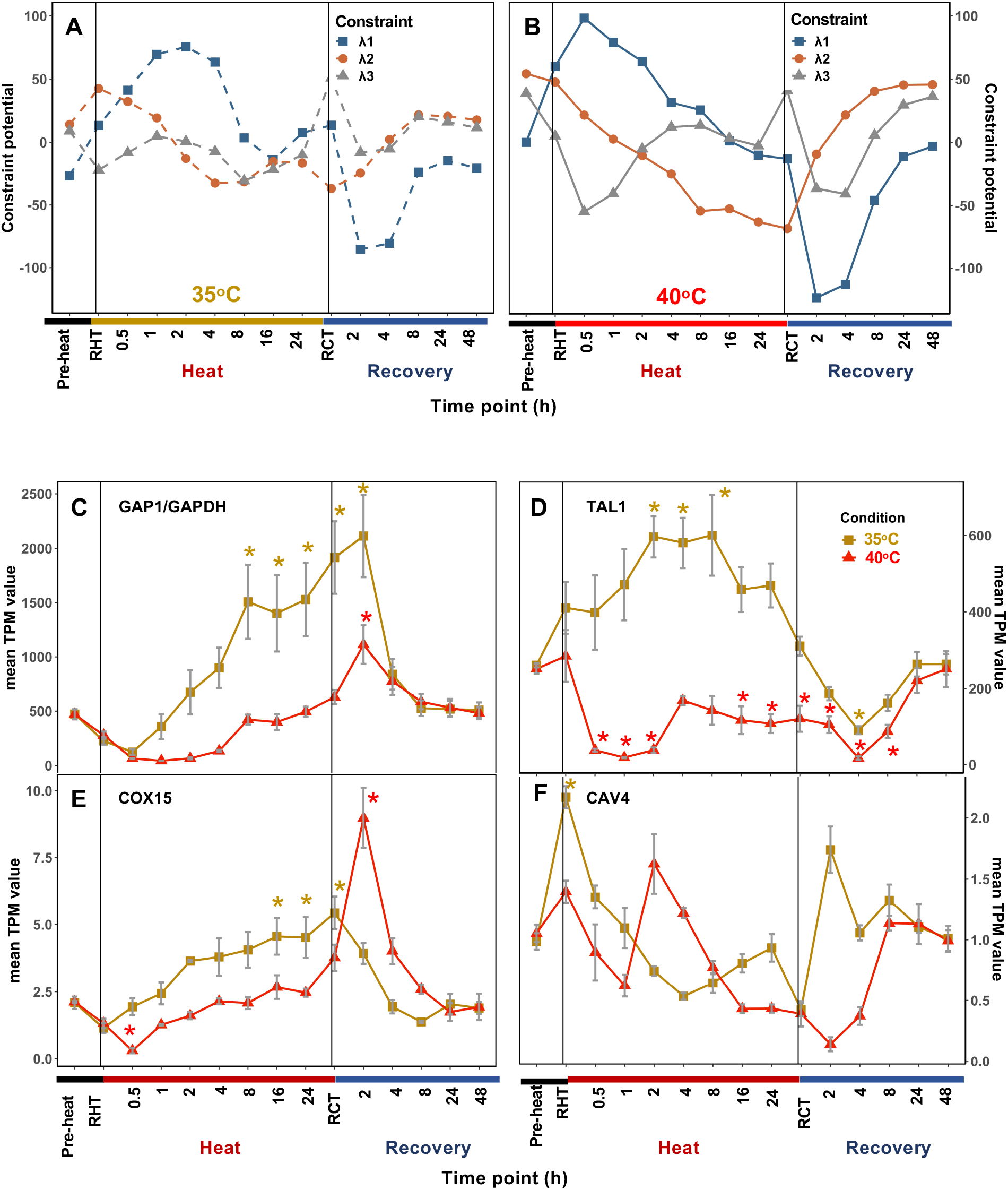
Global transcription patterns were similar between 35 and 40°C treatments, but detailed analysis revealed uniquely differentially expressed genes in the 35°C treatment. Time course of the three major constraint potentials (l1-3) derived from surprisal analysis for 35°C (**A**) and 40°C (**B**) experiments, respectively. The constraint potentials indicate the most important transcriptional patterns during the time course. **(C-F)** Mean transcript per million (TPM) read counts at each time point for select genes that were uniquely up-regulated during 35°C (brown) but not 40°C (red) heat treatment period. Values are mean ± SE, *n* = 3 biological replicates, asterisks indicate significance in differential expression modelling. (**C**) GAP1/GAPDH: Cre12.g485150, Glyceraldehyde 3-phosphate dehydrogenase, involved in gluconeogenesis, glycolysis, and Calvin-Benson Cycle; (**D**)TAL1: Cre01.g032650, transaldolase, involved in the pentose phosphate pathway, which acts upstream of the glycolytic and gluconeogenic pathways; (**E**) COX15: Cre02.g082700, encoding mitochondrial cytochrome c oxidase assembly factor; (**F**) CAV4: Cre11.g467528, encoding a putative voltage-gated calcium channel. Vertical black lines indicate the start and end of heat treatments. Time points were labeled at the bottom. RHT, reach high temperature of 35 or 40°C. RCT, reach control temperature of 25°C for recovery after heat treatment.

**Supplemental Figure 6:**
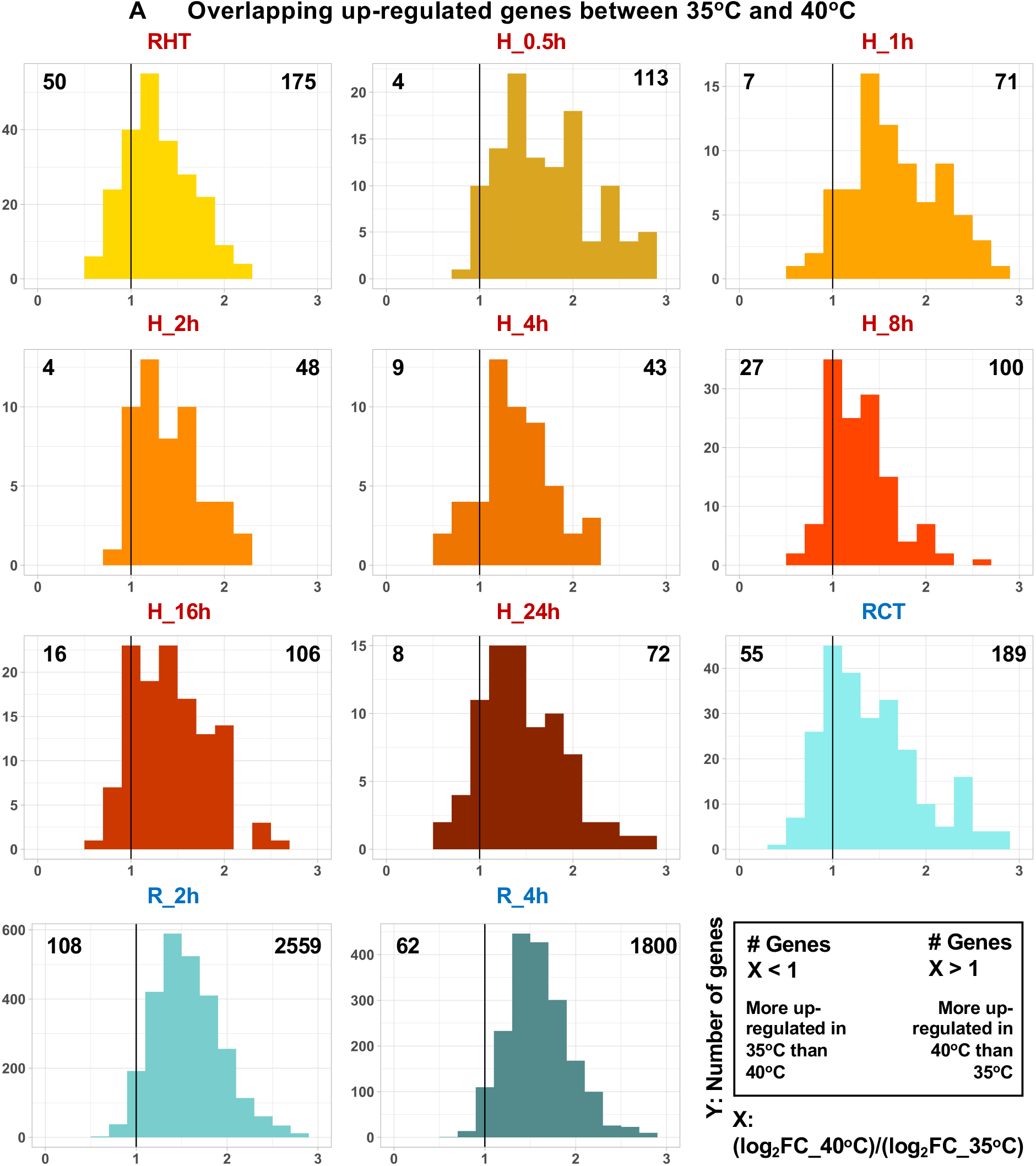
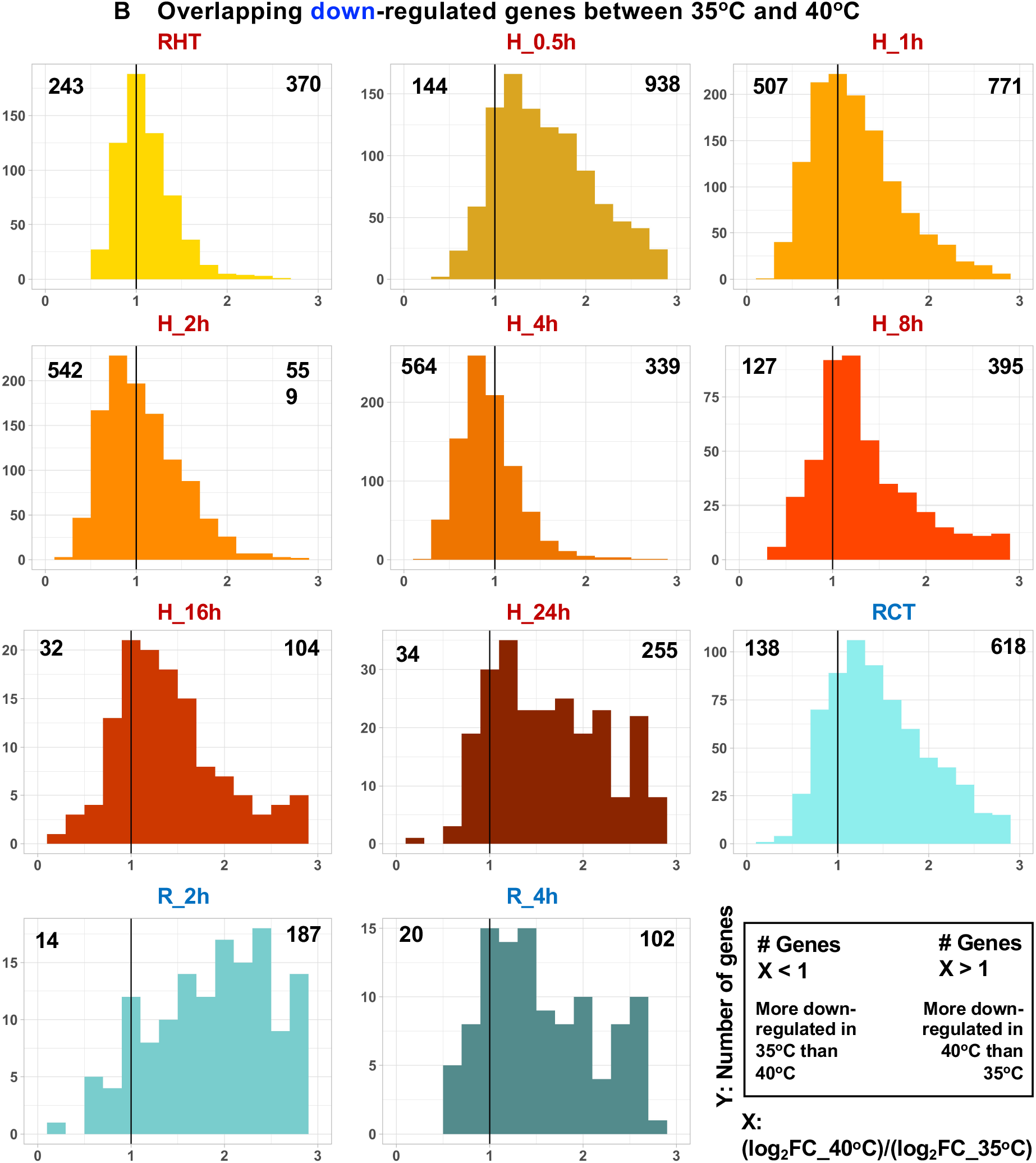
Some overlapping DEGs between 35 and 40°C were more differentially regulated with 35 than 40°C treatment. DEGs, differentially expressed genes. FC, fold-change. Histograms of log_2_(FC in 40°C)/log_2_(FC in 35°C) for overlapping up-regulated (**A**) and down-regulated (**B**) genes between 35 and 40°C are displayed for each time point. Very few overlapping DEGs between 35 and 40°C were identified at 8, 24, and 48 h of recovery, which were thus omitted. Black vertical lines indicate equal differential expression between 35 and 40°C treatments. Bars to the left of the black line indicate genes more differentially expressed in the 35°C treatment group while bars to the right of the black line indicate genes more differentially expressed in the 40°C treatment group. Numbers in the top left and right corners of each histogram represent the number of genes with higher fold change values in 35 or 40°C, respectively. Pre-heat, before heat treatments. RHT, reach high temperature of 35 or 40°C. H_1h, heat at 35 or 40°C for 1 h, similar names for other time points during heat. RCT, reach control temperature of 25°C for recovery after heat. R_2h, recovery at 25°C for 2 h, similar names for other time points during recovery.

**Supplemental Figure 7:**
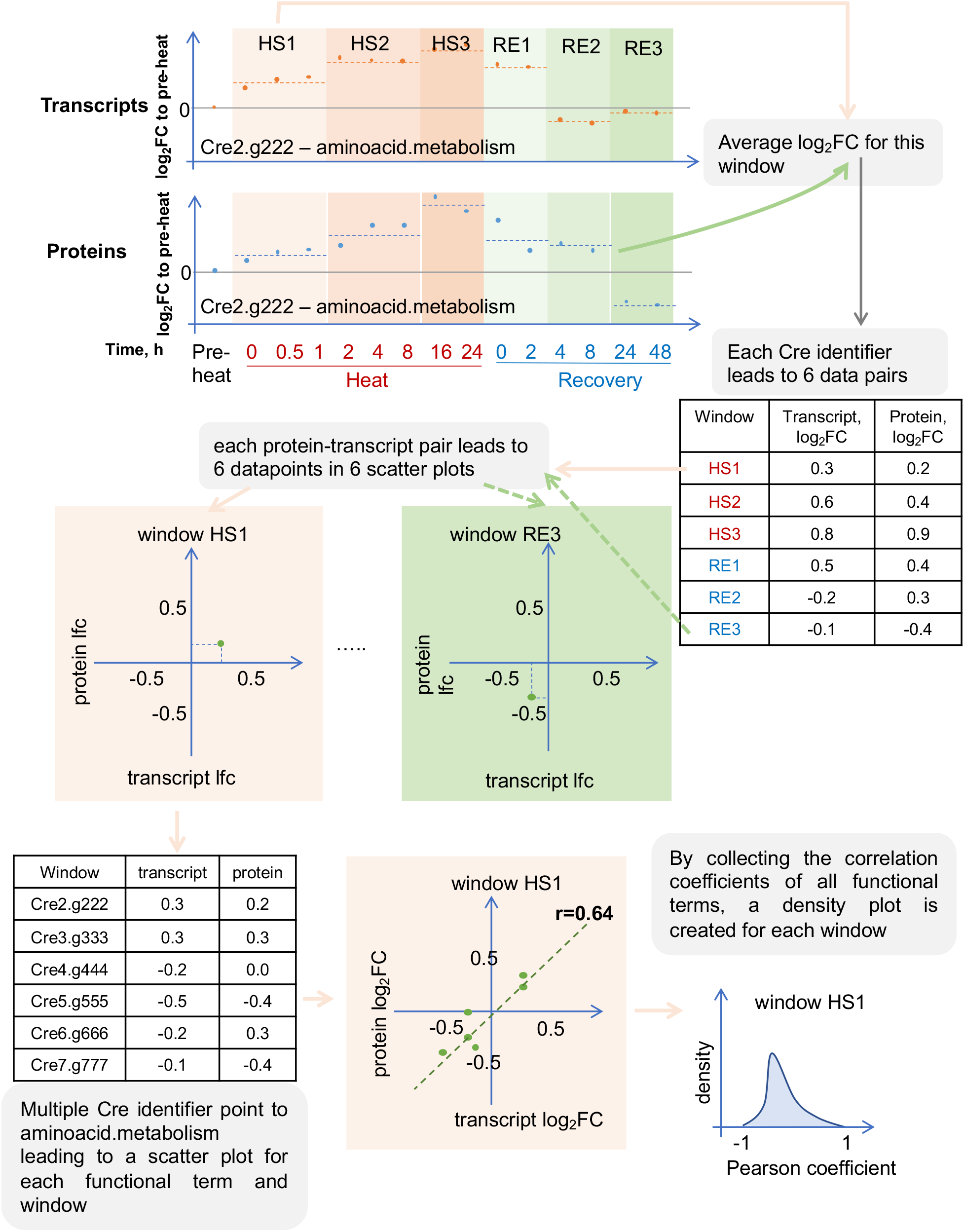
The fold-change correlation between transcripts and proteins were investigated. Transcript and protein correlation analysis by using Cre2.g222 (hypothetical gene ID as Cre identifier) of functional term aminoacid.metabolism as an example. Log_2_(fold-changes, FC) of transcript reads and protein abundance are calculated in respect the pre-heat sample. The heat stress period (HS) as well as the recovery period (RE) are split into three windows each (HS1-3 and RE1-3). HS1-3 windows: 0-1 h, 2-8 h, 16-24 h during the heat period; RE1-3 windows, 0-2 h, 4-8 h, 24-48 h during the recovery period after heat treatment. Every identifier that has transcripts as well as proteins associated to it, results in a transcript-protein fold-change pair for each window. The average Log_2_FC is determined for each transcript-protein pair in each window. By collecting all Cre identifiers that are associated with aminoacid.metabolism, a scatter plot of transcript-protein fold-change pairs is generated and the Pearson correlation coefficient is calculated. By repeating the workflow for all functional terms for each window, a density plot for each of the six windows is created to describe the overall correlation of transcript reads and protein abundance for that window.

**Supplemental Figure 8.**
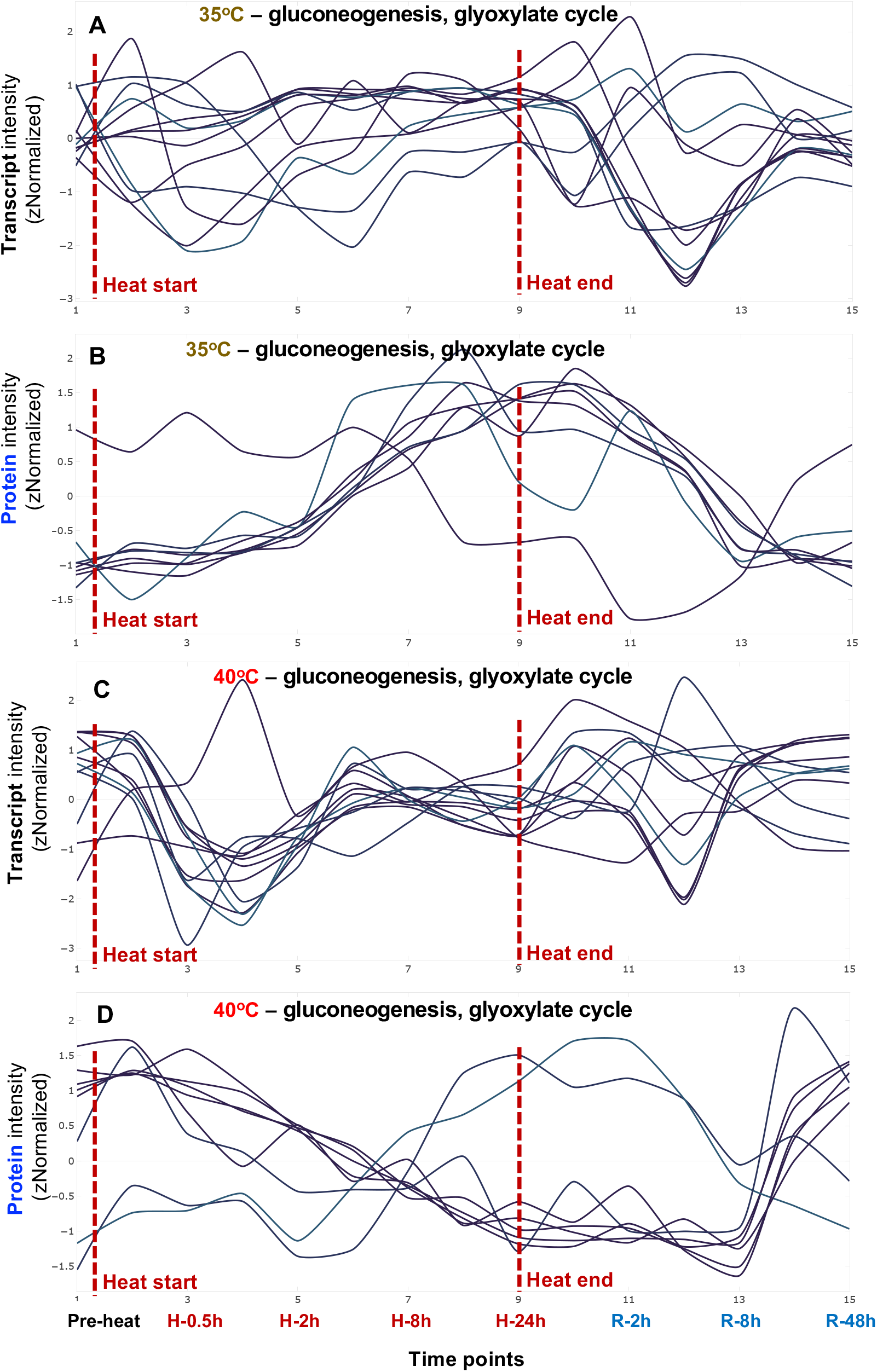
The kinetics of transcripts and proteins suggest gluconeogenesis and glyoxylate cycle increased during 35°C but decreased during 40°C heat. Transcript (**A**, **C**) and protein (**B**, **D**) signals related to the MapMan bin gluconeogenesis and glyoxylate cycle were standardized to z scores (standardized to zero mean and unit variance) and are plotted against equally spaced time point increments. The red dashed lines indicate the start and end time of heat treatment for 35°C (**A**, **B**) and 40°C (**C**, **D**) respectively. Total 12 transcripts and 8 proteins that are quantifiable and related to gluconeogenesis and glyoxylate cycle are plotted. Time points are labeled at the bottom. Timepoint 1: pre-heat. Time points 2-9, heat treatment at 35 or 40°C, including reaching high temperature, 0.5, 1, 2, 4, 8, 16, 24 h during heat; time points 10-15, recovery phase after heat treatment, including reaching control temperature, 2, 4, 8, 24, 48 h during recovery. See the interactive figures with gene IDs and annotations in Supplemental Dataset 10, gluconeogenesis_glyoxylate cycle.html.

**Supplemental Figure 9:**
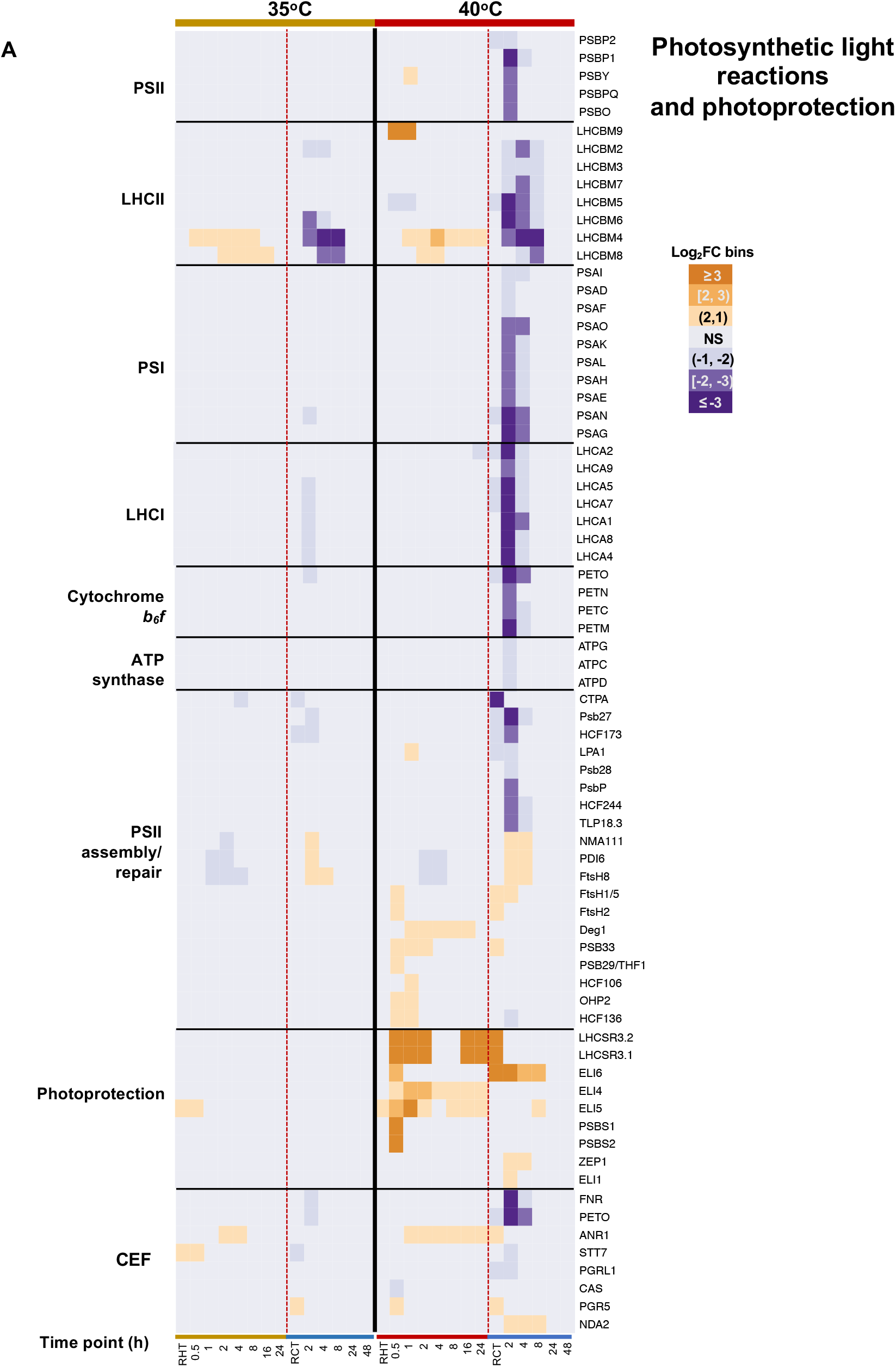
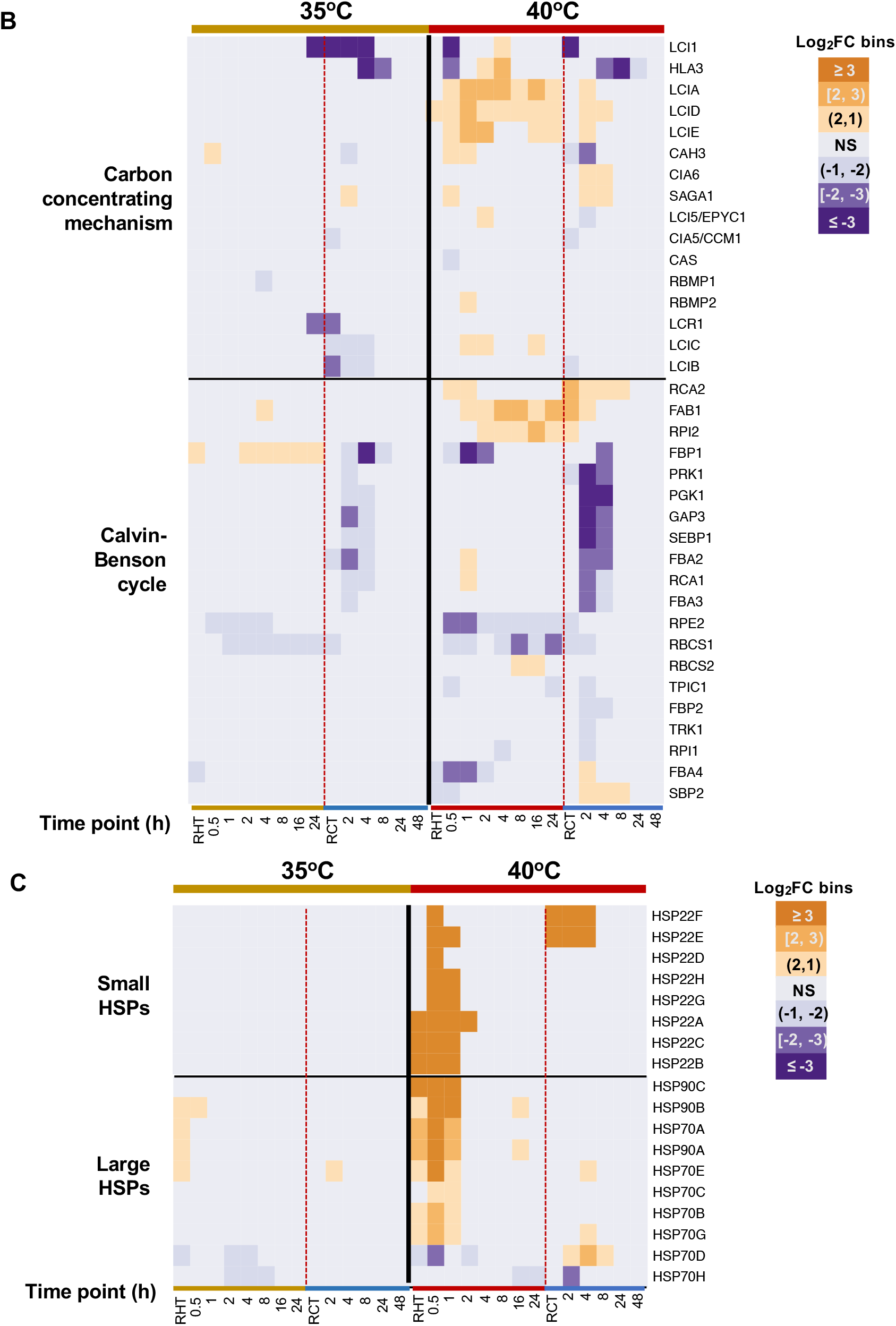
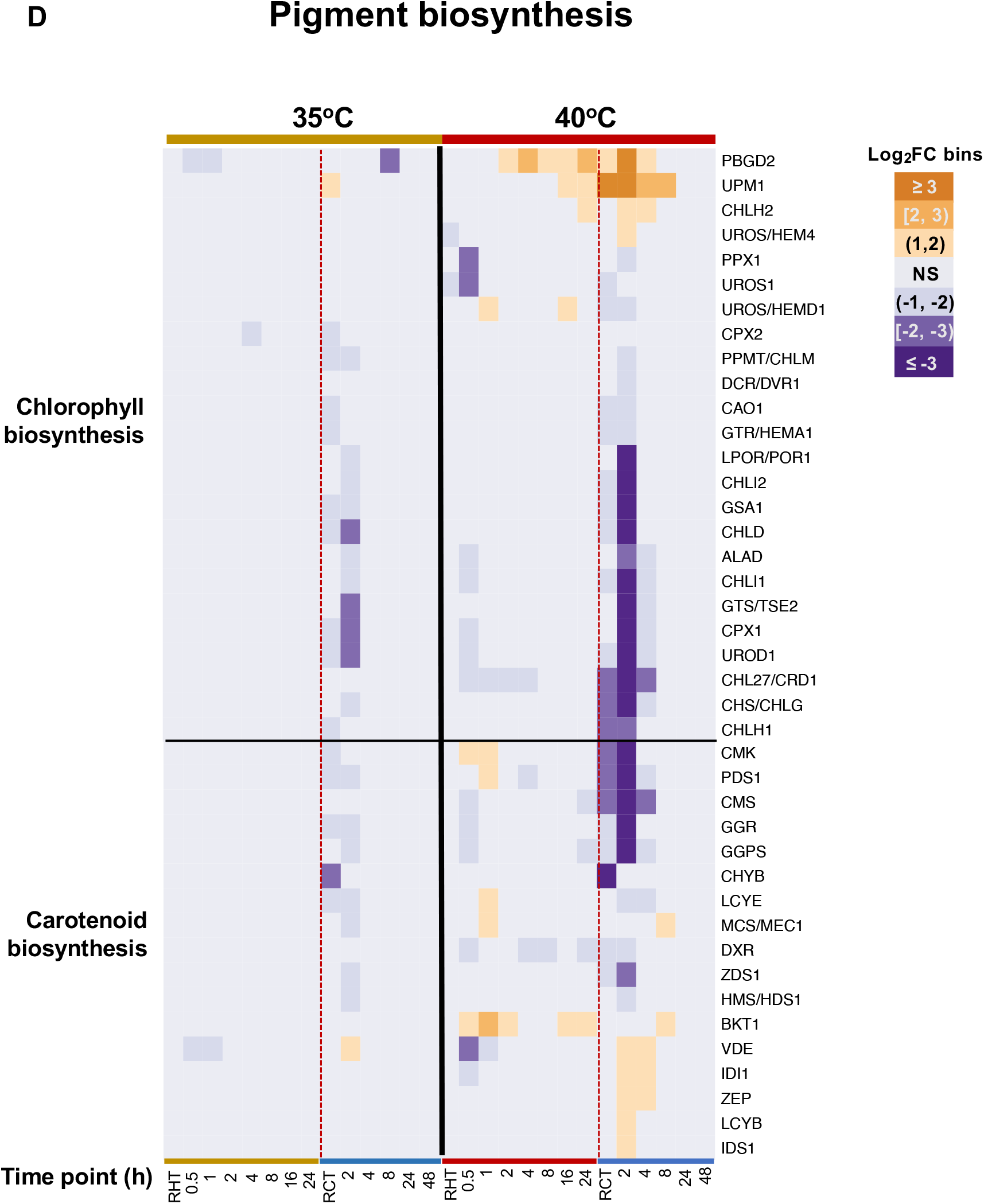
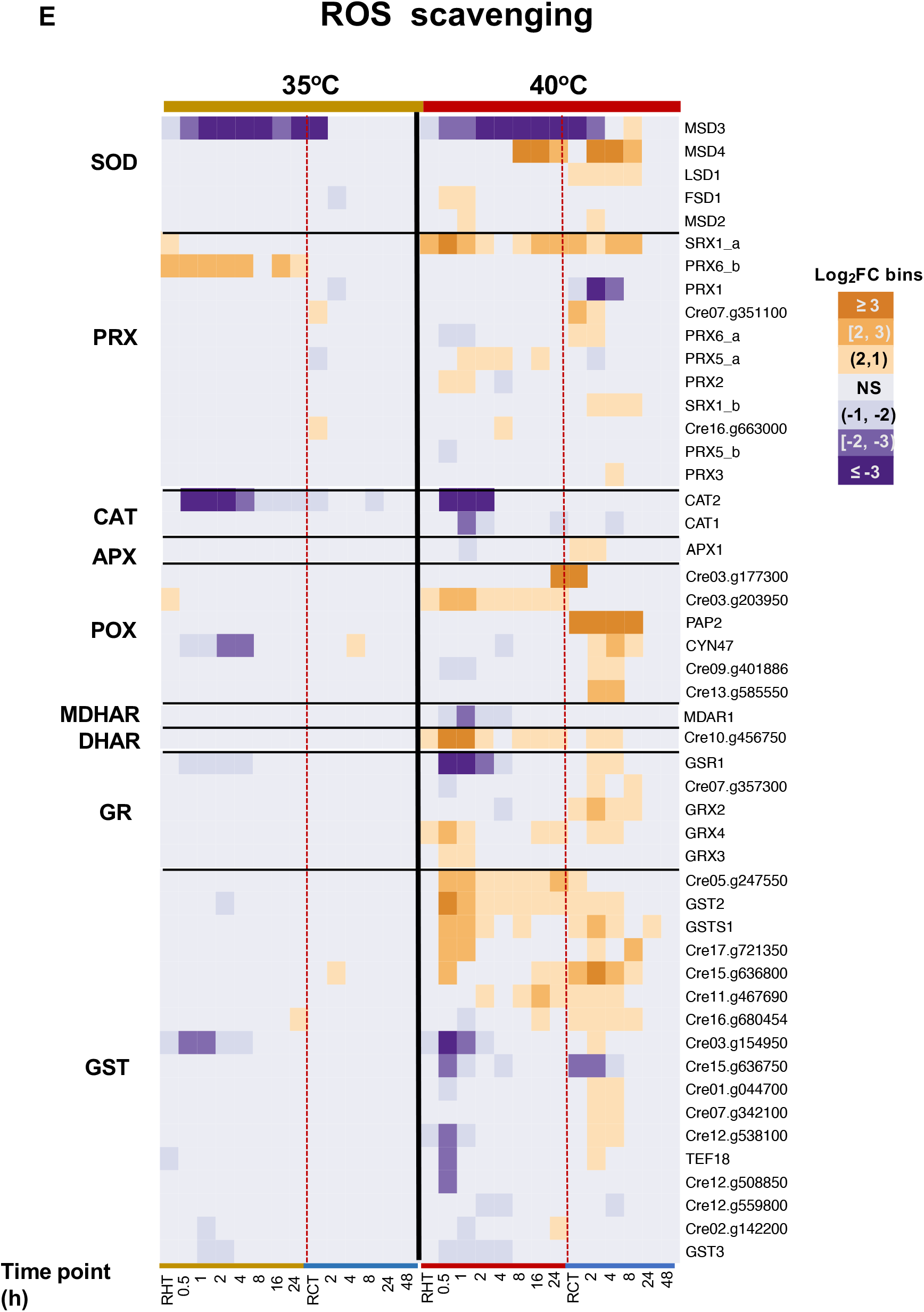
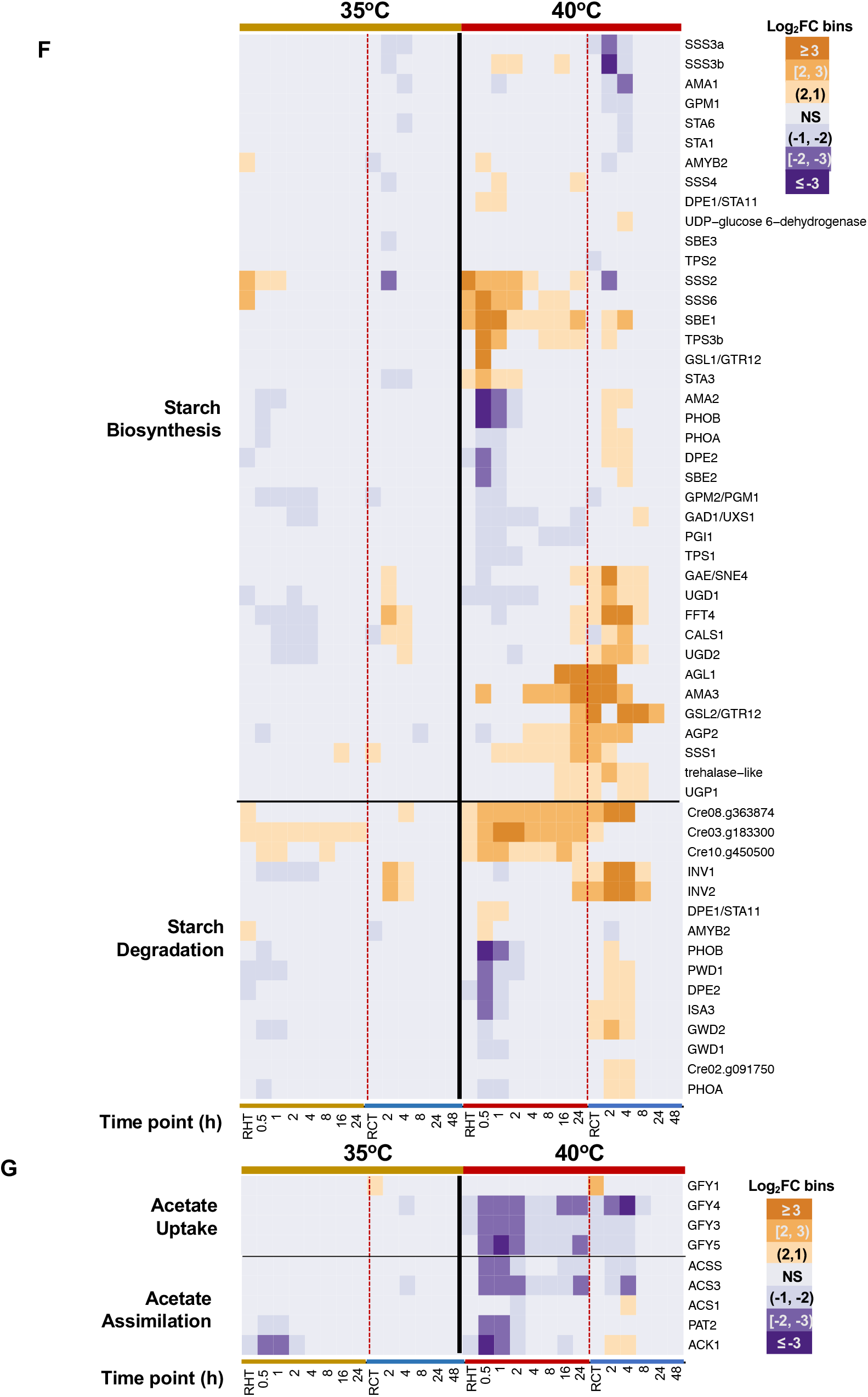
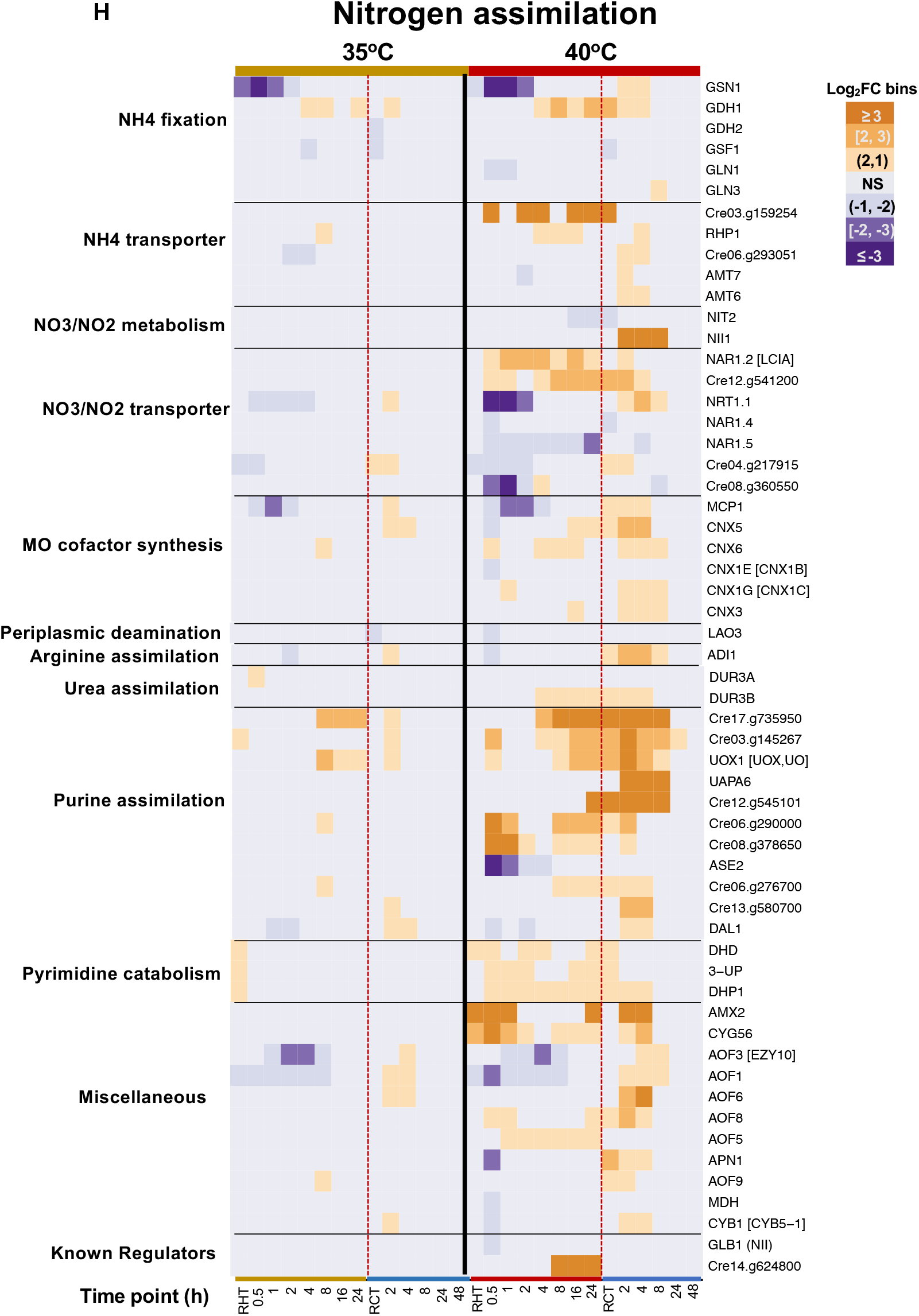
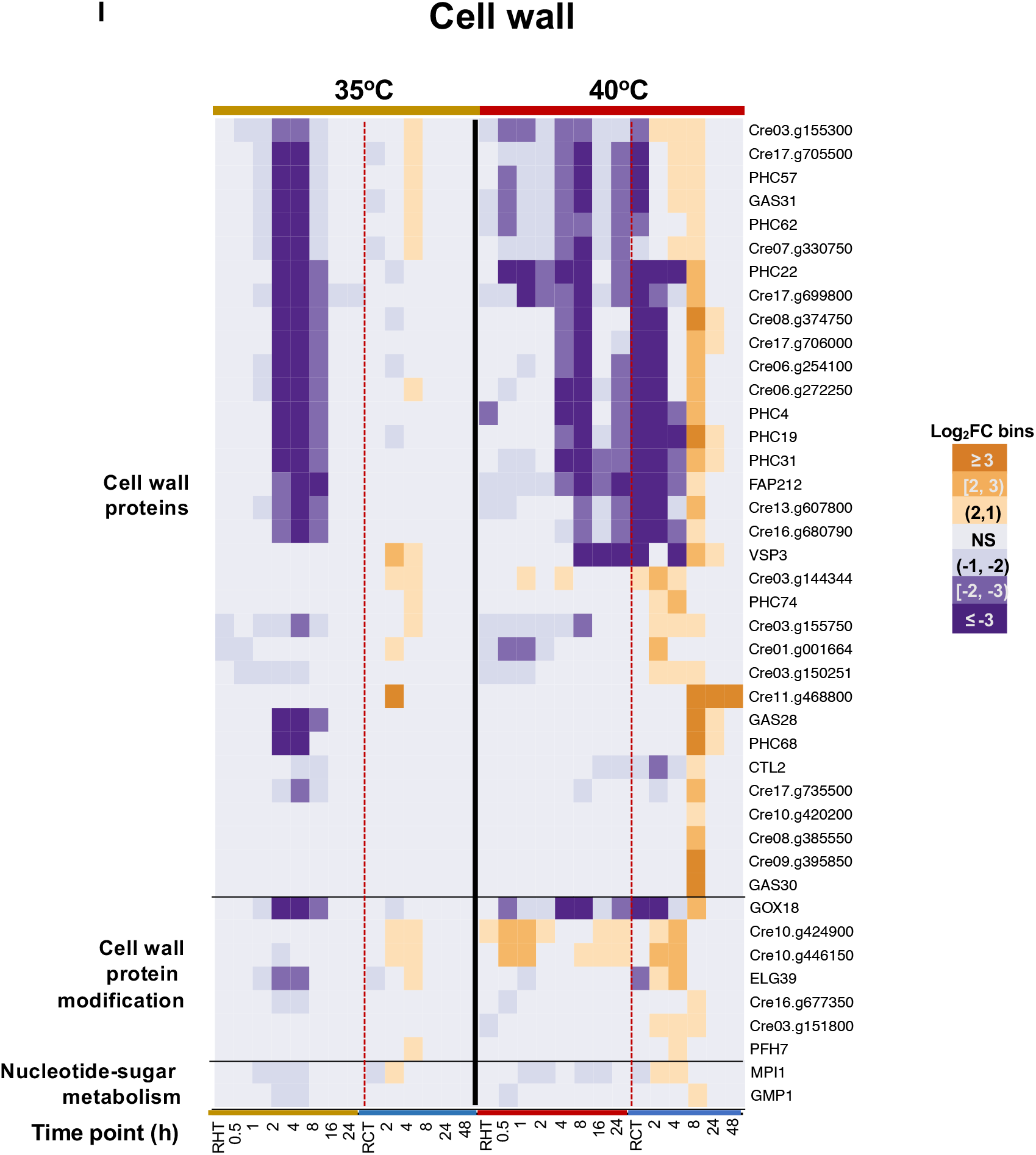
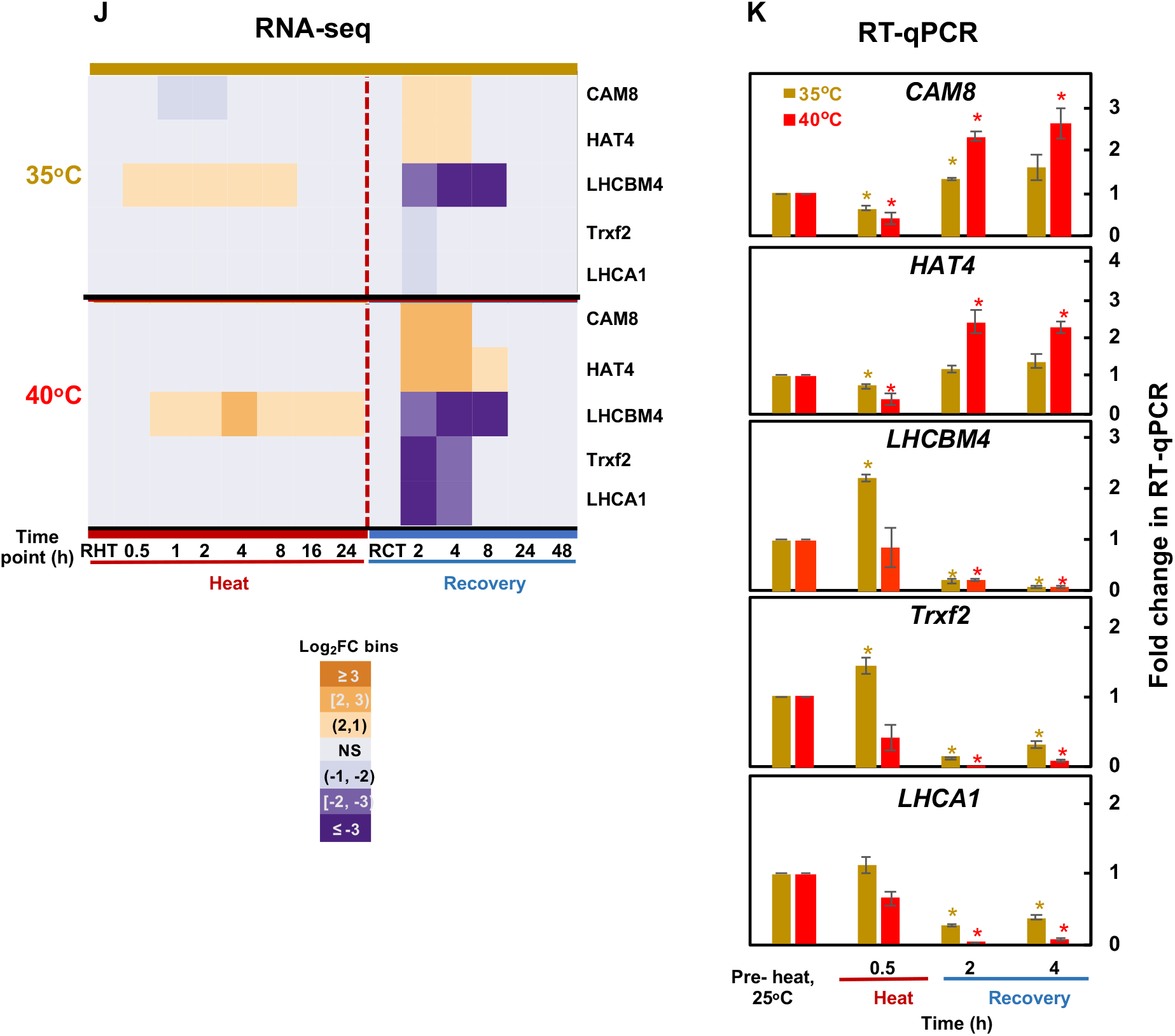
The expression of key pathways was differentially regulated with 35 or 40°C treatment. Expression patterns of differentially expressed genes in the select pathways are displayed: **(A)** Photosynthetic light reactions and photoprotection, **(B)** carbon concentrating and fixation, **(C)** heat shock proteins (HSPs), **(D)** chlorophyll and carotenoid biosynthesis, **(E)** reactive oxygen species (ROS) scavenging pathways, (**F**) starch biosynthesis/degradation, (**G**) acetate uptake and assimilation pathways, (**H**) nitrogen assimilation pathways, (**I**) cell wall pathways. (**J, K**) RT-qPCR results validated several down-regulated genes related to photosynthetic light reactions (bottom three genes) during the recovery of 2 and 4 h. The top two genes showing opposite expression during recovery are served as controls. *CAM8*, Cre03.g150300, encoding calmodulin-like protein; *HAT4*, Cre07.g354100, encoding histone acetyltransferase; *LHCBM4*, Cre06.g283950, encoding light-harvesting Chlorophyll a/b binding protein of LHCII; *TRXF2*, Cre05.g243050, encoding thioredoxin f2; *LHCA1*, Cre06.g283050, encoding light-harvesting protein of photosystem I. **(J)** Fold-change heatmap of select genes. Data for 35°C (top) and 40°C (bottom) treatment groups were separated by a black horizontal solid line. **(G)** RT-qPCR validation of select genes at the indicated time points. For RT-qPCR results, the fold-changes were calculated by normalizing the relative expression values at different time points with different treatments to the reference gene *CBLP* and the pre-heat time point. Values are mean ± SE, *n* = 3 biological replicates. Statistical analyses were performed with two-tailed t-test assuming unequal variance by comparing treated samples with pre-heat samples (*, P<0.05, the color of the asterisks matches the treatment condition). (**A**-**J**) FC, fold-change. Differential expression model output log_2_FC values were sorted into different expression bins as follows: highly up-regulated: log_2_FC ≥ 3; moderately up-regulated: 3 > log_2_FC ≥ 2; slightly up-regulated: 2 > log_2_FC > 1; NS: not significant; slightly down-regulated: −2 < log_2_FC < −1; moderately down-regulated: −2 ≤ log_2_FC < −3; highly down-regulated: log_2_FC ≤ −3. Vertical red dashed lines indicate the switch from high temperature to recovery period in each treatment group. Time points were labeled at the bottom. RHT, reach high temperature of 35 or 40°C. RCT, reach control temperature of 25°C for recovery after heat treatment.

**Supplemental Figure 10:**
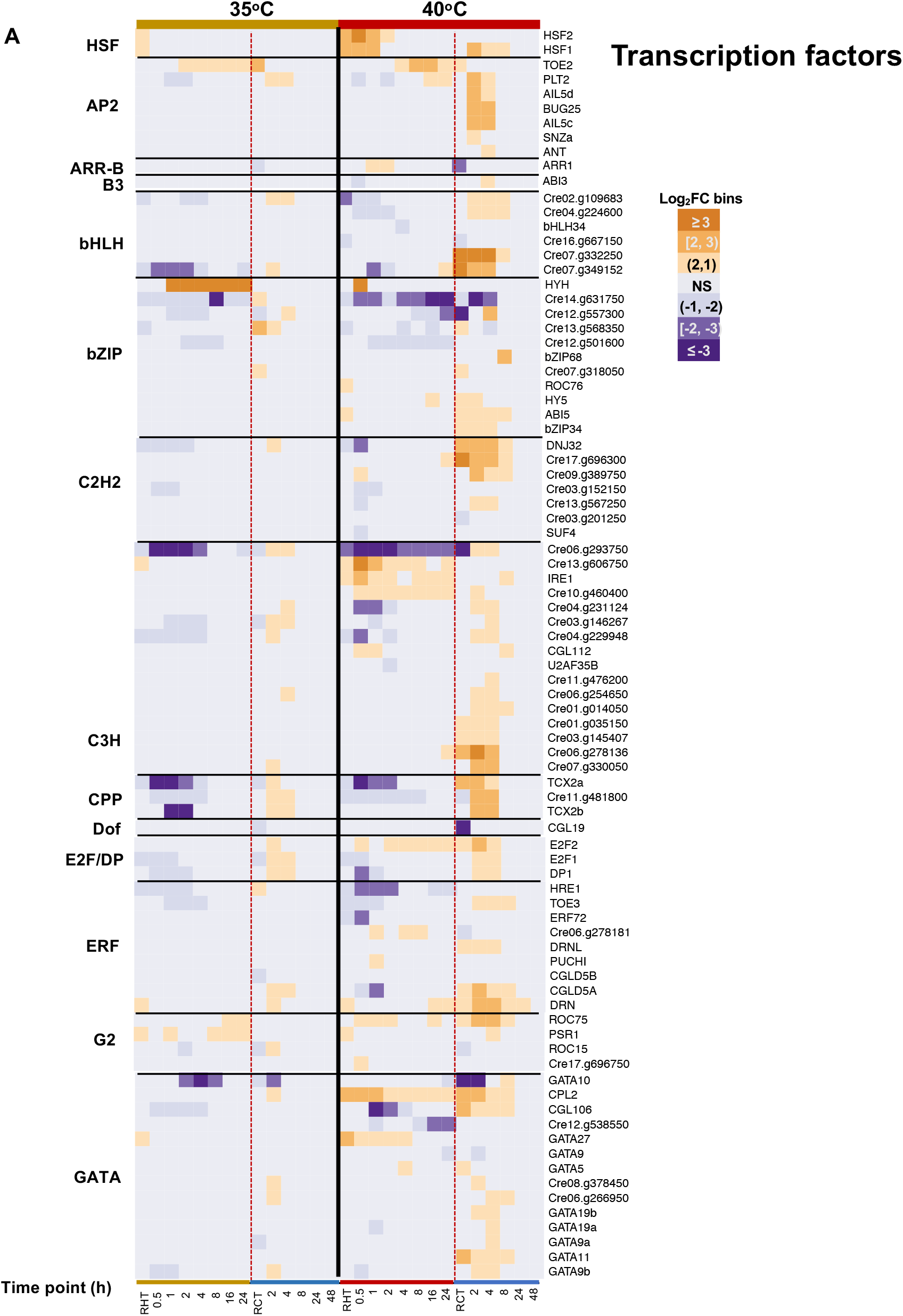
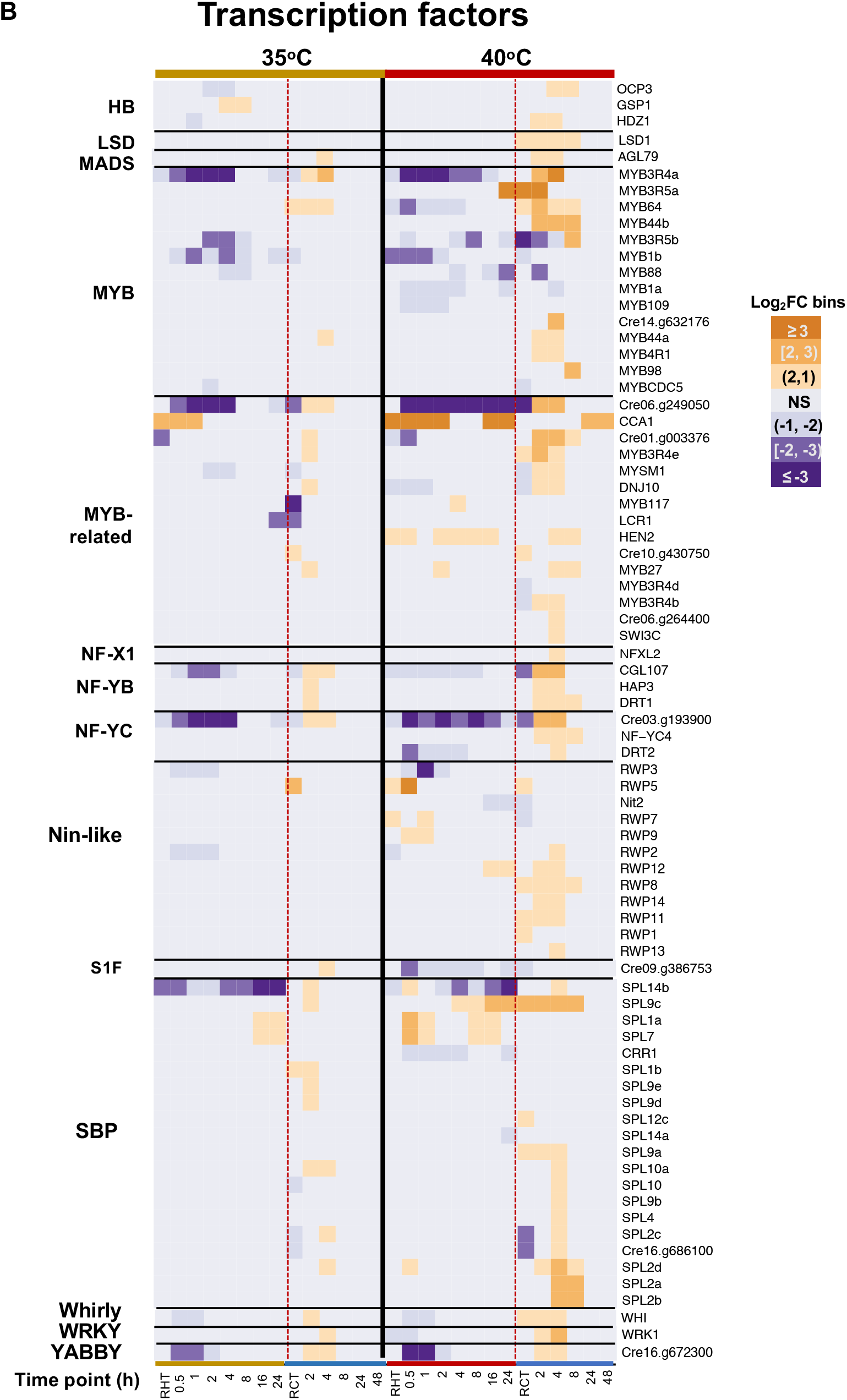
Genes encoding transcription factors had different expression patterns with 35 or 40°C treatment. Expression patterns of differentially expressed genes that encode transcription factors identified using Plant Transcription Factor Database. FC, fold-change. Differential expression model output log_2_FC values were sorted into different expression bins as follows: highly up-regulated: log_2_FC ≥ 3; moderately up-regulated: 3 > log_2_FC ≥ 2; slightly up-regulated: 2 > log_2_FC > 1; NS: not significant; slightly down-regulated: −2 < log_2_FC < −1; moderately down-regulated: −2 ≤ log_2_FC < −3; highly down-regulated: log_2_FC ≤ −3. All treatment time points are shown in 35°C (left) and 40°C (right), separated by a vertical solid black line. Vertical red dashed lines indicate the switch from high temperature to recovery period in each treatment group. Time points were labeled at the bottom. RHT, reach high temperature of 35 or 40°C. RCT, reach control temperature of 25°C for recovery after heat treatment.

**Supplemental Figure 11.**
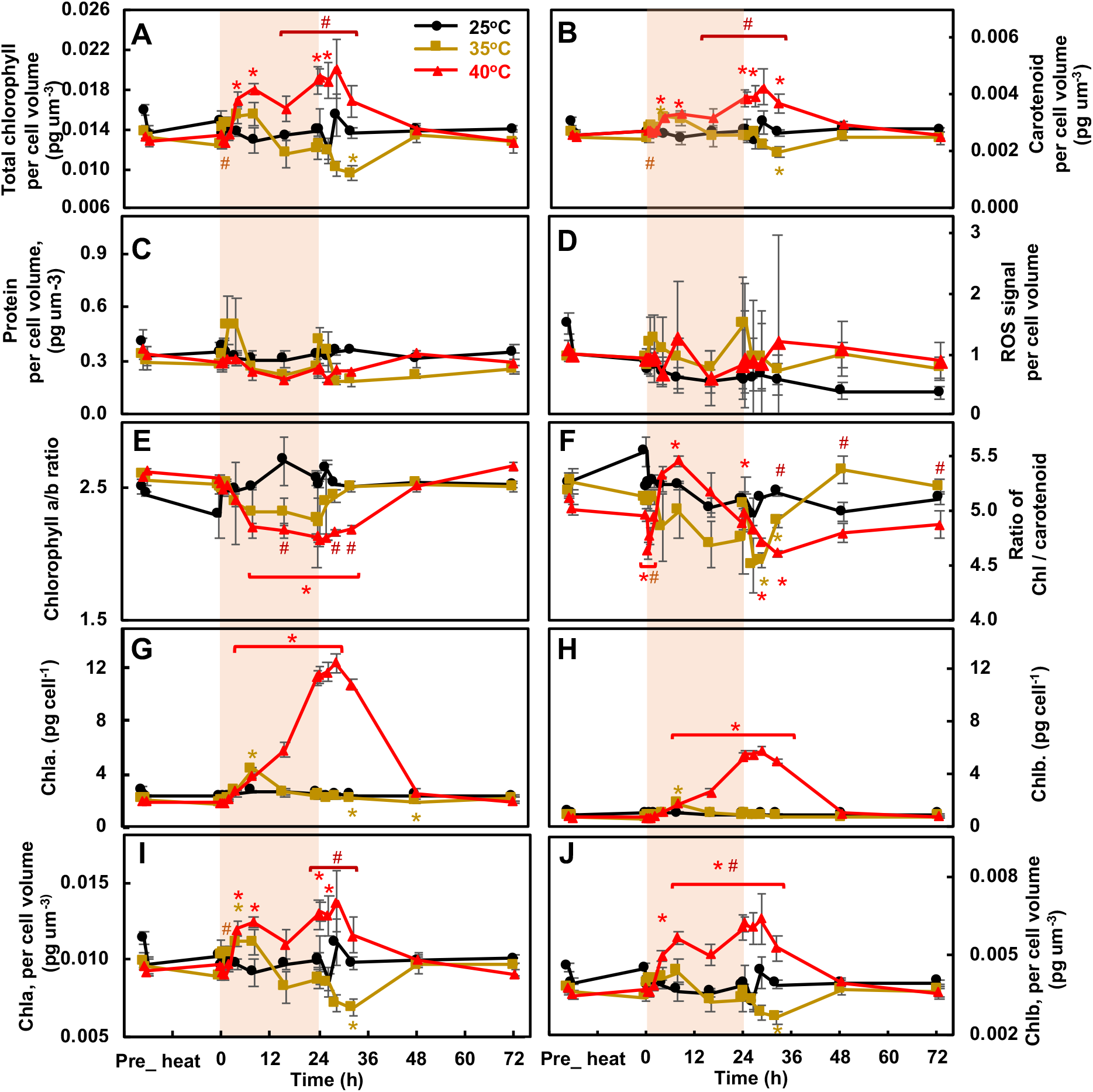
Heat at 40°C increased chlorophyll and carotenoid accumulation per cell volume but not the level of protein or ROS production. (**A-D**) Chlorophyll (Chl), carotenoid, protein, and ROS levels normalized to cell volume, respectively; **(E)** Ratios of Chl a and b**; (F)** Ratios of chlorophyll and carotenoid**; (G, H)** Chla and Chlb per cell; (**I**, **J**) Chla and Chlb per cell volume. Red shaded areas depict the duration of high temperature. Values are mean ± SE, *n* = 4 biological replicates. Statistical analyses were performed with two-tailed t-test assuming unequal variance by comparing treated samples with pre-heat samples (*, p<0.05, the colors of asterisks match treatment conditions) or comparing treatments at 35 and 40°C at the same time point (#, p<0.05).

**Supplemental Figure 12.**
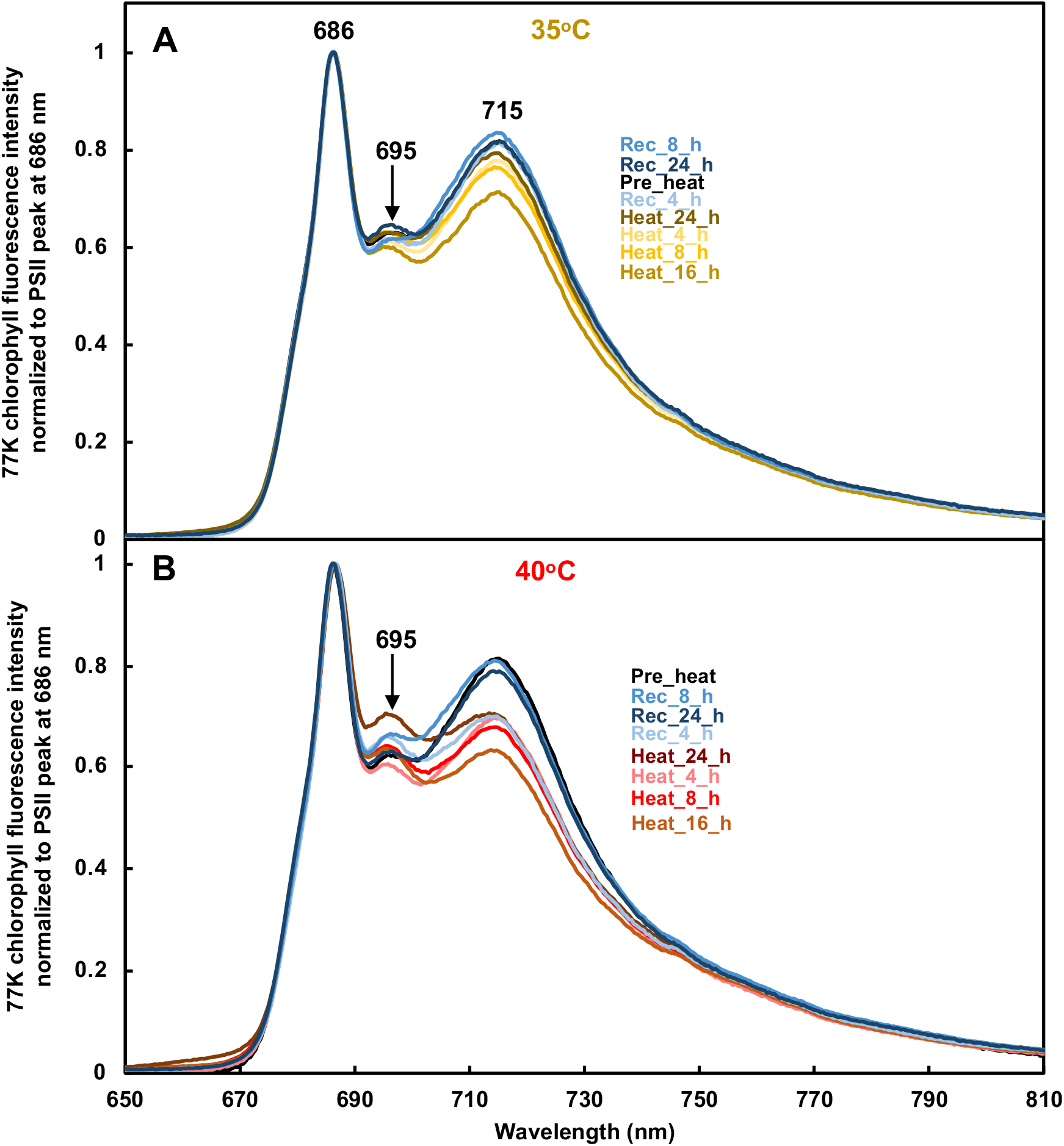
Both 35 and 40°C heat decreased PSI emission peak around 715 nm but increased PSII emission peak around 695 nm in 77 K chlorophyll fluorescence measurement. 77 K chlorophyll fluorescence spectra from algal cells at different time points before, during and after heat treatments at 35 or 40°C. Each curve is an average of 3 biological replicates and normalized to PSII peak at 686 nm.

**Supplemental Figure 13.**
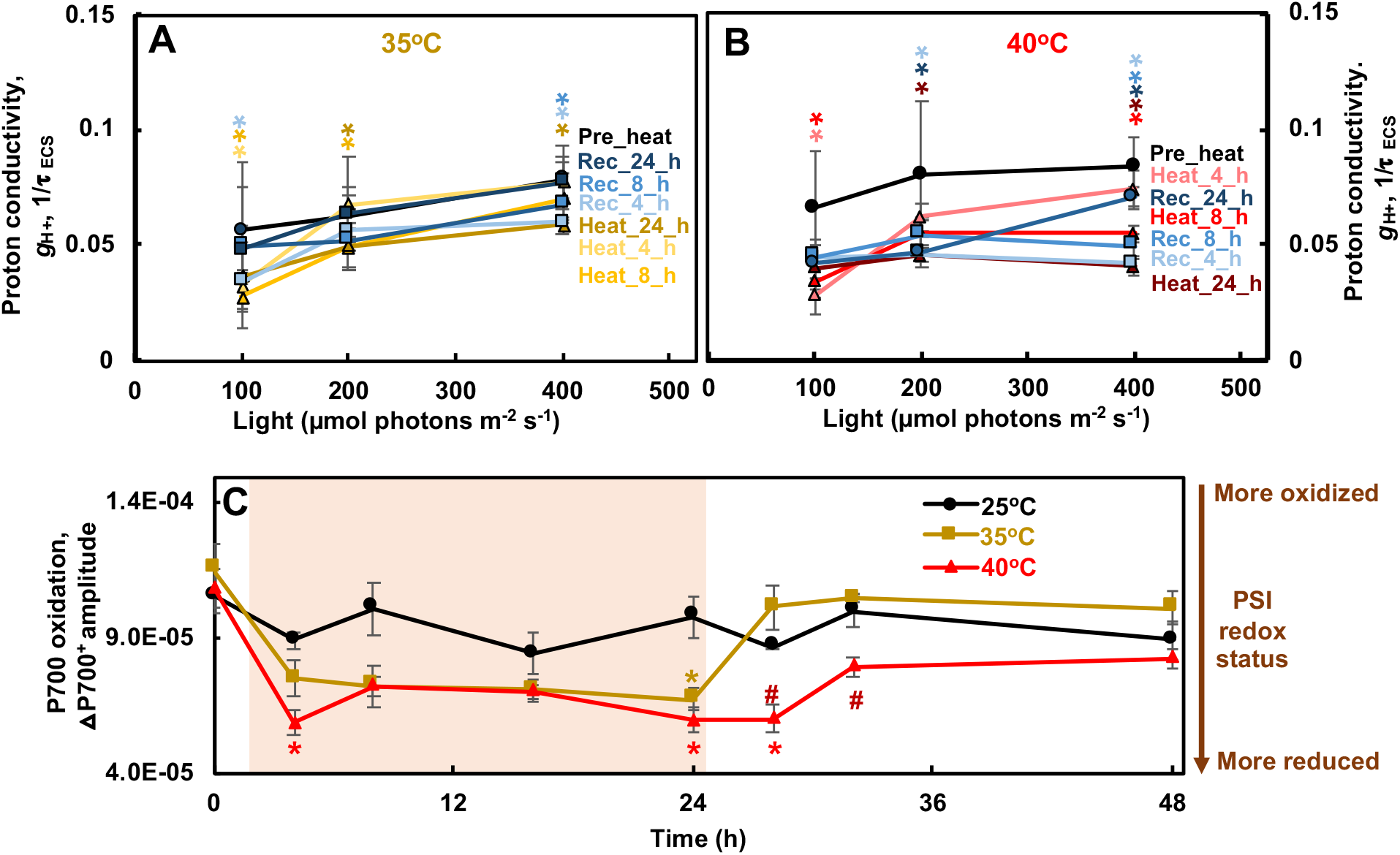
Heat at 35 and 40°C decreased ATP synthase activity and caused reduced PSI. **(A, B)** Proton conductivity (ɡ_H_^+^), proton permeability of the thylakoid membrane and largely dependent on the activity of ATP synthase, inversely proportional to the decay time constant of light-dark transition induced electrochromic shift (ECS) signal (τ_ECS_); ɡ_H_^+^ =1/τ_ECS_. (**C**) P700 oxidation, measured by the amplitude of absorbance change due to far-red induced oxidized P700^+^ followed by dark reduction, normalized to chlorophyll contents. Smaller number means smaller amount of oxidizable P700 and more reduced PSI. Red shaded areas depict the duration of high temperature. Mean ± SE, *n*=3 biological replicates. Statistical analyses were performed using two-tailed t-test assuming unequal variance by comparing treated samples with the pre-heat samples under the same light (**A, B**) or constant 25°C samples at the same time point (**C**). A, B, p values were corrected by FDR. *, p<0.05, the colors and positions of asterisks match the treatment conditions and time points, respectively.

**Supplemental Figure 14.**
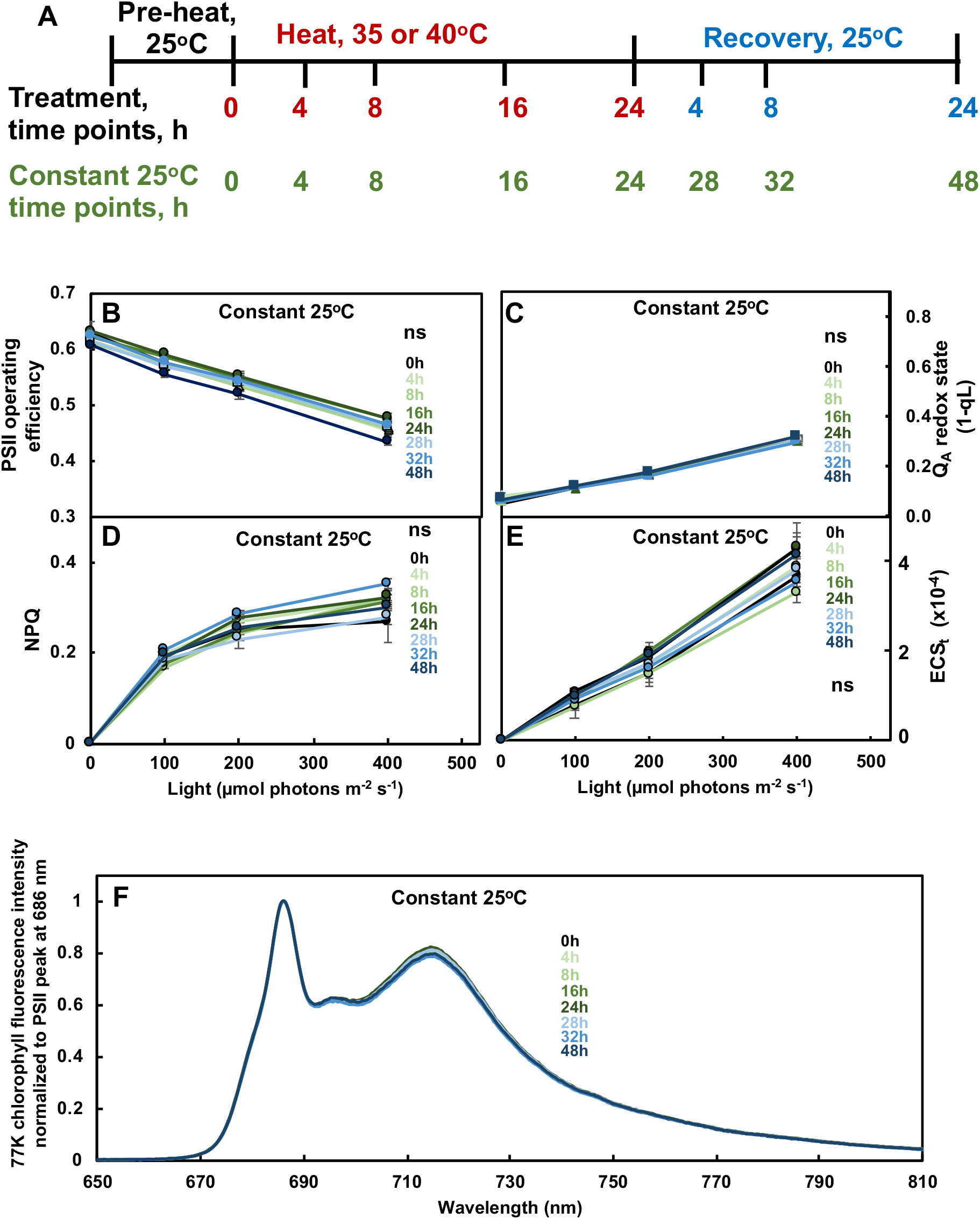
Photosynthetic parameters of algal cultures grown in PBRs with turbidostatic mode were constant without heat treatment. **(A)** Time points before, during and after heat treatments at 35 or 40°C and corresponding time points at constant 25°C. (**B-F**) Photosynthetic parameters from algal cells at different time points during constant 25°C without heat. (**B**) PSII operating efficiency; (**C**) The Q_A_ redox state refers to the redox state of chloroplastic quinone A (Q_A_), the primary electron acceptor downstream of PSII; (**D**) NPQ (non-photochemical quenching); (**E**) ECS_t_, measured by electrochromic shift (ECS), representing the transthylakoid proton motive force, *pmf*. (**B-E**) Values are mean ± SE, *n* = 3 biological replicates. Statistical analyses were performed with two-tailed t-tests assuming unequal variance by comparing different time points with 0-h time point (equivalent to the pre-heat time point for experiments with heat treatments). (**F**) 77 K chlorophyll fluorescence spectra. Each curve is an average of 3 biological replicates and normalized to PSII peak at 686 nm. No significant (ns) differences among different time points without heat.

**Supplemental Figure 15.**
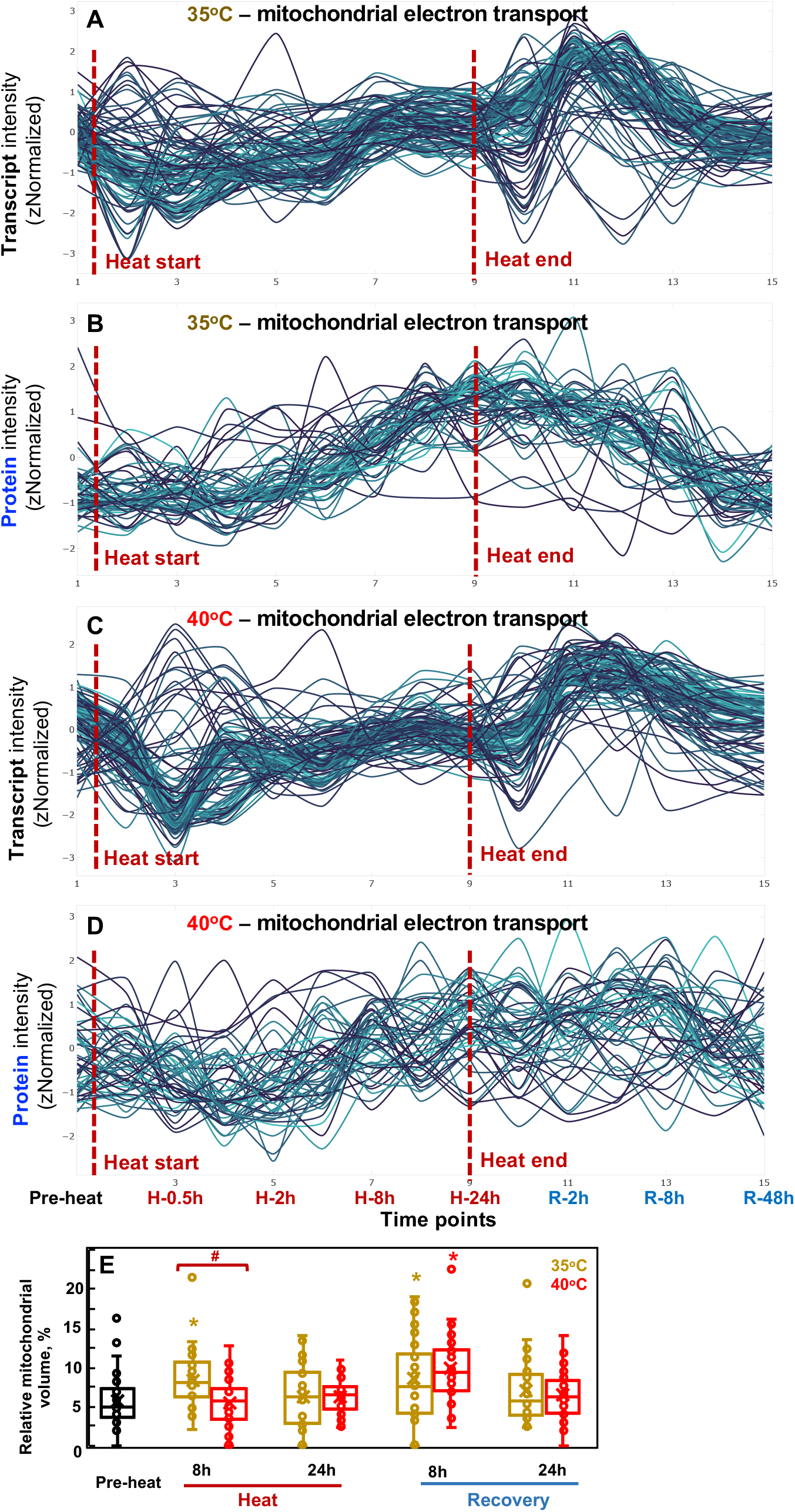
Transcript/protein kinetics and TEM analysis suggested increased and reduced mitochondrial electron transport during 35 and 40°C heat treatments, respectively. Transcript (**A**, **C**) and protein (**B**, **D**) signals related to the MapMan bin mitochondrial electron transport were standardized to z scores (standardized to zero mean and unit variance) and are plotted against equally spaced time point increments. The red dashed lines indicate the start and end time of heat treatment for 35°C (**A**, **B**) and 40°C (**C**, **D**) respectively. Time points are labeled at the bottom. Timepoint 1: pre-heat. Time points 2-9, heat treatment at 35 or 40°C, including reaching high temperature, 0.5, 1, 2, 4, 8, 16, 24 h during heat; time points 10-15, recovery phase after heat treatment, including reaching control temperature, 2, 4, 8, 24, 48 h during recovery. See the interactive figures with gene IDs and annotations in Supplemental Dataset 10, mitochondrial electron transport _ ATP synthesis.html. (**E**) Relative volume fractions of mitochondria were quantified using TEM images and Stereo Analyzer with Kolmogorov–Smirnov test for statistical analysis compared to the pre-heat condition (*, p<0.05, the colors of asterisks match treatment conditions) or between 35 and 40°C at the same time point (#, p<0.05). Each treatment had three biological replicates, total 30 images per treatment.

## Notes

### Competing Interest Statement

The authors have declared no competing interest.

### Summary of Updates

Add Supplemental Figure 1F. Minor revision of the discussion about the up-regulation of cell cycle genes and down-regulation of photosynthetic light reaction genes during the recovery.

