## Supplementary material for "Systems-wide Analysis Revealed Shared and Unique Responses to Moderate and Acute High Temperatures in the Green Alga *Chlamydomonas reinhardtii*": 8supplementary_dataset9_transcript_protein_correlation: PS.photorespiration.html

### Panel order

| Position | Window | Condition | Time points |
| --- | --- | --- | --- |
| top left | HS1 | heat treatment | 0 h - 1 h |
| top center | HS2 | heat treatment | 2 h - 8 h |
| top right | HS3 | heat treatment | 16 h - 24 h |
| bottom left | RE1 | recovery | 0 h - 2 h |
| bottom center | RE2 | recovery | 4 h - 8 h |
| bottom right | RE3 | recovery | 24 h - 48 h |
