## Supplementary figures and images for "Systems-wide Analysis Revealed Shared and Unique Responses to Moderate and Acute High Temperatures in the Green Alga *Chlamydomonas reinhardtii*"

### Fig9_TEM_high_resolution

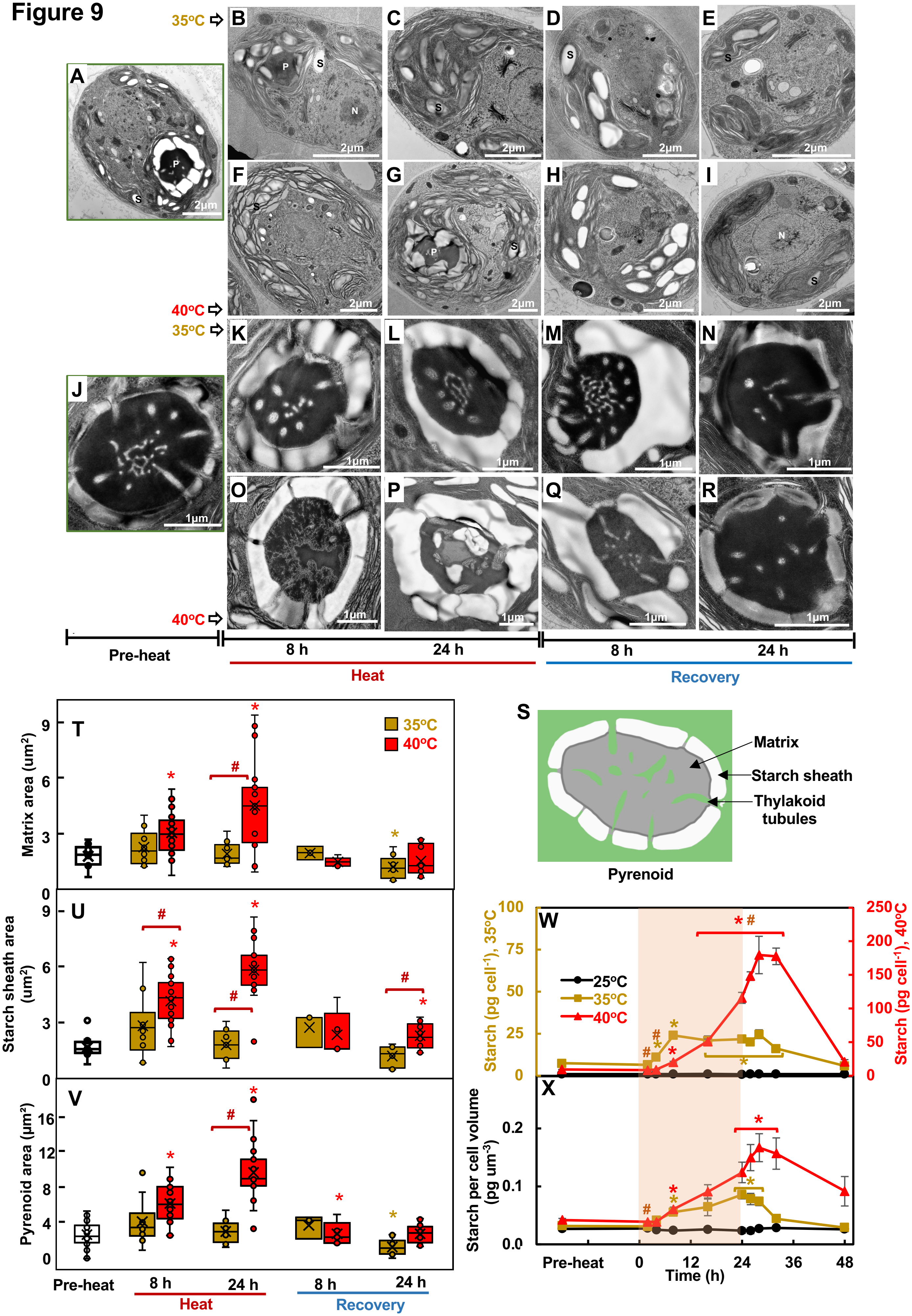
